# The “motive cocktail” in altruistic behaviors

**DOI:** 10.1101/2023.08.20.554026

**Authors:** Xiaoyan Wu, Xiangjuan Ren, Chao Liu, Hang Zhang

## Abstract

Prosocial motives such as social equality and efficiency are key to altruistic behaviors. However, predicting the range of altruistic behaviors in varying contexts and among different individuals proves challenging if we limit ourselves to one or two motives, as most previous studies have done. Here we demonstrate the numerous, interdependent motives in altruistic behaviors and the possibility to disentangle them through behavioral experimental data and computational modeling. In one laboratory experiment (*N*=157) and one pre-registered online replication (*N*=1258) across 100 different situations, we found that both third-party punishment and helping behaviors aligned best with a model of seven socioeconomic motives, referred to as a “motive cocktail”, including two compound motives. For instance, the “inequality discounting” motives imply that individuals, when confronted with costly interventions, behave as if the inequality between others barely exists. The motive cocktail model also provides a unified explanation for a range of phenomena in the literature.

## Introduction

Many people voluntarily provide resources like shelter, food, and healthcare to refugees fleeing war-torn regions, while others advocate sanctioning responsible nations, even at the expense of personal cost. This altruistic behavior, known as third-party punishment (3PP) and helping (3PH), involves sacrificing personal interests to punish transgressors or help victims. Such behaviors have been observed both in laboratory (Fehr & Fischbacher, 2004; Henrich et al., 2006; Jordan et al., 2016) and field studies (Balafoutas et al., 2014, 2014; Singh & Garfield, 2022). What, then, motivates these actions?

According to one line of theories, third-party intervention serves as a strategic means to obtain future rewards, by signaling one’s trustworthiness to potential cooperators (Bénabou & Tirole, 2006; Jordan et al., 2016) or deterring potential transgressors from harming oneself or valued others (Delton & Krasnow, 2017). However, third-party intervention in one-shot, anonymous scenarios (Fehr & Fischbacher, 2004) aligns more with the strong reciprocity theory (Gintis, 2000), where individuals may reward cooperation, punish non-cooperation, or more generally, sanction violations of social norms (Claessens et al., 2024; Kimbrough & Vostroknutov, 2016) even without prospect of personal gain. These two lines of theories are not necessarily conflicting; the motives for sanctioning norm violations can be viewed as internalized external motivations. A widely observed norm in human societies is egalitarian distribution. By quantifying inequality—a violation of this norm—as a loss in a utility maximization framework, Fehr and Schmidt (1999) provide a unified explanation for various social-economic phenomena, including altruistic punishment and helping behaviors (Fehr & Fischbacher, 2004; Stallen et al., 2018; Zhong et al., 2016). Human representation of inequality is further supported by neuroimaging studies (Hsu et al., 2008; Stallen et al., 2018; Tricomi et al., 2010).

The power of this normative framework (Fehr & Fischbacher, 2004), lies in its potential to integrate different motives into one utility measure to address the complexity of human altruistic behaviors. However, its potential is far from thoroughly explored, because most previous studies only focused on one or two motives (other than self-interest) and often contrasted models with distinctive motives (Engelmann & Strobel, 2004; Zhong et al., 2016), as if human behaviors were guided exclusively by one of the alternative motives at each moment. Such practice makes it difficult to unify the knowledge gained from different studies that examine different motives. Furthermore, it limits the power of the normative framework to explain intricate behavioral patterns.

For example, when a victim seeks revenge against the transgressor, a trade-off between self-interest and inequality reduction would predict either no punishment or full punishment to restore equality, depending on whether the impact ratio of the punishment is below or above a certain threshold (see Fig. S1). But people often choose to punish the transgressor without fully restoring equality (Fehr & Fischbacher, 2004), which some researchers explain by resorting to a separate personal tendency called “willingness to punish” (Stallen et al., 2018), a factor not motivated by socio-economic utilities. The hesitation of previous studies to simultaneously test multiple motives may be partly due to limitations in their experimental designs (Engelmann, 2012), where different motives often yield similar predictions (Charness & Rabin, 2002), making them empirically indistinguishable. However, practices from relatively developed modeling-reliant fields such as human decision making (Peterson et al., 2021) and working memory (Huang, 2023; Van Den Berg et al., 2014) suggest that including multiple motives in one model and empirically teasing them apart are both plausible and valuable for advancing our understanding of human altruistic behaviors.

Here we aimed to extend the normative framework of utility maximization to provide a unified explanation for a wider range of phenomena in altruistic behaviors. We constructed a series of computational models that assume altruistic behaviors are driven jointly by multiple socioeconomic motives. These “cocktail motive” models cover a comprehensive set of socioeconomic motives. Five of the motives are based on established theories from the literature, including two variants of self-centered inequality (Fehr & Fischbacher, 2004; Zhong et al., 2016), victim-centered inequality (Zhong et al., 2016), efficiency (Engelmann & Strobel, 2004; Hsu et al., 2008), and reversal preference (Li et al., 2022; Xie et al., 2017). While some of the established socioeconomic motives are qualitatively similar, they lead to different quantitative patterns and can thus be told apart through computational modeling. Furthermore, we also identified two new “compound” motives that are non-linear combinations of more elementary motives.

To separate the effects of different socioeconomic motives, we need an experimental setup that can systematically vary all the motives in the same context. We thus designed a third-party intervention task—the Intervene-or-Watch task (Fig. 1a–b), which enables an unusually rich set of experimental conditions for testing this variety of motives that would otherwise be undistinguishable. On each trial (Fig. 1c–d), participants saw the outcomes from a dictator game, where the dictator (“transgressor”) allocated more to themselves than to the receiver (“victim”, e.g., 88 vs. 12 tokens). As the unaffected third-party, participants decided whether to accept a specific intervention offer (e.g., spending 10 tokens of their own payoff to reduce the transgressor’s payoff by 15 tokens). Each participant completed 300 trials in 100 different conditions that varied in the transgressor-victim inequality as well as the scenario (punishment vs. helping), the cost and the impact-to-cost ratio of the intervention offer.

**Fig. 1.**
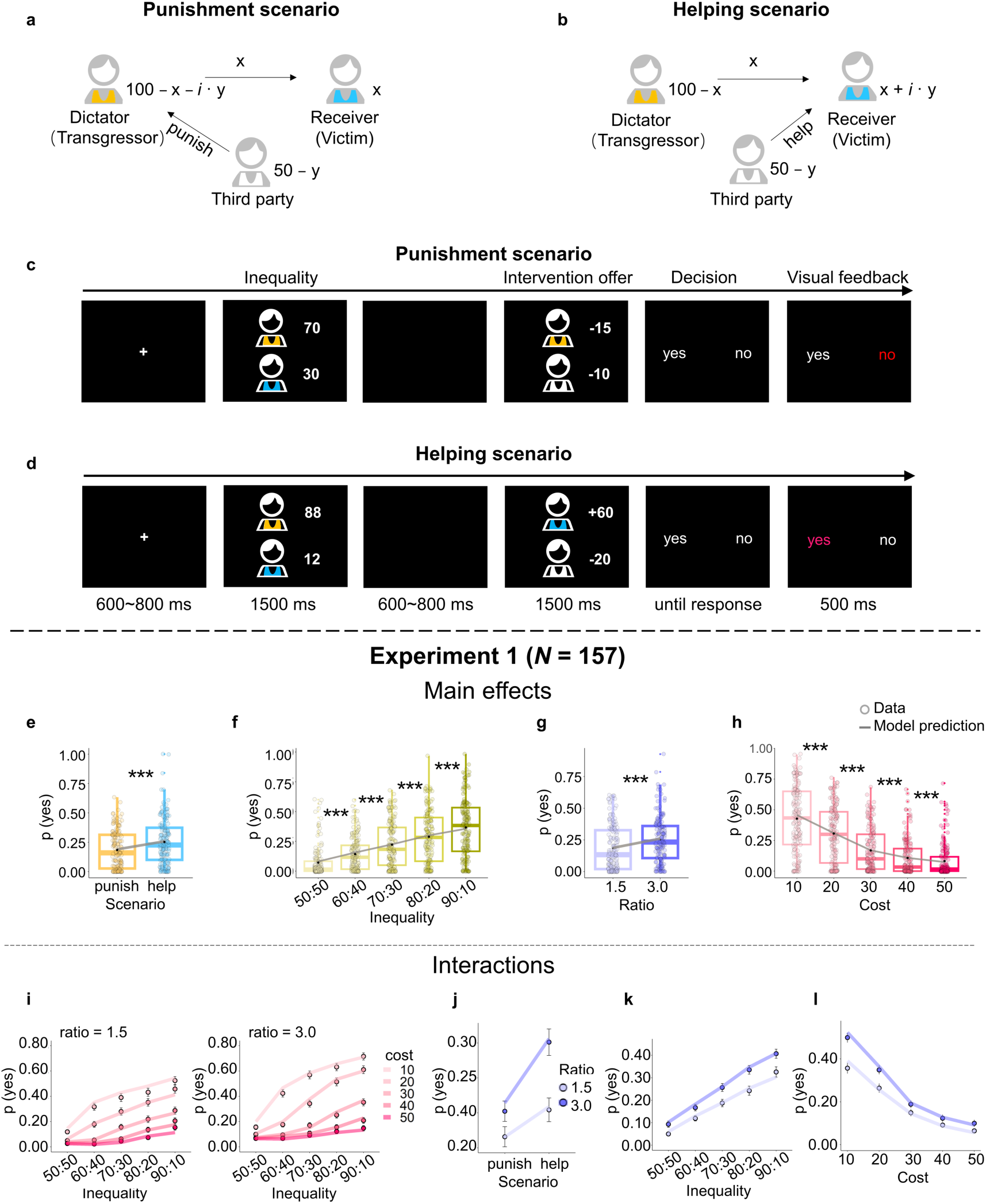
The Intervene-or-Watch task and participants’ behavioral patterns. **a**–**b**, Schema of the Intervene-or-Watch task for the punishment scenario (**a**) and the helping scenario (**b**). **c**–**d**, Time course of a trial for the punishment (**c**) and helping (**d**) scenarios. On each trial, participants first saw the outcome of a dictator game—out of 100 tokens how much the dictator (“transgressor”, cartoon figure in orange shirt) allocated to themselves and to the receiver (“victim”, blue shirt), such as 70 vs. 30 (**c**) or 88 vs. 12 (**d**). As a third party starting with 50 tokens, participants (white shirt) were provided with an intervention offer, such as spending their own 10 tokens to reduce the transgressor’s payoff by 15 tokens (**c**), or spending their own 20 tokens to increase the victim’s payoff by 60 tokens (**d**). Participants’ task was to decide whether to accept the intervention offer (press “yes”) or do nothing (press “no”). **e**–**h**, Main effects of scenario (**e**), transgressor-victim inequality (**f**), impact-to-cost ratio (**g**), and intervention cost (**h**) on the probability of accepting the intervention offer, *p*(yes). Participants had a higher probability to help the victim than to punish the transgressor (**e**), were more willing to intervene when the transgressor-victim inequality was more extreme (**f**), when the impact-to-cost ratio was higher (**g**), and when the intervention cost was lower (**h**). For each box plot, the bottom, middle, and top lines of the box respectively indicate the first quartile, the median, and the third quartile; the whiskers represent 1.5 times the interquartile range (IQR), which is the distance between the third quartile (Q3) and the first quartile (Q1). Data points beyond 1.5 times the IQR from the upper and lower quartiles are considered outliers and are represented by the color points. Each filled circle denotes one participant. The *** denotes *p* < 0.001 for the difference between adjacent conditions from the post hoc comparison, and *p*-values were corrected using the Bonferroni method. The black dot inside each box denotes the group mean. The gray line superimposed on the boxes denotes the prediction of the best-fitting model (i.e., the seven-motive “motive cocktail” model, described later). **i**–**l**, Interaction effects on *p*(yes). The inequality × cost × ratio three-way interaction (**i**) shows that participants’ probability of intervention increased more slowly with the transgressor-victim inequality when the intervention cost was higher, with this modulation effect more pronounced under a higher impact-to-cost ratio. The two-way interactions involving the impact ratio show that under a higher ratio, the preference for helping over punishment was stronger (**j**), and the probability of intervention changed more dramatically with inequality (**k**) and with cost (**l**). Each dot denotes the mean across participants. Error bars denote SEM. As in **e**–**h**, the lines denote the predictions of the best-fitting model.

We performed one laboratory experiment (*N*=157) and a pre-registered online experiment (*N*=1258), with all major findings of the former replicated in the latter. A three-way interaction of inequality × cost × impact ratio found in participants’ intervention decisions suggests utility calculations that go beyond linear combinations of different motives. Indeed, participants’ behavioral patterns were best fit by a “motive cocktail” model whose utility calculation involves seven socioeconomic motives, including two compound motives. We called the compound motives “inequality discounting”, which refers to people’s tendency to behave as if they are underestimating the inequality between others as the intervention cost increases. Individuals’ cocktail motives fall into three groups: *justice warriors* who have a strong intention to intervene whenever there is inequality, *pragmatic helpers* who are sensitive to the impact of their intervention to help the victim, and *rational moralists* who seek to achieve an acceptable standard of morality at the lowest cost to self-interest. Our model provides a unified explanation not only for 3PP and 3PH, but also for a wider range of phenomena in the literature.

## Results

On each trial of our Intervene-or-Watch task (Fig. 1c–d), participants saw the results of a dictator game, where an anonymous dictator (“transgressor”) allocated more amounts to themselves than to an anonymous receiver (“victim”), such as 88 vs. 12 tokens. As the third-party, participants received 50 tokens in each trial and were offered an opportunity to intervene, such as spending 10 tokens (intervention cost) to reduce the transgressor’s payoff by 15 tokens (impact ratio = 15/10 = 1.5). The task was to decide whether to accept this intervention offer to intervene or to keep all 50 tokens to themselves. Each trial was either in a punishment scenario (as the example above, Fig. 1a & 1c) or in a helping scenario (to increase the victim’s payoff, Fig. 1b & 1d). Across trials, we varied the inequality between the transgressor and the victim (50:50, 60:40, 70:30, 80:20, or 90:10, with ±2 jitters), the intervention cost (10, 20, 30, 40, or 50), and the impact ratio (1.5 or 3.0). Each participant completed 300 trials (5 inequality levels × 5 cost levels × 2 impact ratios × 2 scenarios × 3 repetitions) of intervention decisions.

### Behavioral patterns in third-party punishment and helping

In Experiment 1, there were 157 participants (all students). We first performed a generalized linear mixed model analysis (GLMM1, see Table S1) on participants’ decisions (to intervene or not) to assess the effects of each independent variable and their interactions.

#### Preference for helping over punishment

Consistent with most previous studies, participants had a higher probability to help the victim (M = 0.25) than to punish the transgressor (M = 0.18, *b* of scenario = –1.22, 95% CI [–1.64, –0.80], *p* < 0.001; Figure 1e).

#### Inequality aversion and rationality

As one would expect from inequality aversion, participants were more willing to intervene when the transgressor-victim inequality was more extreme (*b* = 1.61, 95% CI [1.40, 1.81], *p* < 0.001; Figure 1f) and when the impact-to-cost ratio was higher, that is, when the same cost yielded a higher reduction in inequality (*b* = 0.82, 95% CI [0.62, 1.01], *p* < 0.001; Figure 1g). Meanwhile, participants were also rational decision makers who cared about their own interest, being less willing to intervene under a higher cost of intervention (*b* = –2.12, 95% CI [–2.37, –1.86], *p* < 0.001; Fig. 1h).

#### Interaction effects

Thanks to our factorial experimental design with 4 dimensions and 100 conditions, we also identified three two-way and one three-way interaction effects that were seldom documented before. Under a higher impact-to-cost ratio, the preference for helping over punishment was stronger (scenario × ratio interaction: *b* = –0.39, 95% CI [–0.47, –0.30], *p* < 0.001; Fig. 1j), and the probability of intervention changed more dramatically with the transgressor-victim inequality (inequality × ratio interaction: *b* = –0.08, 95% CI [–0.14, –0.02], *p* = 0.017; Fig. 1k) and with cost (cost × ratio interaction: *b* = –0.08, 95% CI [–0.14, –0.02], *p* = 0.015; Fig. 1l). According to the three-way interaction of inequality × cost × ratio (*b* = –0.21, 95% CI [–0.27, –0.15], *p* < 0.001), a higher ratio also led to a stronger modulation of the intervention cost to participants’ sensitivity to inequality (Fig. 1i).

In sum, the behavioral patterns in our Intervene-or-Watch task not only agree with classic effects about altruistic punishment and helping, but also reveal intriguing interaction effects.

### Seven socioeconomic motives and their hypothetical effects

What socioeconomic motives may have driven the observed 3PP and 3PH behaviors? Besides self-interest (the core of classical economic models), we considered five classes of computationally well-defined socioeconomic motives (Fig. 2a), which expand into seven motive terms in utility calculation (see Table S2 and Table S3 for examples in fictitious characters and real-life scenarios). Five of these motives are adapted from the literature, including three variants of inequality aversion (Fehr & Fischbacher, 2004; Zhong et al., 2016), efficiency (Engelmann & Strobel, 2004; Hsu et al., 2008), and reversal preference (Li et al., 2022; Xie et al., 2017). The remaining two motives under the class of “*inequality discounting*” are newly defined here to capture the interaction between self-interest and inequality aversion. They are partly motivated by the observed interaction effect that under higher intervention cost, participants’ probability of intervention not only was lower, but also increased more slowly with the transgressor-victim inequality (Fig. 1i). As unfolded below, each motive affects the utility gain from intervention relative to non-intervention (thus the tendency to intervene) in a different way (Fig. 2b).

**Fig. 2.**
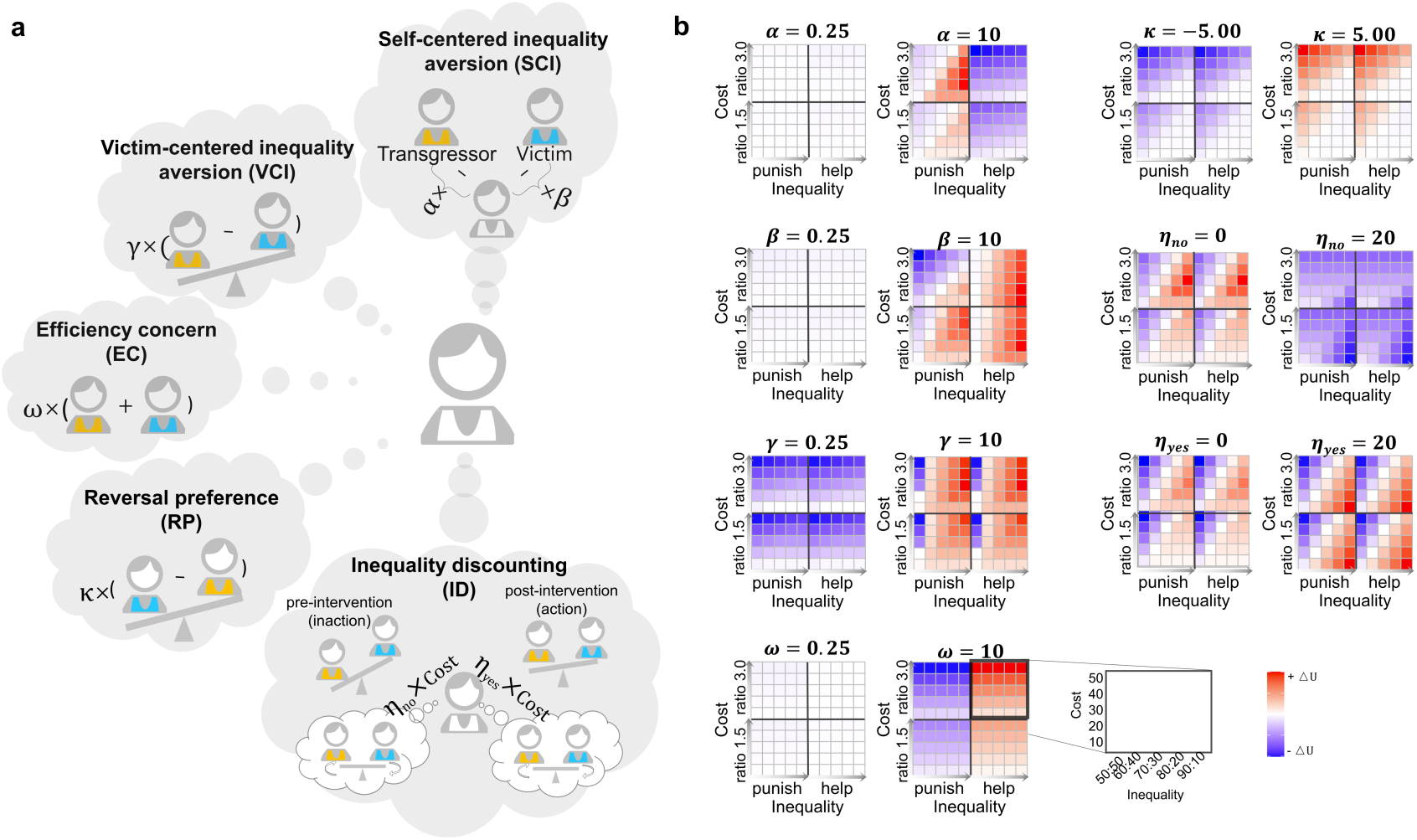
The seven socioeconomic motives and their hypothetical effects on the third-party’s utility gain to intervene relative to not to intervene. **a**, Five classes of computationally well-defined socioeconomic motives that expand into seven motive terms in utility calculation. The parameters *⍺* and *β* respectively control the motives of disadvantageous (self < other) and advantageous (self > other) inequality aversion. Note that the illustration for SCI in the figure (disadvantageous SCI between self and transgressor, advantageous SCI between self and victim) may not apply to the inequality after intervention, where the direction of inequality between the participant and the other two parties might be reversed. Which type of SCI applies only depends on whether self > other or self < other, irrespectively of the other is transgressor or victim. The parameter *γ* controls the motive of victim-centered disadvantageous (victim < transgressor) inequality aversion. The parameter *κ* (could be positive or negative) controls the direction and strength of the reversal preference motive (victim > transgressor after intervention). The parameter *ω* controls the strength of efficiency concern (maximizing the total payoff of others). The parameters *η*_*no*_ and *η*_*yes*_ respectively control the motives of inaction and action inequality discounting (attenuated perception of inequality under higher intervention cost). **b**, How the strength of each motive would influence the ΔU (utility of choosing yes – utility of choosing no) of the third-party intervention decision. The effect of each motive is illustrated by a pair of panels, with the small and large parameters controlling the motive’s magnitude differently. For simplicity, when examining the effects of one single parameter, we set all other parameters to zero. The parameters *η*_*no*_ and *η*_*yes*_ are exceptions, where the parameter *γ* is set to 1, because their utility terms are multiplied by *γ*. Each heatmap contains four submaps, horizontally divided by scenario (left for punishment, right for helping) and vertically by impact ratio (bottom for 1.5, top for 3.0). The x-axis of each submap denotes the severity of inequality, running from near equality (left: 50:50) to extreme inequality (right: 90:10). The y-axis of each submap denotes the intervention cost, from low cost (bottom: 10) to high cost (top: 50). Colors code ΔU, where more reddish (more positive ΔU) corresponds to a stronger preference for choosing yes while more bluish (more negative ΔU) a stronger preference for choosing no. For illustration purposes, the values of ΔU were scaled separately for each column and separately for positive and negative values. Each motive shows a distinct influence on ΔU and would thus lead to distinguishable effects on third-party intervention decision behaviors.

Self-centered inequality refers to the payoff difference between self and others (Fehr & Fischbacher, 2004). Depending on whether it favors self or other, it can be further divided into disadvantageous inequality (self < other) and advantageous inequality (self > other), which lead to two variants of inequality aversion motives that are respectively controlled by parameters *⍺* and *β*. *⍺* implies a stronger aversion to receiving lower payoff than others (e.g., self 50 vs. transgressor 88). *β* implies a stronger aversion to receiving higher payoff than others (e.g., self 50 vs. victim 12). In our experimental setting, before intervention, the participant always had lower payoff than the transgressor but higher payoff than the victim. As the result, *⍺*, participants would be more motivated to penalize the transgressor to reduce their disadvantageous inequality, but less motivated to help the victim because helping would increase their disadvantageous inequality with the transgressor and even create a disadvantageous inequality with the victim (Fig. 2b, row 1 left pair). In contrast, the larger *β*, participants would be more motivated to intervene in both the punishment and helping scenarios, unless the larger punishment leads to an undesirable advantageous inequality over the transgressor (Fig. 2b, row 2 left pair).

Victim-centered inequality refers to the payoff difference between the transgressor and the victim (Zhong et al., 2016). This variant of inequality aversion implies that participants have a distaste for the higher payoff of the transgressor over the victim. Participants with larger *γ* would intervene more in almost all punishment and helping scenarios (Fig. 2b, row 3 left pair), unless the victim-centered disadvantageous inequality is too small (e.g., transgressor 51 vs. victim 49) to compensate for the cost of intervention.

Efficiency concern is a motive frequently used for modeling economic games (Engelmann & Strobel, 2004; Hsu et al., 2008) but seldom for 3PP or 3PH before assumes that people care about the overall welfare of others, that is, the sum of the transgressor’s and the victim’s payoffs in our case. Participants with larger *ω* would be more likely to help the victim to increase the overall welfare, but less likely to penalize the transgressor to avoid reducing the overall welfare, regardless of the inequality between others (Fig. 2b, row 4 left pair).

Reversal preference refers to the motive that participants intend to reverse the payoff difference between the transgressor and the victim, rewarded by their payoff difference in the opposite direction (i.e., after intervention the victim would be more well-off than the transgressor). The parameter *κ* controlling reversal preference is allowed to be either positive or negative, respectively implying a willingness or reluctance to reverse the economic status of others, thus making the term a generalized form of rank reversal aversion (Li et al., 2022; Xie et al., 2017). Individuals with more positive *κ* would be more willing to punish or help when the impact (i.e., cost × ratio) is large enough (relative to the inequality) to yield a rank reversal between the transgressor and the victim (Fig. 2b, row 1 right pair).

Inequality discounting refers to people’s tendency to behave as if they are underestimating the inequality between others as the intervention cost increases. We defined two types of inequality discounting motives: *inaction inequality discounting* (controlled by *η*_*no*_) and *action inequality discounting* (controlled by *η*_*yes*_), which represent a diminished awareness of inequality when choosing not to intervene and when opting to intervene, respectively. Inequality discounting motives are “compounds” that are not just the lack of motivation to reduce inequality as characterized by smaller *γ* (victim-centered inequality aversion), but capture the modulation of self-interest on victim-centered inequality aversion in both directions. Participants with larger *η*_*no*_ would be less likely to intervene, which differs from that of smaller *γ* in that it may cause no intervention even when the transgressor-victim inequality is high (Fig. 2b, row 2 right pair). Conversely, participants with larger *η*_*yes*_ would be more likely to intervene, as if they believe inequality is always minimized following a costly intervention (Fig. 2b, row 3 right pair).

Many of these motives would remain unidentifiable in a task involving only two parties, testing exclusively either punishment or helping scenarios, or lacking variation in cost or impact ratio. However, in our Intervene-or-Watch task, the seven motives forecast unique effects on intervention decisions, thus making them distinguishable in behavioral data. Subsequent modeling analysis validated the discernibility of each parameter, even under simultaneous modeling (see Methods and Fig. S2).

### The “motive cocktail” model with all seven motives best predicts human behaviors

We assessed the seven socioeconomic motives’ contribution to altruistic behavior by incrementally incorporating them into utility calculations, creating a series of increasingly complex computational models. The introduction of different motives follows a descending order concerning how central and established a specific motive is in the literature of third-party punishment and helping. We then compared these models’ predictive power for the behavioral patterns observed in Experiment 1. Such solution-oriented approach is similar to the idea of “quasi-comprehensive exploration” introduced by a recent study on spatial working memory (Huang, 2023). Starting from a baseline coin-flipping model, which intervened at a fixed probability, and a self-interest (SI) model, we introduced five motive classes as utility terms in the following order: self-centered inequality (SCI), victim-centered inequality (VCI), efficiency concern (EC), reversal preference (RP), and inequality discounting (ID). This process yielded seven different models (see Methods) with different predictions (Fig. 3). We used maximum likelihood estimation to fit each model to individual participants’ decisions, and the corrected Akaike Information Criterion (AICc, Hurvich & Tsai, 1989) to evaluate each model’s relative goodness-of-fit, accounting for complexity. We also computed the protected exceedance probability (PEP, Rigoux et al., 2014) to provide a group-level measure of each model’s potential superiority.

**Fig. 3.**
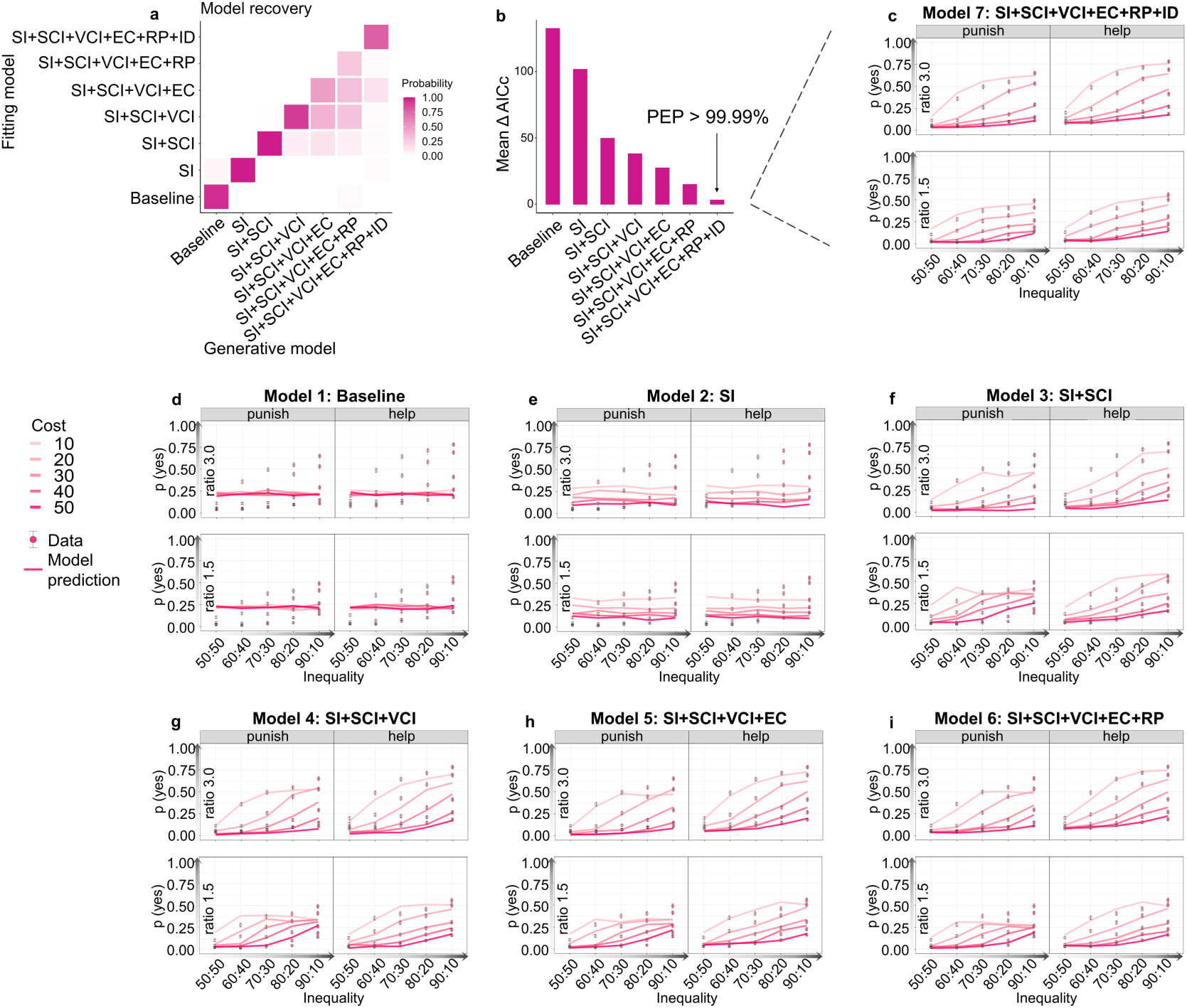
The “motive cocktail” model with all seven socioeconomic motives best predicts participants’ behavioral patterns. **a**, Model recovery analysis. Each model was used to generate 100 synthetic datasets, for each of which model fitting and comparison were performed. Each column is for one generative model. Each row is for one fitting model. The color in each cell codes the probability that the synthetic datasets from the generative model in the column are best fit by the fitting model in the row, with darker color indicating higher probability. **b**, Model comparison results. For each participant, the model with the lowest AICc was used as a reference to compute ΔAICc by subtracting it from the AICc of the other models (ΔAICc = AICc – AICc_lowest_). Lower ΔAIC indicates better fit. Protected exceedance probability (PEP) is a group-level measure of the likelihood of each model’s superiority over others. The name of a model (e.g., SI+SCI) conveys the motives included in its utility calculation. SI: self-interest. SCI: self-centered inequality. VCI: victim-centered inequality. EC: efficiency concern. RP: reversal preference. ID: inequality discounting. Both ΔAICc and PEP suggest that the full “motive cocktail” model best fits participants’ decision behaviors. Panels **c–i** show data versus model predictions separately for the seven models compared in **b**. The title of each panel indicates its model name. The probability of intervention, *p*(yes), is plotted against the inequality (from 50:50 to 90:10). Different colors code different levels of intervention cost (from 10 to 50, darker color for higher cost). Each sub-panel corresponds to one scenario and impact ratio condition. The dots and error bars respectively denote the mean and SEM across participants. The solid lines denote the predictions of the models.

The full “motive cocktail” model that includes all the motives best predicted participants’ decisions (lowest AICc, PEP > 99.99% among the seven models). A model recovery analysis (see Methods) further confirmed that the superiority of the full model was real and could not attribute to model misidentification: Among the 700 synthetic datasets generated by the six alternative models, none was misidentified as the full model (Fig. 3a). Integrating each motive class (SI, SCI, VCI, EC, RP, & ID) into our models led to considerable improvements in their fits (as indicated by lower AICc values in Fig. 3b).

The full model closely mirrored changes in participants’ intervention probabilities across the 100 experimental conditions (Fig. 3c), successfully predicting the main and interaction effects of different variables (lines in Fig. 1e–1l). In contrast, alternative models failed to replicate certain patterns within the data (Fig. 3d–3i). A supplementary analysis that compared more model variants further demonstrated the necessity of the inequality discounting assumption (the interaction items) in the full model as well as the nonlinear modulation of self-interest on the VCI (Fig. S3) in fitting the behavioral data. The inequality discounting term follows the form of a sigmoid function (Fig. S4b), which has the desired mathematical property of ensuring its value being between 0 and 1. To conclude, participants’ third-party intervention decisions were jointly driven by self-interest and the seven socioeconomic motives, including the two inequality discounting terms.

### Individual differences: justice warriors, pragmatic helpers, and rational moralists

Our Intervene-or-Watch task, with its 100 factorially-designed conditions, yielded a rich, multifaceted profile that captured not only the collective behavioral tendencies but also the nuanced 3PP and 3PH behaviors of individual participants. A clustering analysis of the behavioral patterns of the 157 participants revealed that they were best summarized by three distinct clusters (see Methods and Fig. 4a-b). Among them, the “justice warriors” (35% of participants) had an overall high probability to intervene, especially when the transgressor-victim inequality was high, and the cost was relatively low (Fig. 4j). The “pragmatic helpers” (18%) also had a high probability to intervene, but were insensitive to inequality or cost, and preferred helping over punishment (Fig. 4k). The “rational moralists” (47%) barely intervened unless their intervention cost was minimal (Fig. 4l). The full “motive cocktail” model accurately predicted not only the average behavior (Fig. 4i) but also the behavioral patterns specific to each individual cluster (Fig. 4j–l).

**Fig. 4.**
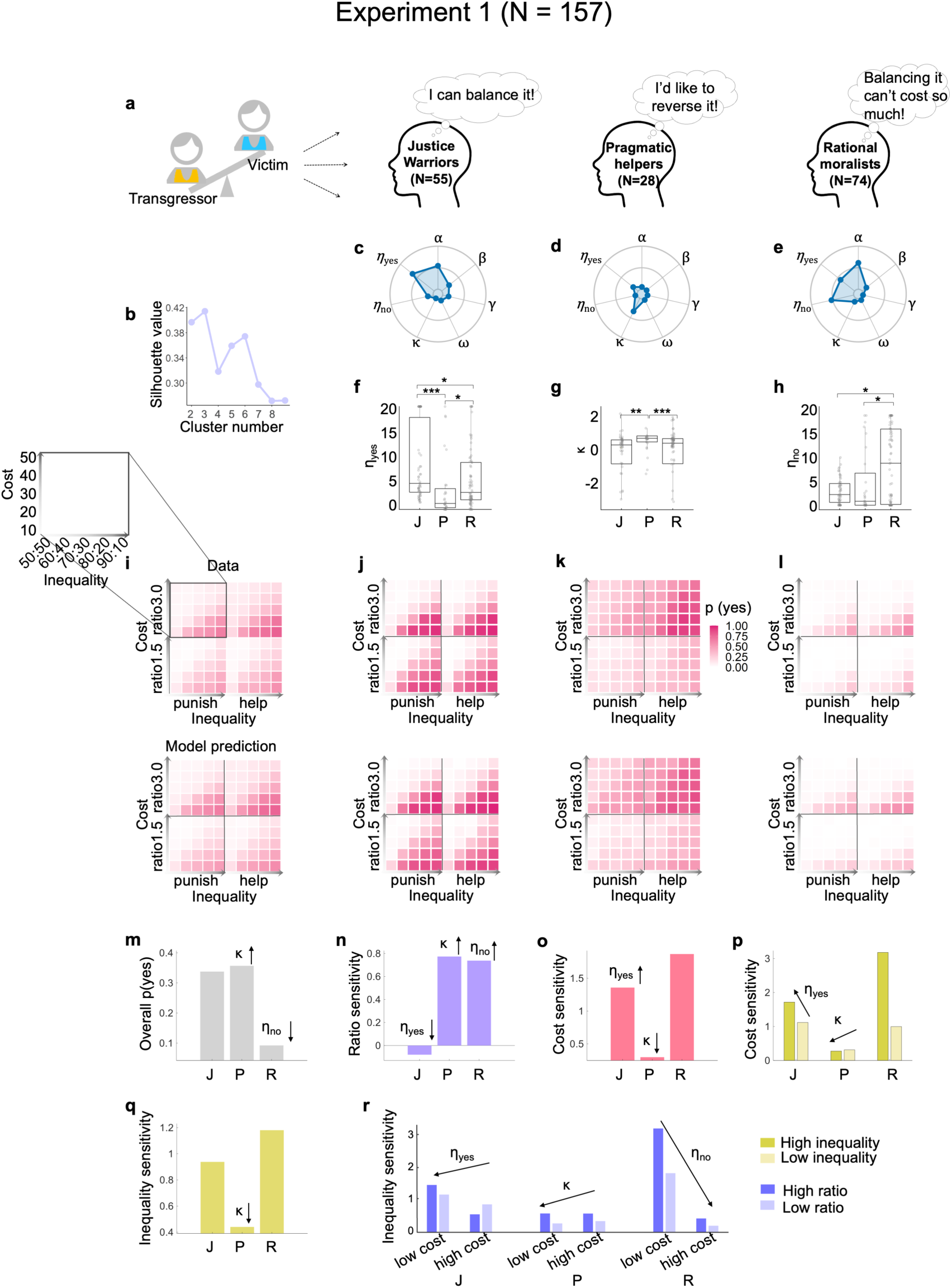
Three types of 3PP and 3PH behaviors: justice warriors, pragmatic helpers, and rational moralists. **a**, Illustration of the three types of behavioral patterns. **b**, The clustering performance of behavioral patterns was best for three clusters. To identify the distinct types of behaviors in the 157 participants of Experiment 1, we used *k*-means to classify participants’ behavioral patterns into clusters and used the silhouette value as the metric for the goodness of clustering, where a higher value indicates a larger ratio of the between-cluster to within-cluster distance. **c**–**e,** The median value of the motive parameters for each cluster. The outer contour of the spider plot indicates the highest median value for a specific parameter summarized across all clusters. **f**–**h,** The values of the action inequality discounting parameter *η*_*yes*_ (**f**), reversal preference parameter *κ* (**g**) and inaction inequality discounting parameter *η*_*no*_ (**h**) compared across the three clusters of participants. J: justice warriors. P: pragmatic helpers. R: rational moralists. The highest values of *η*_*yes*_, *κ* and *η*_*no*_ respectively occurred at justice warriors (J), pragmatic helpers (P), and rational moralists (R). For each box plot, the bottom, middle, and top lines of the box respectively indicate the first quartile, the median, and the third quartile; the whiskers represent 1.5 times the IQR, which is the distance between the Q3 and the Q1. **i–l**, The intervention probability *p*(yes) in the 100 different conditions for all participants (**i**) and for each cluster (**j–l**), with data (top panels) contrasted with the prediction of the full motive cocktail model (bottom panels). Similar to Fig. 2b, each heatmap contains four submaps, horizontally divided by scenario (left for punishment, right for helping) and vertically by impact ratio (bottom for 1.5, top for 3.0). The x-axis of each submap denotes the severity of inequality, running from near equality (left: 50:50) to extreme inequality (right: 90:10). The y-axis of each submap denotes the intervention cost, from low cost (bottom: 10) to high cost (top: 50). Colors code *p*(yes), where darker colors correspond to higher probabilities to choose intervention. **m–r**, How the three parameters (*η*_*yes*_, *κ* and *η*_*no*_) may contribute to the behavioral differences across the three clusters of participants. Each panel is for one main or interaction effect that was detected across individuals (as in Fig. 1), with the bar height denoting the corresponding effect size in each cluster. The arrow accompanying a parameter in a specific panel indicates the significant correlation between parameters and behavioral measures, and how the parameter modulates the plotted behavioral measure (the orientation of the arrow) in the group level (Fig. S6). For example, panel **m** shows that higher *κ* was associated with higher overall *p*(yes) and higher *η*_*no*_ with lower overall *p*(yes), which coincide with the observed high *p*(yes) in pragmatic helpers (**k**) and low *p*(yes) in rational moralists (**l**). The sensitivity for a specific variable was calculated as the normalized intervention probability difference between the high and low manipulated conditions. Specifically, “low inequality” refers to trials with inequality levels of 60:40 and 50:50, whereas “high inequality” refers to inequality levels of 90:10, 80:20, and 70:30. “Low cost” and “high cost” respectively refer to trials with cost ≤ 20 and cost > 20. “Low ratio” and “high ratio” respectively refer to trials with impact ratio of 1.5 and 3.

These dramatic individual differences were associated with different combinations of motive parameters (Fig. 4c–e). Kruskal-Wallis tests and the following post-hoc tests revealed significant differences across the three clusters for three out of the seven motive parameters (Fig. 4f–h & Fig. S5): action inequality discounting *η*_*yes*_ (*H*(2) = 22.18, *p* < 0.001), reversal preference κ (*H*(2) = 15.57, *p* < 0.001) and inaction inequality discounting *η*_*no*_ (*H*(2) = 9.71, *p* = 0.008). The highest values of *η*_*yes*_, κ and *η*_*no*_ respectively occurred at justice warriors, pragmatic helpers, and rational moralists. To unravel the relationship of these parameters to the observed individual differences, we carried out a series of correlation analyses between individuals’ parameter value and their sensitivity to different variables in the group level (multiple comparisons corrected for each parameter using *FDR*; see Fig. S6), where participants’ sensitivity to a variable was defined as the normalized intervention probability difference after the corresponding variable was dichotomized. The observed behavioral differences across clusters coincide with the correlational effects of these parameters (Fig. 4m–r) and agreed with the insights we obtained through simulation (Fig. 2). For example, higher *η*_*yes*_ implies increased tendency to perceive one’s action as effective in reducing inequality, irrespective of the actual impact, when the intervention cost is high. Indeed, individuals with higher *η*_*yes*_ were less sensitive to the impact ratio. Justice warriors, those who had the highest *η*_*yes*_ among the three clusters, were least sensitive to the impact ratio (Fig. 4n).

### Replication in a pre-registered, large-scale online experiment

To test whether our findings can be generalized to a large population with different cultural backgrounds, we performed a pre-registered, large-scale online experiment using the same experimental procedures, with 1258 participants (all students, sample size pre-determined based on a model-based power analysis, Fig. S7) from over 60 countries (or regions, see Table S4). All major statistical and modeling findings of Experiment 1 were replicated in Experiment 2 (Fig. 5; see Supplement for the GLMM results).

**Fig. 5.**
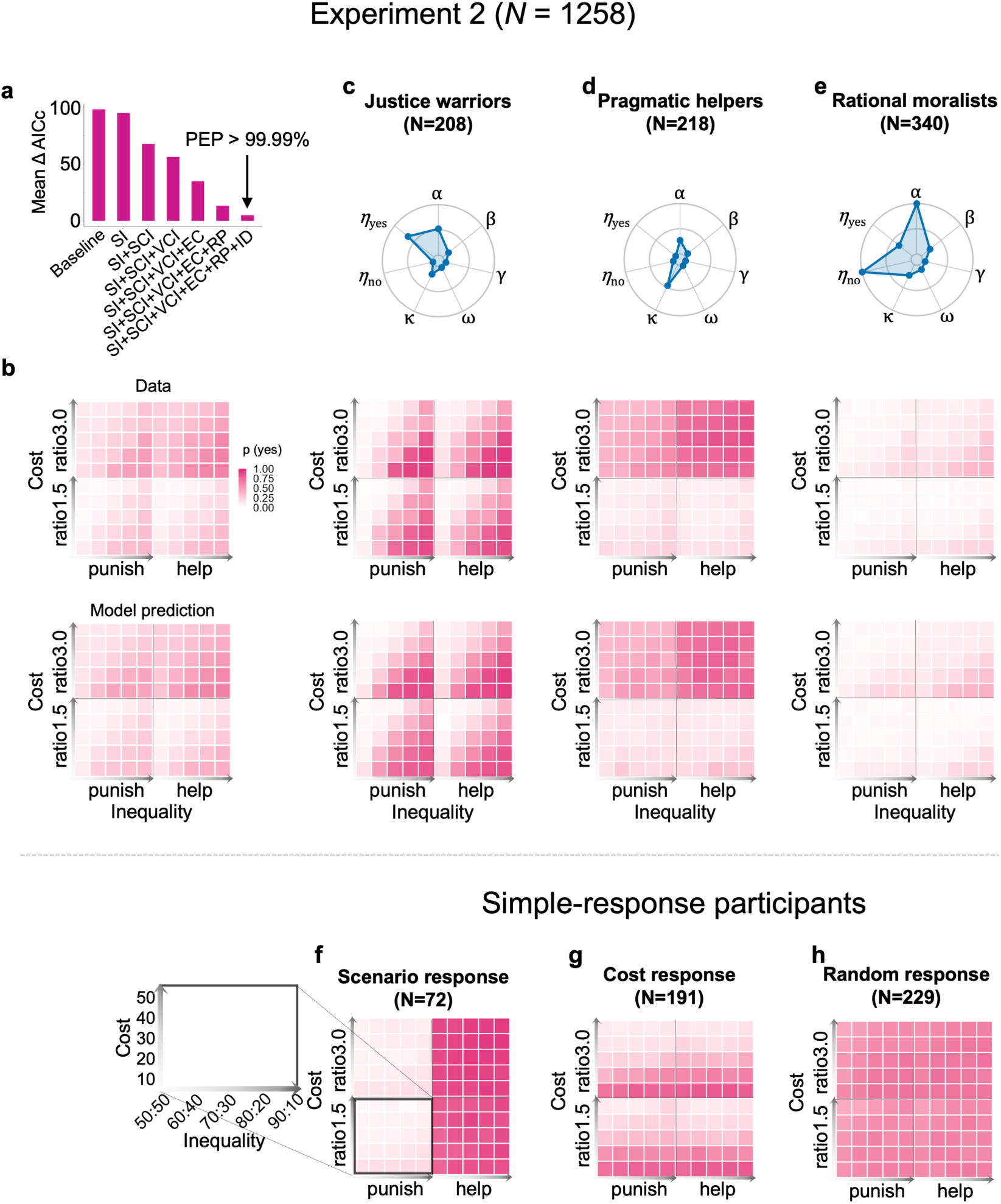
All major findings were replicated in the pre-registered, large-scale online Experiment 2. **a**, Model comparison results. As in Experiment 1, the full motive cocktail model best fit participants’ decision behaviors, as indicated by the lowest ΔAICc and a PEP over 99.9%. **b**, Data versus model prediction. As in Experiment 1, the full model can accurately predict not only participants’ average behaviors, but also that of individual clusters. **c–e**, The median value of the motive parameters for the first three clusters. These three clusters had similar behavioral patterns and parameter combinations to those of the justice warriors, pragmatic helpers, and rational moralists identified in Experiment 1. **f–h**, Data of the three clusters newly observed in Experiment 2. These three clusters were best fit by a simple-response model (Model 9) instead of by the motive cocktail model. **f,** The scenario response cluster, where participants varied their choices only with the scenario, consistently choosing “yes” for the helping scenario but “no” for the punishment scenario. **g,** The cost response cluster, where participants varied their choices only with the cost of intervention. **h,** The random response cluster, where participants seemed to choose randomly, without responding to any variables. These patterns are clues of low effort or engaging participation, which is more frequent among online participants. Conventions follow Fig. 4.

As in Experiment 1, the full motive cocktail model outperformed the other models and accurately captured the behavioral patterns in Experiment 2 (Fig. 6a–b, see Fig. S8 for model recovery analysis). The behavioral patterns of the 1258 participants were best captured by six clusters (Fig. S9), in which the first three clusters agreed with those in Experiment 1—justice warriors (16.60%, Fig. 6c), pragmatic helpers (17.30%, Fig. 6d), and rational moralists (27.00%, Fig. 6e). As in Experiment 1, each of these three clusters were best fit by the full motive cocktail model (or its derivatives; Fig. S9a). The remaining 3 clusters of participants (39.10%, Fig. 5f-h) seemed to respond to one single stimulus dimension (e.g., always help but seldom punish) or even purely randomly, whose choice behaviors were best described by a simple-response model that linearly combines different independent variables (see Methods and Fig. S9b). These choice patterns likely resulted from these participants’ less engaging participation (lower attention check accuracy than participants in the first three clusters: *t*(1256) = –9.78, *p* < 0.001), which is more common in online settings, rather than representing real-world behavioral patterns.

**Fig. 6.**
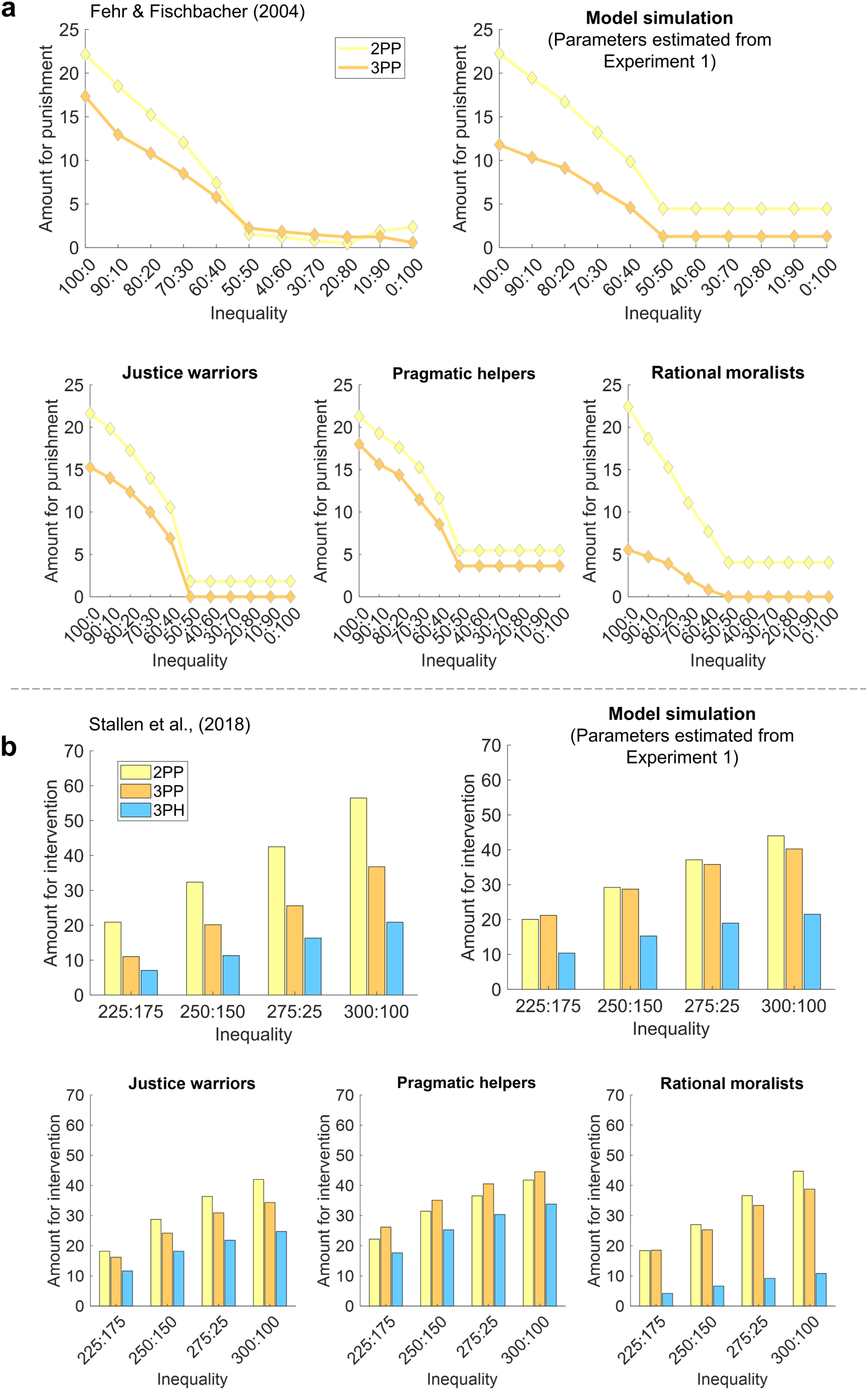
The motive cocktail model successfully reproduces a broader range of phenomena. We used the full motive cocktail model estimated from the Intervene-or-Watch task (3PP and 3PH) to simulate the second-party punishment (2PP) as well as the 3PP and 3PH behaviors in previous publications. In each sub-plot (**a** or **b**), the top-left panel is the data; the upper-right and three lower panels are model simulations respectively based on the estimated parameters of all participants and the three clusters of our Experiment 1. **a,** Reproduction of the 2PP and 3PP behaviors in Figure 5 of Fehr & Fischbacher. The amount participants would use to punish the allocator in a dictator game is plotted as a function of the level of inequality favoring the allocator. In simulating 2PP behaviors, participants—as the second party (the receiver)—were treated as a third party who had all the motives of third parties except for efficiency concern. Our model simulation (with no free parameters) reproduced two effects in the data: (1) the amount participants use for punishment decreases almost linearly with the decrease of inequality when the inequality favors the allocator and is nearly zero when the inequality favors the receiver, and (2) 2PP is larger than 3PP. The simulation based on justice warriors’ parameters best matched the data. **b,** Reproduction of the 2PP, 3PP, and 3PH behaviors in Stallen et al. (2018). The amount participants would use to intervene is plotted as a function of the level of inequality. The task scenario of Stallen et al. (2018) differed from that of Fehr & Fischbacher (2004) in that the first party robs from the second party, causing a more severe violation of social norms. In this case we assume the efficiency concern is excluded from the motive cocktail for all intervention behaviors, which leads to larger amounts for punishment than helping. As in **a**, the simulation based on justice warriors’ parameters best matches the data.

Upon completion of the experiment, participants were asked to fill out personality questionnaires that assessed their prosocial inclinations in everyday life, including the social value orientation scale (SVO, Murphy et al., 2011) to measure selfishness, the interpersonal reactivity index (IRI) (Davis, 1983) for empathy concern. We computed Pearson’s correlation coefficients (*r*) between each participant’s model parameters (from the motive cocktail model) and the participant’s personality measures (Fig. S10 and Fig. S11). In both Experiments 1 and 2, we found that stronger self-centered disadvantageous inequality aversion (*⍺*) or inaction inequality discounting (*η*_*no*_) was associated with more selfishness. When one of these two parameters was controlled, the correlations between *η*_*no*_ and selfishness (Experiment 1: ρ = –0.22, *p* = 0.006; Experiment 2: ρ = –0.16, *p* < 0.001) was still significant, but the correlation between *⍺* and selfishness was significant only in Experiment 2 (Experiment 1: ρ = –0.11, *p* = 0.16; Experiment 2: ρ = –0.12, *p* < 0.001). We also found that both inaction inequality discounting (*η*_*no*_) and action inequality discounting (*η*_*yes*_) was associated with empathy in the opposite direction. When one of these two parameters was controlled, the correlation between *η*_*no*_ and empathy was still significant in both experiments (Experiment 1: ρ = –0.25, *p* = 0.002; Experiment 2: ρ = – 0.12, *p* < 0.001), the correlation between *η*_*yes*_ and empathy was significant only in Experiment 2 (Experiment 1: ρ = 0.12, *p* = 0.13; Experiment 2: ρ = 0.12, *p* < 0.001).

Before the main experiments, we recorded the amounts participants allocated to their receiver in a dictator game. Kruskal-Wallis tests revealed significant differences across the three clusters for both Experiment 1 (*H*(2) = 14.56, *p* < 0.001) and Experiment 2 (*H*(2) = 46.72, *p* < 0.001). In both experiments, rational moralists allocated least to their receiver (see Fig. S12 for post-hoc tests). We also found significant differences between the three clusters of participants in selfishness (Kruskal-Wallis tests, Experiment 1: *H*(2) = 11.70, *p* = 0.003; Experiment 2: *H*(2) = 74.02, *p* < 0.001) and empathy concern (Experiment 1: *H*(2) = 4.21, *p* = 0.122; Experiment 2: *H*(2) = 21.32, *p* < 0.001). According to the personality questionnaires, the rational moralists were the most selfish and the justice warriors had the highest empathy (see Fig. S13 for post-hoc tests), which echoes the highest inaction inequality discounting (*η*_*no*_) in the former and highest action inequality discounting (*η*_*yes*_) in the latter (Fig. 4h and 4f). We also reported some exploratory analyses on cultural differences in the Supplement.

### The motive cocktail model quantitatively reproduces a broader range of phenomena

One may wonder whether the motive cocktail estimated in participants’ Intervene-or-Watch decisions is specific to this specific third-party intervention task. To demonstrate that such motive cocktail underlies human responses to inequality in general, we performed an out-of-sample prediction, using the motive cocktail model (with slight adaptations) to simulate the behavioral patterns in published studies with different experimental settings (Fehr & Fischbacher, 2004; Stallen et al., 2018). Indeed, we found that the motive cocktail model can predict the behavioral patterns in second-party punishment (2PP) as well as 3PP and 3PH (Fig. 6).

One robust phenomenon is that interveners spend more to penalize transgressors when they themselves are victims rather than unaffected third parties (i.e., 2PP > 3PP). This can be explained by the motive of deterrence (Delton & Krasnow, 2017), which is not in conflict with our utility maximization framework. We integrate this by assuming that deterrence motives lead to reduced efficiency concern (parameter ω) in second-party situations. More broadly, ω may decreases with social distance (Tang et al., 2023) and intent viciousness (Gummerum & Chu, 2014).

In our simulations, we model second-party interveners as having all the motives of third-party interveners except efficiency concern (i.e., ω=0, see Methods). Using parameters estimated from Experiment 1 participants, our model reproduces both the 2PP>3PP phenomenon and the increase in punishment with increasing inequality observed in previous laboratory experiments (Fehr & Fischbacher, 2004; Stallen et al., 2018). For both experiments, simulations with the justice warriors’ parameters best matched the data.

Stallen et al. (2018) used a scenario where the first party robs the second party. The inequality here was caused by the more vicious intentions of the transgressor, thus triggering stronger third-party punishment than the same level of inequality caused by a dictator allocator (See Fig. S14). For this case, we assume even unaffected third parties have no efficiency concern, allowing our model to reproduce the less common 3PP > 3PH phenomenon they observed.

## Discussion

It has been widely known that people have an innate aversion for inequality, willing to change others’ payoffs at the expense of their own, even when the action does not augment their reputation or encourage future cooperation (Dawes et al., 2007). What we have demonstrated here, through 100 different experimental conditions and a total of 1415 participants, is that such altruistic behaviors may result from a trade-off among as many as seven socioeconomic motives. Moreover, we have shown the plausibility to disentangle the effects of these different motives via computational modeling. The motive cocktail model we constructed here can explain a wide range of phenomena observed in the literature as well as in our two experiments. Overall, this study advances our knowledge on human altruistic behaviors by applying systematic and quantitative empirical tests in a unified modeling framework. It demonstrates how each motive, as well as their collective influence, shapes the behavioral outputs that ultimately contribute to individual differences in 3PP and 3PH.

Part of the phenomena found in our experiments and the literature, such as the increased third-party intervention with the increased inequality between the transgressor and the victim (Egas & Riedl, 2008; Jordan et al., 2014; Stallen et al., 2018; Zhong et al., 2016) and with the increased impact-to-cost ratio of the intervention (Egas & Riedl, 2008), can be readily explained by previous models of inequality aversion as well as by the three different inequality aversion terms in our model.

However, inequality aversion alone cannot explain why participants in our experiments were more likely to help the victim than to punish the transgressor, a finding in line with many previous studies (Batistoni et al., 2022; FeldmanHall et al., 2014; Singh & Garfield, 2022; Wiessner, 2020). Helping and punishment are equally efficient in reducing victim-centered inequality but differ in their influences on self-centered inequality, which would lead to a preference of punishment over helping, unless one is more uneasy about their advantage over others than the reverse. What enables our motive cocktail model to explain the preference for helping over punishment is the inclusion of efficiency concern as a utility term, that is, people also care about the overall payoff of the transgressor and the victim.

With an additional assumption that the motive of efficiency concern is weakened when oneself is the victim or when the transgressor violates social norms in a more aggressive way such as robbing or stealing from the victim (Stallen et al., 2018; van Prooijen, 2009), we can also explain why people spend more resources for 2PP than for 3PP (Fehr & Fischbacher, 2004; Stallen et al., 2018) and why a preference for punishment than helping is found in some studies (Stallen et al., 2018; van Prooijen, 2009), as our simulation shows (Fig. 6). The simulation also reveals that the proportional increase of punishment with inequality can be the consequence of the joint effect of the motive cocktail, without any “willingness to punish” to be assumed. Our motive cocktail model thus provides a unified account for 2PP, 3PP and 3PH behaviors.

One motive documented in previous studies, which seems to go against inequality aversion, is rank reversal aversion (Li et al., 2022; Xie et al., 2017). Our motive cocktail model includes a generalized form of this motive and found that participants in our experiment prefer to reverse the initial inequality so that the victim has an advantage over the transgressor, similar to what happens in Shakespeare’s The Merchant of Venice. This reversal preference motive is opposite to rank reversal aversion and our finding suggests that the latter may only apply to situations where the inequality is not caused by the benefited party (Li et al., 2022; Xie et al., 2017).

While Waldfogel et al. (2021) identified individual differences in attention to others’ inequality, we found that even within the same individual, attention to others’ inequality can be systematically modulated by the personal cost associated with intervening. The sixth and seventh motive terms—the inequality discounting terms—distinguished our model from all previous models in that they assume interactions between cost and inequality aversion. In other words, different motives are not necessarily independent; the perceived inequality between others may be modulated by one’s self-interest.

In line with the joint functioning of multiple motives identified in our modeling analysis, we found a three-way interaction between cost, impact ratio, and the inequality between the transgressor and the victim. Such interaction was not reported in previous studies, probably because most studies used cost as a dependent rather than an independent variable, measuring the amount of money participants were willing to spend on the intervention, which would prevent such effects from being detected by usual statistical analysis. In contrast, the cost is manipulated by the experimenter in our task, resembling another type of real-world scenarios where individuals are confronted with limited options when it comes to addressing others’ inequalities.

The two forms of inequality discounting—inaction inequality discounting and action inequality discounting—have distinct psychological implications. The former assumes that people act as if they are increasingly likely to ignore the inequality imposed on the victim due to the increasing cost of intervention. Consequently, they may be reluctant to engage in altruistic actions that pose potential harm to their own interests. The latter form of inequality discounting assumes that people act as if ignoring the remaining inequality faced by the victim after their intervention. As a result, they may be willing to intervene even when their intervention hardly improves equality. The co-existence of these two types of inequality discounting demonstrates the motive diversity in altruistic behaviors across various social contexts. These findings have significance for addressing real-world social issues. For example, in term of policy formulation, reducing barriers and costs for reporting injustices and corruption can encourage public engagement against social inequities; emphasizing the resolution achieved by intervention actions, such as highlighting how small acts of kindness can effectively mitigate real social issues, can further encourage altruistic behavior.

In both the laboratory experiment (N=157) and the large-scale online replication (N=1258), we found diverse behavior patterns across individuals. Three types of interveners—justice warriors, pragmatic helpers, and rational moralists—differed in their probability of intervention, their sensitivity to variables such as cost and inequality, and their preference for helping over punishment. The clustering in behaviors we observed aligns with the findings of Fehr et al., (2023) that the majority of individuals possess some form of pro-social preference and few are purely self-interested. The participants in this study extend beyond the typical WEIRD populations (Western, Educated, Industrial, Rich, Democracies) that are often overrepresented in research, thereby improving the generalizability of our findings to the broader population. The motive parameters estimated from the motive cocktail model provides a multi-facet measure of such individual differences, opening questions such as how individuals’ motives may depend on their personal experiences, cultural background, or genetic makeup.

The “motive cocktail” model proposed in this study extends the economic modeling of altruistic behaviors and has important implications for understanding and promoting prosocial behavior on a societal level. By elucidating the cognitive processes underlying prosocial behavior and identifying new motives and individual differences, our model can provide insights into psychiatric disorders characterized by social dysfunction and inform future research on the neural basis of human morality and its disorders (Sanders, 2021). Our model and task framework can also be used to investigate the developmental trajectories of altruistic motives, guiding efforts to foster prosocial behaviors across life stages (Lockwood et al., 2021). By capturing the interplay of multiple motives and their impact on behavioral patterns, our model enables more precise predictions of prosocial behavior. Leveraging insights from the “motive cocktail” model, interventions can be designed to account for the diverse motivations and experiences of individuals within society as well as cross culture background (Claessens et al., 2024), with the goal of creating a more cohesive and prosocial community. Meanwhile, further research needs to cover the gap between our over-simplified laboratory task and real-world applications.

### Limitations

We used a one-shot anonymous interaction setting, a common practice in previous studies (Dawes et al., 2007; Fehr & Charness, 2023; Fehr & Fischbacher, 2004; Fehr & Gächter, 2002; FeldmanHall et al., 2014; Rockenbach & Milinski, 2006; Stallen et al., 2018; van Baar et al., 2019; Wang et al., 2024; Zhong et al., 2016), to minimize participants’ concern for their own reputation, a motive that is instrumental to the long-term reciprocity in human society (Milinski et al., 2002). Consequently, our motive cocktail model, which adequately explained our data, excluded reputation as a motive. But in real-world scenarios with more interaction opportunities, reputation concern is likely to influence 3PP and 3PH behaviors (Bénabou & Tirole, 2006; Jordan et al., 2016). The victim’s reputation (e.g., once a transgressor or not) also matters, with reputation-based expectancies emerging early in human development (Ting et al., 2019). Similarly, deterrence (Delton & Krasnow, 2017), reciprocity (Gintis, 2000), or social norms beyond egalitarian distribution (Kimbrough & Vostroknutov, 2016) are other real-world motives not examined here. Integrating these motives into the motive cocktail model will be topics for future research. Whether the three types of interveners relate to the different cooperative types found in public goods games (Kurzban & Houser, 2001), thus connecting to a larger picture of human altruistic behaviors, also deserves future research.

## Methods

### Experiment 1

#### Participants

Experiment 1 was conducted in a laboratory room at Beijing Normal University and 157 university students (59 males, mean age ± SD: 21.24 ± 2.56) were recruited. Participants completed the screening form before the task to confirm that they had normal or corrected-to-normal vision and no history of psychiatric or neurological illness. All participants provided informed consent. This study was approved by the ethics committee of Beijing Normal University.

#### Experimental procedure

Participants were self-paced to read the instructions of the task. A quiz followed the completion of each sub-section of the instruction. Participants proceeded to the next section of the instruction only if they gave the correct answer to the quiz. Before the formal task, participants underwent several practice trials to ensure that they fully understood the rules of the game. The Intervene-or-Watch task (detailed below) lasted approximately 45 minutes. After completing the task, participants were asked whether they had any doubts or questions during the task in an open-ended question. In Experiment 1, four participants reported doubts about whether all the players were real people. To examine whether participants who reported doubts used different strategies compared to those who did not have doubts during the task, we conducted a generalized linear mixed model similar to GLMM1 but added “doubt” as an additional predictor (a categorical variable) in the model. We found that the predictor “doubt” couldn’t predict participants’ choice (b = –2.74, 95% CI [–7.19, 2.24], *p* = 0.304). Thus, we concluded that participants who reported doubts did not employ different strategies in the task. Therefore, all participants were included in the following analysis. In the final section, participants were asked to fill out a few personality questionnaires (detailed below), including measures of social value orientation (SVO), Machiavellianism Scale (MACH–IV), and interpersonal reactivity index (IRI), to assess their prosocial personalities.

#### The Intervene-or-Watch task and experimental design

The Intervene-or-Watch task was a paradigm adapted from a third-party punishment task (Fehr & Fischbacher, 2004). In the task, participants played the role of an unaffected third party who watched an anonymous dictator (“transgressor”) allocated amounts between himself/herself and an anonymous receiver (“victim”), and then decided whether to intervene. On each trial, the transgressor allocated the 100 game tokens between himself/herself and the victim, while the victim had to accept the offer without any other options. Participants were told that all offers between a transgressor and a victim were made by other real participants, and that their decisions would affect their own payoffs as well as those of the victims and the transgressors. In reality, the offers between the transgressors and the victims were generated by a custom code and were designed to disentangle different hypotheses. To give the participants a more realistic experience and to familiarize them with the roles in the game, they were instructed to play two trials of the dictator game, in which they played the role of transgressor and victim respectively. In the Intervene-or-Watch task, participants had 50 game tokens in each trial which could be used to reduce the payoff of the transgressor in the punishment scenario or increase the payoff of the victim in the helping scenario. To avoid serial or accumulative effects, participants were instructed that their payoff was independent across trials and would not be accumulated through the task. They were also informed that 10% of the trials would be randomly selected and implemented at the end of the study to determine the payoffs of all players (or roles). Specifically, participants’ actual payment was calculated by adding a base payment to the average remaining tokens from these randomly selected trials, with each token being exchanged for 1 yuan. Additionally, participants were explicitly informed that the roles of the transgressor and the victim were played by different participants in each trial, hence encouraging them to make decisions based solely on the current situation. We are aware that our experimental setting included deception, in the sense that participants’ intervention to the players in the dictator game was not real implemented. Nevertheless, all of the offers we used in the Intervene-or-Watch task were what real human players may make in the dictator game (Doñate-Buendía et al., 2022; Engel, 2011). Such use of deception is a common practice of previous studies (FeldmanHall et al., 2014; Stallen et al., 2018). Furthermore, participants’ payoff was actually determined by the randomly selected 10% of their decisions, akin to a random lottery design (Bardsley, 2000), which did not involve deception.

Since all players in the task were anonymous, no reputation concern was involved in this task. The players also had no opportunities for interaction; thus, reciprocity could be excluded. Therefore, participants’ decisions to help and to punish in the Intervene-or-Watch task were altruistic.

Each trial (Fig. 1c & 1d) began with a fixation cross (600–800 ms), followed by an inequality window (1500 ms) displaying the allocation between the transgressor and the victim, and an intervention offer window (1500 ms) showing the intervention cost for the participant and the consequence of the intervention (impact ratio × intervention cost) to the transgressor or victim. Subsequently, in the decision window, participants were asked whether they would like to accept the intervention offer: Yes (to intervene) or No (not to intervene). The intervention would only be implemented if participants chose Yes. For example, if the intervention offer window displays an intervention cost of x in a trial, a decision of intervention would result in the transgressor losing (or the victim gaining) 1.5x or 3.0x in the punishment (or helping) scenario. There was no time limit for the decision. A visual feedback window after the decision highlighted the selected choice in red. Four independent variables were varied across trials: scenario (punishment and helping), inequality (transgressor vs victim, 50:50, 60:40, 70:30, 80:20, 90:10, jitter ± 2), cost (10, 20, 30, 40, 50), and impact ratio (1.5 and 3.0). This led to 100 unique conditions, with each condition repeated 3 times for each participant. The scenario variable varied between blocks and the other three variables were randomly interleaved within blocks. Before each block, participants were told whether the following section was the “increase” condition (the helping scenario) or the “reduce” condition (the punishment scenario). In total, each participant completed 300 trials in 6 blocks, with 3 blocks each for the punishment and helping scenarios. The main experiment of the Intervene-or-Watch task lasted 30.86 ± 3.25 minutes for Experiment 1 and 33.97 ± 7.59 minutes for Experiment 2. The main experiment included 6 blocks, with each block lasting around 5 minutes, followed by a 30-second rest between blocks.

##### Personality questionnaires

Following the Intervene-or-Watch task, participants completed several personality questionnaires that allowed us to access their prosocial tendencies in daily life. Specifically, a Social Value Orientation (SVO, Murphy et al., 2011) was used to measure individual preference about how to allocate financial resources between themselves and others. A higher score on the SVO scale reflects a greater degree of concern for others’ payoffs and, therefore, indicates a more prosocial personality. A Machiavellianism Scale (MACH–IV, Rauthmann, 2013) was used to assess an individual’s level of Machiavellianism, related to manipulative, exploitative, deceitful, and distrustful attitudes. Individuals with higher scores on the MACH–IV scale are indicative of a more pronounced degree of Machiavellian traits. An Interpersonal Reactivity Index (IRI, Davis, 1983) was used to measure the multi-dimensional assessment of empathy, including (1) perspective-taking, assessing an individual’s tendency to consider a situation from others’ perspective; (2) fantasy, evaluating an individual’s inclination to identify with the situation and emotions of characters in books, movies, or theatrical performances; (3) empathy concern, measuring an individual’s inclination to care about the feelings and needs of others; (4) personal distress, assessing an individual’s tendency to experience distress and discomfort in challenging social situations.

##### Model-free analysis

All statistical analyses (except for group-level Bayesian model comparisons) were conducted in R 4.2.1 (R Core Team, 2013). Generalized linear mixed models (GLMM) assuming binomial distributed responses were used to model the probability of intervention, given various predictors (e.g., scenario, inequality) and their interactions. The GLMMs were implemented by “lme4” package (Bates et al., 2015), with the fixed-effect coefficients output from the binomial GLMM on the logit scale and the significance of each coefficient determined by the z-statistics. The standard linear mixed-effect models (LMM), which assume the error term is normally distributed, were estimated using the “afex” package to model participants’ decision times. For the estimation of marginal effects and the post hoc analysis, the “emmeans” package was used (Lenth et al., 2018). Interaction contrasts were performed for significant interactions and, when higher-order interactions were not significant, pairwise or sequential contrasts were performed for significant main effects.

##### GLMM1

participants’ choices of all trials in Experiment 1 are the dependent variable; fixed effects include an intercept, the main effects of the scenario, inequality, cost, ratio, trial number, and all possible interaction effects of the independent variables; random effects include correlated random slopes of scenario, inequality, cost, ratio and trial number within participants and random intercept for participants. The scenario is a category variable. Trial number, inequality, cost, and ratio are continuous variables that were normalized to Z-score prior to model estimation. The inclusion of trial number controls for time-related confounds, such as potential fatigue or practice effects. See Table S1 for the statistical results of GLMM1.

##### GLMM2

participants’ choices of all trials in Experiment 2 are the dependent variable. The fixed and random effects remain the same as GLMM1. See Table S5 for the statistical results of GLMM2. Both the main and interaction effects of the independent variables on intervention decisions of Experiment 1 (as in Fig. 1e–l) were replicated in Experiment 2 (Fig. S15a–m).

##### LMM1

participants’ decision times of all trials in Experiment 1 are the dependent variable. In addition to the fixed and random effects included in GLMM1, participants’ intervention decisions (choice) are added as well. See Table S6 and Fig. S16 for the statistical results of LMM1.

We found an inverted U-shaped relationship between the intervention probability (*p*(yes)) and decision time (Fig. S16j), which implies that participants made decisions with more difficulty when the decision uncertainty (or entropy) was higher. This result is in line with prior research demonstrating an inverted U-shaped relationship between confidence levels and decision times (Ratcliff & Starns, 2013).

##### LMM2

participant’s decision times of all trials in Experiment 2 are the dependent variable. The fixed and random effects remain the same as LMM1. See Table S7 for the statistical results of LMM2. The inverted U-shaped relationship between the probability of intervention (*p*(yes)) and decision time was replicated in Experiment 2 (Fig. S17).

##### The sensitivity analysis to different variables

We measured participants’ intervention sensitivity to different variables, which was defined as the normalized intervention probability difference after the corresponding variable was dichotomized (Fig. 4n-r and Fig. S18). Specifically, participants’ sensitivity to the main effects, including scenario, ratio, cost and inequality, was calculated as the intervention probability difference in the helping trials compared with the punishment trials, the high-impact ratio trials (i.e., 3.0) compared with the low-impact ratio trials (i.e., 1.5), the low-cost trials (i.e., cost<=20) compared with high-cost trials (cost>20), and the high-inequality trials (i.e., the inequality level between the transgressor and the victim is 80:20 and 90:10) compared with the low-inequality trials (i.e., 70:30, 60:40 and 50:50), divided by their overall *p*(yes), respectively. For the interaction effects, the sensitivity (i.e., the normalized intervention probability difference) was calculated in a similar way as the main effect, that is, marginalizing over the other variables.

#### Behavioral modeling

We assumed that participants would make decisions on each trial by calculating the utility of the two options (yes and no) and choosing the option with the higher utility. In the Intervene-or-Watch task, participants were given the context regarding inequality between a transgressor and a victim as well as other related variables (e.g., cost, impact ratio) from the perspective of a third-party and afterwards made a decision between two alternatives, yes (to intervene) and no (not to intervene). In general, participants calculated the utilities of the choices by estimating the reduction in inequality for others through their intervention and considering the associated cost to themselves. Specifically, if they chose “yes” (decide to intervene), they could reduce the inequality between the transgressor and the victim to some extent but at a cost. On the contrary, by choosing “no” (decide not to intervene), they could keep the inequality between the transgressor and the victim without incurring any cost. To investigate how individuals make decisions in the Intervene-or-Watch task, we constructed a series of computational models with different utility calculation hypotheses (i.e., combinations of multiple socioeconomic motives) and compared their goodness of fit.

Participants’ choices were then modelled using the Softmax function (Eq. 1, Luce, 1959), with the utilities of not intervention (*U*_*no*_) and intervention (*U*_*yes*_) from different models as the inputs:

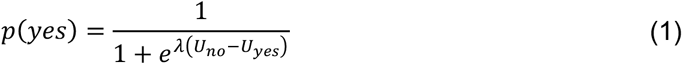

where the inverse temperature, parameter *λ* ∈ [0, 10], controls the stochasticity of participants’ choices, with a larger λ corresponding to less noisy choices.

In the following descriptions, we will use *x*_1_, *x*_2_ and *x*_3_ to denote the payoffs of the transgressor, the victim, and the third-party (participant) if the third-party does not intervene (choose ‘no’), and use 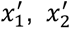 and 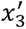 to denote the counterpart payoffs if the third-party intervenes (choose ‘yes’). In particular, 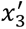 is equal to *x*_3_ − *cost* in both scenarios. In the punishment scenario, 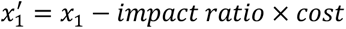 and 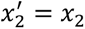. While in the helping scenario, 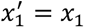 and 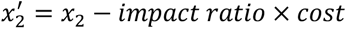.

##### Model 1: The baseline model

We modeled each participant’s choices of intervention in each trial (i.e., whether to choose the yes option) as outcomes from a Bernoulli distribution, where the intervention probability is controlled by a parameter, *q* ∈ [0, 1]. For each participant, the probabilities of choosing the intervention (*p*(yes)) and not choosing the intervention (*p*(no)) are denoted as follows:

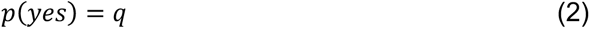

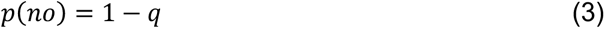

##### Model 2: Self-interest model (SI)

The models based on socioeconomic motives started with SI, where participants only consider self-interest when making decisions and thus, always leading to a reduced utility of the intervention. Participants’ choices were then modeled using the Softmax function (Eq.1).

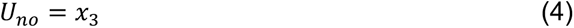

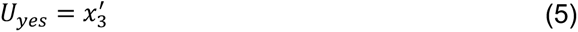

where *x*_3_ denotes the payoff of the third-party when choosing “no” (without intervention), which is always 50 tokens in each trial. 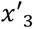 denotes the payoff of the third-party after choosing “yes” (with intervention), equaling to 50 − *cost*.

Building upon the SI model, the following hypothetical socioeconomic components were progressively introduced into the utility calculation and participants’ choices were modeled using the Softmax function. The necessity of each component to explain participants’ decisions was determined through model comparisons.

##### Model 3: SI and self-centered inequality aversion model (SI+SCI)

Based on SI model, we added a self-centered inequality aversion component, which assumes that participants are averse to the inequality between themselves and others in both directions (Fehr & Schmidt, 1999). The self-centered Disadvantageous Inequality (DI) aversion corresponds to that participants are averse to others having more payoffs than themselves, while the self-centered Advantageous Inequality (AI) aversion denotes participants are averse to themselves having more payoffs than others. The contributions of self-centered disadvantageous and advantageous inequality are controlled (Fehr & Schmidt, 1999)separately by the parameters *⍺* (*⍺* ∈ [0, 10]) and *β* (*β* ∈ [0, 10]) and are subtracted from the self-interest. Under the assumption of the SI+SCI model, participants are motivated to maximize their self-interest and meanwhile, minimize the inequality between themselves and others and then make a choice between no intervention and intervention based on their respective utilities:

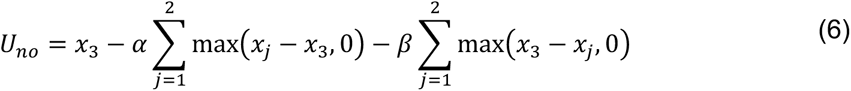

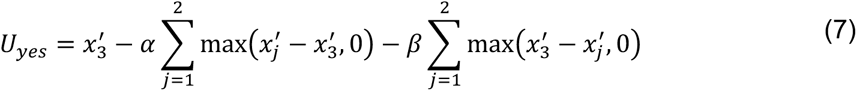

where *j* denotes the index of the transgressor and victim; *x*_1_ and *x*_2_ represent the payoffs of the transgressor and the victim when participants (the third-party) choose “no”; 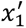 And 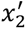 represent the payoffs of the transgressor and the victim after the intervention of the third party.

##### Model 4: SI+SCI and victim-centered disadvantageous inequality aversion model (SI+SCI+VCI)

Based on the SI + SCI model, we introduced another previously proposed inequality component, the victim-centered disadvantageous inequality aversion (VCI). The VCI assumes that participants are averse to the transgressor having more payoff than the victim (Zhong et al., 2016), with its contribution to the utility calculation determined by a parameter *γ* (*γ* ∈ [0,10]). Participants with larger *γ* will be more willing to intervene in almost all punishment and helping scenarios. Within this model, participants were motivated to maximize self-interest and simultaneously, minimize the two kinds of inequality aversions (SCI and VCI):

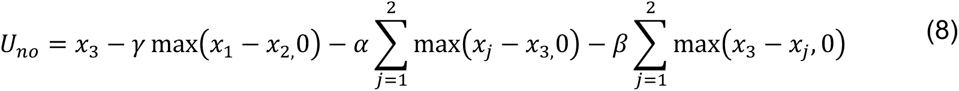

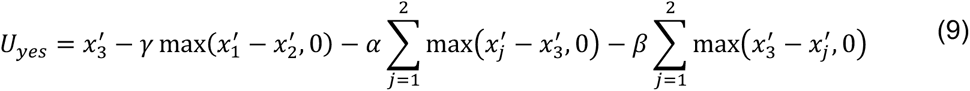

##### Model 5: SI+SCI+VCI and efficiency concern model (SI+SCI+VCI+EC)

Based on the SI+SCI+VCI model, efficiency concern (Engelmann & Strobel, 2004) component was added into the model. The efficiency concern (EC) assumes that participants are motivated to maximize the total payoff of others, which is weighted by parameter *ω* (*ω* ∈ [0,10]). Participants with larger ω will more likely intervene in the helping scenario, but not in the punishment scenario:

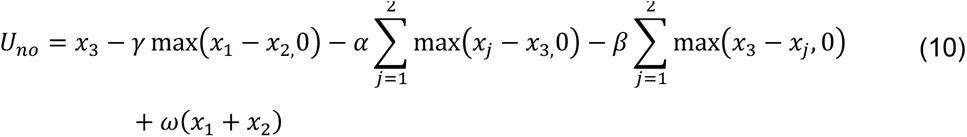

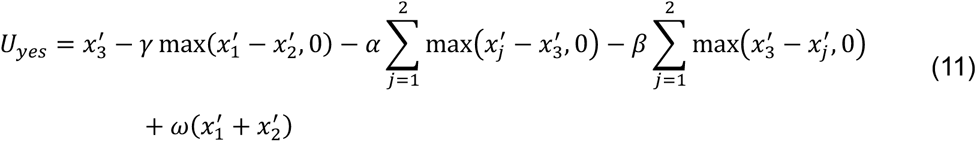

##### Model 6: SI+SCI+VCI+EC and reversal preference for victim-centered advantageous inequality model (SI+SCI+VCI+EC+RP)

Based on the SI+SCI+VCI+EC model, we introduced another component, the reversal preference for victim-centered advantageous inequality (RP), into the model. RP is mutually exclusive to VCI and assumes that participants prefer to reverse the economic status of the victim. That is, RP motivates participants to make the victim have more payoff than the transgressor by punishing the transgressor or helping the victim. The reversal preference is controlled by the parameter *κ* (*κ* ∈ [−10,10]). A positive value of *κ* indicates that participants are in favor of the victim having more money than the transgressor, while a negative value indicates that they are averse to the newly created reverse inequality. Participants with larger *κ* will more likely intervene when the initial victim-centered disadvantageous inequality is small enough or the impact is large enough to guarantee an inequality reversal:

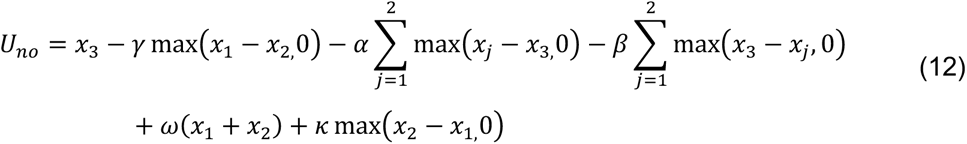

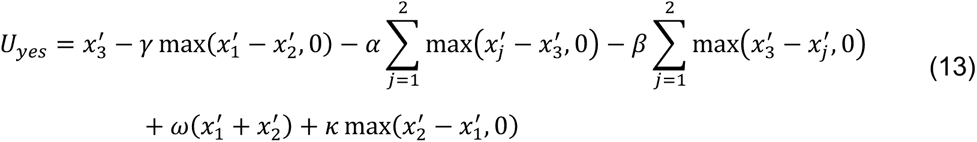

##### Model 7: SI+SCI+VCI+EC+RP and inequality discounting model (the motive cocktail model, SI+SCI+VCI+EC+RP+ID)

Based on the SI+SCI+VCI+EC+RP model, we included a newly proposed inequality discounting (ID) component. Thus, the motive cocktail model includes seven socioeconomic motives. ID is derived from the rational framework of economic decisions and is implemented to capture the interaction between self-interest and VCI. Specifically, ID assumes that people will systematically disregard the victim-centered disadvantageous inequality as costs increase. We proposed two types of inequality discounting: inaction inequality discounting (IID, controlled by parameter *η*_*no*_) and action inequality discounting (AID, controlled by *η*_*yes*_), which are respectively blind to the initial and residual disadvantageous inequalities between the transgressor and the victim under no intervention and intervention with rising costs, respectively. In the model fitting, the range of parameters *η*_*no*_ and *η*_*yes*_ is restricted to between 0 and 20.

Participants with larger *η*_*no*_ would have a lower probability to intervene. The effect differs from victim-centered disadvantageous inequality aversion (small γ) in that at large *η*_*no*_ the tendency to intervene would barely increase with inequality. Conversely, participants with larger *η*_*yes*_, who subjectively exaggerate the reduction of inequality by intervention, would have a higher probability to intervene. Those with large *η*_*yes*_ would have similarly high probability to intervene regardless of the impact ratio, as if they optimistically believe that the inequality would be minimized by any of their intervention:

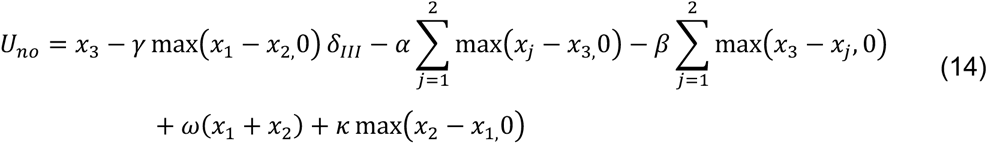

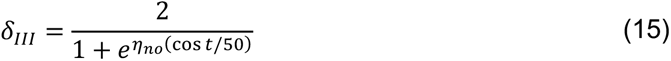

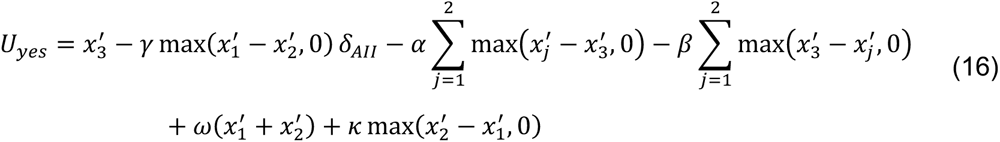

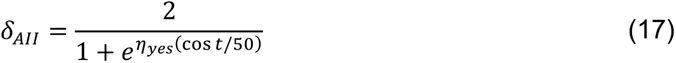

#### Redundancy checks on the parameter space for the motive cocktail model

In the estimated parameters, we observed three highly correlated pairs in the parameter space of the motive cocktail model: the value of parameter *β* (self-centered advantageous inequality aversion) and *γ* (victim-centered disadvantageous inequality aversion), *⍺* (self-centered disadvantageous inequality aversion) and *ω* (efficiency concern), *γ* and *η*_*no*_ (inequality inaction inattention). To exclude the possibility that the correlation was due to parameter redundancy in the model, we performed redundancy checks as follows. We first randomly shuffled participants’ labels for different parameters to eliminate correlations in the shuffled parameters. Based on these shuffled parameters, we generated 157 synthetic datasets and used them to estimate the model parameters. We found little correlations between the parameters estimated from these synthetic datasets, which indicates that the high correlations found in the data reflect the behavioral characteristics of human participants rather than redundancy in the model itself (Fig. S19).

#### Model fitting and model comparison

For each participant, we fit each model to their intervention decisions across all trials using maximum likelihood estimates. The likelihood function derived from the binomial distribution was used to describe the relationship between participants’ choice and the model’s prediction. The function *fmincon* in MATLAB (MathWorks) was used to search for the parameters that minimized negative log-likelihood. To increase the probability of finding the global minimum, we repeated the search process 500 times with different starting points. We compared the goodness of fit of each model based on two metrics: Akaike information criterion with a correction for sample size (AICc) (Hurvich & Tsai, 1989) and the protected exceedance probability of group-level Bayesian model selection (Rigoux et al., 2014). We chose to use the AICc as the metric of goodness-of-fit for model comparison for the following statistical reasons. First, BIC is derived based on the assumption that the “true model” must be one of the models in the limited model set one compares (Burnham et al., 2002; Gelman & Shalizi, 2013), which is unrealistic in our case. In contrast, AIC does not rely on this unrealistic “true model” assumption and instead selects out the model that has the highest predictive power in the model set (Gelman et al., 2014). Second, AIC is also more robust than BIC for finite sample size (Vrieze, 2012).

#### Model identifiability and parameter recovery analyses

We further performed a model identifiability analysis to rule out the possibility of model mis-identification in model comparisons. For each model, the parameters estimated from the data of all participants were used to generate a synthetic dataset of 157 participants. Each synthetic dataset regarding a specific model was then used to fit each of the 7 alternative models and identify the best fitting model by model comparison. We repeated the above procedure 100 times to calculate the percentages that each model was identified as the best model based on all synthetic datasets from a specific generating model. The highest percentage assigned to the same fitting model as the generating model suggests that the model is identifiable. To assess parameter recovery in the motive cocktail model (model 7: SI+SCI+VCI+EC+RP+ID), we computed the Pearson correlation between the parameters estimated from the 100 synthetic datasets (recovered parameters) and the parameters used to generate the synthetic datasets. The larger correlation coefficient between the recovered parameter and the estimated parameter indicates a non-redundancy in parameter space.

#### Clustering analysis

To gain further insight into whether the motive cocktail model (model 7: SI+SCI+VCI+EC+RP+ID) could explain the varying behavioral patterns of individuals, we classified participants’ intervention decisions using K-means clustering and then investigated the distributions of the estimated parameters across participants as well their unique contributions to behavioral patterns within each cluster. K-means clustering is an unsupervised machine learning algorithm relying on the Euclidean distance to classify each participant into a specific cluster with the nearest mean (Hartigan & Wong, 1979). The clustering evaluation criterion was based on silhouette value which denotes how well each participant was matched to its own cluster compared to other clusters, with a higher silhouette value indicating the clustering solution is more appropriate (Rousseeuw, 1987). The optimal cluster solution for 157 participants in Experiment 1 is 3 (Fig. 4a).

#### Correlation analysis between the estimated parameters and behavioral or personality measures

To further validate the psychological significance of the hypothetical socioeconomic motives in the motive cocktail model, we calculated Pearson’s correlation between the estimated parameters and the scores on the personality measurements. A similar correlation analysis between individuals’ motive parameters and their sensitivity to different variables was carried out to unravel the contributions of the parameters to behavioral differences. Partial correlation was conducted when multiple parameters correlated with the same measurement to ensure that the observed relationships were not confounded by the potential influence of other variables. The partial correlation coefficient (*ρ*), ranging between –1 and 1, quantifies the strength and direction of linear links between parameters and measured variables. For multiple comparisons, the False Discovery Rate (FDR) was employed.

#### Simulations to quantitatively reproduce a broader range of phenomena

We made slight modifications to the motive cocktail model and applied it to explain the intervention patterns in second-party punishment (2PP), third-party punishment (3PP), and third-party helping (3PH) models in the following two studies. The adapted model could also be used to explain a broader range of phenomena in previous studies.

In a sub-study conducted by Fehr and Fischbacher (2004), participants attended a dictator game, which contains both 2PP condition and 3PP condition. At the beginning of the experiments, participants were randomly assigned either the role of the transgressor (Player A) or the victim (Player B). In the 2PP condition, the victim also acted as an intervener who could punish his transgressor after observing the transfer from the transgressor accordingly. In the 3PP condition, the victim could only punish the dictator in another group (Player A’ and Player B’), in which he/she served as an unaffected third party. A strategy method was implemented in the 3PP condition: the third-party (Player B) had to indicate how much he would punish the outgroup Player A’ for every possible transfer of A’ to Player B’. The results showed that the intervener as the victim exerted more punishment than the intervener as the third-party for all transfer levels below 50 (i.e., 2PP > 3PP), while the punishment was generally low and similar across transfer levels above 50 (Fig. 6a top left). In the study conducted by Stallen et al. (2018), participants played three conditions of a justice game. In the 2PP games, the participants played the role of the partner (the victim), in which the taker (the transgressor) had the opportunity to take or rob chips (or payoff) from the victim, and afterward, the victim was given the option of punishing the transgressor by spending chips of their own. In 3PP and 3PH games, participants played the role of an observer (the third-party) to watch whether the transgressor robbed chips from the victim and then decided whether to intervene to punish the transgressor or to compensate the victim, at their own cost. Every time participants needed to make a choice, all intervention costs ranging from 0 to 100 with a step of 10 were displayed on the screen. The results indicated the intervener in the 2PP condition punished more on the transgressor than in the 3PP condition (i.e., 2PP > 3PP). In addition, the third-party was more likely to punish than to compensate (i.e., 3PP > 3PH, Fig. 6b top left).

For both studies, we simulated participants’ choices by calculating the utility of selecting “yes” and “no” for each inequality level using Eqs.14–17. We assume that a second-party intervener, who themselves are also the victim, is less concerned about the overall welfare than the third-party does. As the result, the second-party intervener has all the motives a third-party intervener would have except for efficiency concern. To implement this assumption, we replaced x_3_ in Eqs. 14–17 with x_2_, and set the efficiency concern ω to 0 in the second-party punishment condition. The same lack-of-efficiency-concern assumption (i.e., ω = 0) was implemented during the simulation of third-party punishment and compensation games in Stallen et al. (2018). That is, we assume that the unaffected third party would ignore others’ welfare in a robbery situation.

### Experiment 2

To further verify our findings and model specifications, we conducted Experiment 2 using the same experimental paradigm as Experiment 1 on an online participant platform (Prolific, https://www.prolific.co/) by recruiting a larger population with diverse cultural backgrounds.

#### Pre-registration

Experiment 2 was pre-registered on OSF (https://osf.io/gcsqp) on September 29, 2022. All methods and analyses followed the design and analysis plan in the preregistration, except that two additional models were tested: a model with lapse rate parameters and a simple-response model. This was due to more behavioral patterns being observed from the online experiment. The Building on the results of the model-free analysis in Experiment 1, we hypothesized that the main effect of inequality, intervention cost, impact ratio and the interaction of inequality × cost × ratio would be statistically significant, and that participants’ intervention decisions would follow the patterns we observed in Experiment 1. For the model-based analysis, we hypothesized that participants’ decisions would be best described by the full motive cocktail model.

#### Participants

The criteria for participant recruitment were matched between Experiments 2 and 1, including the age ranges (18–30 years old), student status and the degree of education. In addition, the study was only accessible to participants with an approval rate of over 90% in Prolific. We received 1365 participants’ submissions overall. One hundred and seven of them had an accuracy rate below 75% on the attention check task (see details below) and thus were rejected for further analysis. The final valid samples were 1258 (621 male, 631 female, 6 genders unknown, aged 23.30 ± 2.89). No participants met the exclusion criterion of average decision time exceeding 2.5 standard deviations from the mean decision time of all participants. All participants provided informed consent before the task to confirm they voluntarily took part in the study, had a normal or corrected-to-normal vision, and did not have a history of psychiatric or neurological illness. This study was approved by the Ethics Committee of Beijing Normal University.

#### Determination of sample size

The sample size for Experiment 2 was pre-determined using a parametric simulation method (Kumle et al., 2021), derived from the motive cocktail model (the best-fitting model in Experiment 1). The effect we focused on is the three-way interaction of inequality × cost × ratio (Fig. 1i). As a compensation for the higher randomness of online participants’ decisions, we added another two parameters, *p*_*m*i*n*_ and *p*_*max*_ (lapse rates), in the motive cocktail model to capture participants’ minimal and maximal (1 − *p*_*max*_) intervention probability. An online pilot study based on 32 participants showed the motive cocktail model with lapse rates (see Model 8 for more details) fit participants’ behavior better. We, therefore, used model 8 to generate synthetic datasets to determine the sample size for Experiment 2. Parameters *⍺*, *β*, *γ*, *ω*, *η*_*no*_, *η*_*yes*_, *λ* were sampled from Gamma distribution, *κ* was sampled from Normal distribution, and *p*_*m*i*n*_ and *p*_*max*_ were sampled from Beta distribution. The generated intervention decisions of virtual participants were then exported to GLMM1 to obtain the effect size for each variable and their interactions. The power was defined as the percentage that the three-way interaction effect reaches significance over a specific sample size. We tested different sample sizes ranging from 100 to 1500 virtual participants, with increments of 100. Within each sample size, we repeated the synthetic data generation and power calculation procedure 500 times. The power of the three-way interaction effect increased monotonically with sample size and achieved a power of 80% with at least 1200 participants (see Fig. S7). Our final valid sample size was 1258 participants from sixty-six countries (see Table S4).

#### Experimental procedure

The procedure of Experiment 2 was the same as that of Experiment 1, except that it was conducted on the Prolific platform. Participants were informed that their base payment was £7 per hour, and 10% trials would be randomly selected to determine their bonus after the experiment. The game tokens accumulated from these randomly selected trials would be exchanged for pennies at a 5:1 exchange rate. After the task, participants were asked “Did you think the experimenter had deceived you in any way at any point during the experiment?”, with a binary choice of “yes” or “no”. Seventy-four participants answered “yes”, while the remaining 1184 participants answered “no”. To investigate whether participants who had doubts (answered “yes”) employed different strategies compared to those who did not have doubts (answered “no”) during the task, we conducted a generalized linear mixed model (like GLMM2) and included “doubt” as a predictor (categorical variable) in the model. We found that the effect of doubt (b = 0.15, 95% CI [– 0.09, 0.41], *p* = 0.221) was not statistically significant to predict participants’ choices, suggesting that participants who reported doubts did not employ different strategies in the task. Therefore, all participants were included in the subsequent analysis.

#### Attention check

We used the same intervene-or-watch task in Experiment 2 and included several attention checks during the task to ensure participants remained constantly attentive to the current task. The attention checks consisted of 12 questions, with two questions interspersed in each block. For each block, the questions appeared randomly without telling the participants, and participants were asked to answer the questions with binary options about their last decision. Specifically, the questions were either “In the last trial, your decision was: yes/no?” or “The last trial was in the increase/reduce scenario?” in each block. For those (107 participants) who gave less than 75% accuracy in the attention checks (incorrect answers on more than three questions) were excluded from further analyses.

#### Model-free analysis

All 1258 participants in Experiment 2 were included in the model-free analysis (Table S5). Among them, 492 (39.10%) out of 1258 participants were best described by a simple-response model and were therefore excluded from the analyses in relation to the motive cocktail model. Specifically, only the remaining 60.90% of participants whose intervention patterns could be categorized as justice warriors, pragmatic helpers, and rational moralists were included in the following analyses: data versus model prediction (Fig. 5), Kruskal-Wallis tests on the parameters *η*_*yes*_, *κ*, *η*_*no*_ (Fig. S5), correlations between the parameters estimated from the motive cocktail model and the intervention sensitivities (Fig. S18) as well as the personality measurements (Fig. S11).

#### Behavioral modeling

##### Model space

We constructed two additional models (Models 8 and 9) in Experiment 2 to capture the behavioral patterns that online participants would make random choices in a certain amount of trials. Model 8 was constructed based on the motive cocktail model. Model 9 is a simple-response model to capture the behavioral patterns of a proportion of participants in the online Experiment 2 (39.10%) who only responded to part of the manipulated variables and seemed also entirely ignored the others.

##### Model 8: The motive cocktail model with two lapse rate parameters

The model assumes that participants make an intervention decision by considering both self-interest and all socioeconomic motives assumed in the motive cocktail model. However, participants’ minimal and maximal intervention probability are bounded by two free parameters. Specifically, participants are willing to randomly intervene with a probability of *p*_*min*_ (*p*_*min*_ ∈ [0, 0.5]). Meanwhile, they constrain their maximum intervention probability below 1 − *p*_*max*_(*p*_*max*_ ∈ [0, 0.5]). The utility calculations and choice mapping remain the same as Eqs.14-17 and Eq.1, respectively.

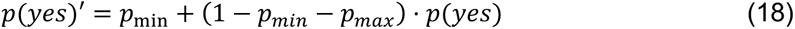

where *p*(*yes*) represents the choice probability based on the motive cocktail model.

##### Model 9: Simple-response model

Some of these online participants were sensitive to only a few of the manipulated variables and seemed to use simple-response rules for responses. Thus, we also included a simple-response model that linearly combines different manipulated variables (scenario, inequality, cost, and ratio) to describe participants’ behavior:

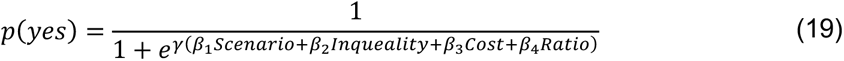

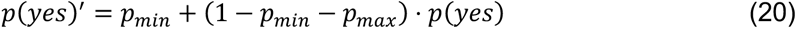

## Acknowledgements

We thank Ernst Fehr, Christian C. Ruff, Fiery Cushman and Jörg Gross for insightful discussions, Guoqiu Chen for helping with programming Experiment 2 and Haoyang Lu for statistical consultation. HZ was supported partly by the National Natural Science Foundation of China (32171095) and funding from Peking-Tsinghua Center for Life Sciences. CL was supported by the Scientific and Technological Innovation (STl) 2030-Major Projects 2021ZD0200500, the National Natural Science Foundation of China (32271092, 32130045), the Major Project of National Social Science Foundation (19ZDA363) and the Beijing Municipal Science and Technology Commission Z151100003915122.

## Author contributions

Conceptualization: XW, XR, CL, HZ.

Investigation: XW.

Data curation: XW, XR.

Formal analysis: XW.

Methodology-development and design of methodology: XW, XR, CL, HZ.

Methodology-creation of models: XW, XR, HZ.

Visualization: XW.

Writing – original draft: XW, XR.

Writing – review & editing: XW, XR, HZ, CL.

Funding acquisition: CL, HZ.

Supervision: CL, HZ.

## Data and code availability

All data and codes will be made publicly available upon acceptance of the paper. For review: https://osf.io/6g293/?view_only=3c7249b5ee67432fbc713628a26a1d39

