## Supplemental Data 1 for "The “motive cocktail” in altruistic behaviors"

#### Supplementary Methods and Results

#### Additional variants for the motive cocktail model

We introduced the concept of “inequality discounting” (II) to explain the observation that participants were still willing to intervene when they could only achieve a small intervention effect (cost × ratio) but were confronted with a large inequality between the transgressor and the victim. The inequality discounting term $\delta = \frac{2}{1+e^{\eta(cost/50)}}$ follows the form of a sigmoid function (Fig. S4b), which has the desired mathematical property of ensuring the value of $\delta$ being between 0 and 1. That is, the value of $\delta$ is 1 for 0 cost and approaches 0 for high cost, while the parameter $\eta$ controls the speed of this transition. Psychologically, this term can be interpreted as the probability or strength that the participant chooses to pay attention to a given transgressor-victim inequality, which, as a multiplying term for the magnitude of the inequality, modulates the effect of the latter on participants' intervention decisions. In a supplementary analysis, we tested four more variants of the full motive cocktail model (Model 7 in the main text) to demonstrate the necessity of the inequality discounting assumption (the interaction items) in the full model as well as the nonlinear modulation of self-interest on the victim-centered inequality term in fitting the behavioral data. Its results are presented in Figure S2.

The further comparison between different modulation forms of II demonstrated that the nonlinear assumption better captures the intervention patterns. Note that the II exerted its effects in the form of an interaction with inequality and only affected the utility calculation, whereas the response mapping remained the same in all models. This bifurcated analysis below, beyond the alternative models in the main texts, was designed to confirm the necessity of inequality discounting as well as its modulation form.

In the first line of analysis, the self-centered and victim-centered inequality were modulated by additional parameters respectively, which allowed us to test whether the assumption of interaction on inequality would improve the performance of the model.

**Variant model 1 (VM1)**. Based on model 6 (SI+SCI+VCI+EC+RP), the model assumes that the self-centered inequalities (SCI) are jointly modulated by two additional free parameters ( $\eta_{no}$ and $\eta_{yes}$ ), resulting in unequal contributions (or weights) of disadvantageous and advantageous SCI in the context of the intervention and not.

$$U_{no} = x_3 - \gamma \max(x_1 - x_{2,}0) - [\alpha \sum_{j=1}^{2} \max(x_j - x_{3,}0) + \beta \sum_{j=1}^{2} \max(x_3 - x_j, 0)]\eta_{no} + \omega(x_1 + x_2) + \kappa \max(x_2 - x_{1,}0) \quad \text{(S1)}$$

$$U_{yes} = x'_3 - \gamma \max(x'_1 - x'_2, 0) - [\alpha \sum_{j=1}^{2} \max(x'_j - x'_3, 0) + \beta \sum_{j=1}^{2} \max(x'_3 - x'_j, 0)]\eta_{yes} + \omega(x'_1 + x'_2) + \kappa \max(x'_2 - x'_1, 0) \quad \text{(S2)}$$

**Variant model 2 (VM2)**. Similar to VM1, VM2 assumes an invariant modulation to the victim-centered inequality (VCI) component by introducing another free parameter, $\eta$ .

$$U_{no} = x_3 - \gamma \max(x_1 - x_{2,}0) - \alpha \sum_{j=1}^{2} \max(x_j - x_{3,}0) - \beta \sum_{j=1}^{2} \max(x_3 - x_j, 0) + \omega(x_1 + x_2) + \kappa \max(x_2 - x_{1,}0) \quad \text{(S3)}$$

$$U_{yes} = x'_3 - \eta \max(x'_1 - x'_2, 0) - \alpha \sum_{j=1}^{2} \max(x'_j - x'_3, 0) - \beta \sum_{j=1}^{2} \max(x'_3 - x'_j, 0) + \omega(x'_1 + x'_2) + \kappa \max(x'_2 - x'_1, 0) \quad \text{(S4)}$$

Model comparison between VM1, VM2 and the first six models in the main texts showed that the model incorporating interaction items excelled the others (see Fig. S3), with VM2 performing the best. The analysis provided direct evidence that the inclusion of the interaction assumption (i.e., inequality discounting) is necessary and that the modulation is sensitive to victim-centered inequality aversion.

Following the conclusion of the modulation assumption, the next line of analysis focused on the modulation form of inequality discounting. Instead of an invariant effect, a variant modulation of victim-centered inequality was assumed. This assumption was derived from the rational framework of economic decision (Scott, 2000), where people systematically disregard victim-centered advantageous inequality as the intervention cost increase. Two forms of modulation were tested: diminishing linear and nonlinear.

**Variant model 3 (VM3)**. The model assumes that the modulation of self-interest to victim-centered inequality decreases linearly with increasing intervention cost, with the parameters $\eta_{no}$ and $\eta_{yes}$ controlling the modulatory magnitudes at different costs.

$$U_{no} = x_3 - \gamma \max(x_1 - x_{2,}0)\,\delta_{III} - \alpha \sum_{j=1}^{2} \max(x_j - x_{3,}0) - \beta \sum_{j=1}^{2} \max(x_3 - x_j, 0)\,\delta_{II} + \omega(x_1 + x_2) + \kappa \max(x_2 - x_{1,}0) \quad \text{(S5)}$$

$$U_{yes} = x'_3 - \gamma \max(x'_1 - x'_2, 0)\delta_{IIA} - \alpha \sum_{j=1}^{2} \max(x'_j - x'_3, 0) - \beta \sum_{j=1}^{2} \max(x'_3 - x'_j, 0) + \omega(x'_1 + x'_2) + \kappa \max(x'_2 - x'_1, 0) \quad \text{(S6)}$$

$$\delta_{III} = -\eta_{no}(cost/50) + 1 \quad \text{(S7)}$$

$$\delta_{IIA} = -\eta_{yes}(cost/50) + 1 \quad \text{(S8)}$$

**Variant model 4 (VM4).** The model assumes a nonlinear modulation of self-interest to victim-centered inequality.

$$\delta_{III} = \frac{2}{1 + e^{\eta_{no}(cost/50)}} \quad \text{(S9)}$$

$$\delta_{IIA} = \frac{2}{1 + e^{\eta_{yes}(cost/50)}} \quad \text{(S10)}$$

Both model comparison and model predictions indicated that the nonlinear assumption outperformed its alternatives in fitting behavioral data (Fig. S3). Therefore, the modulation form assumed in VM4 (i.e., the full model) was reported in the main texts.

### The proportion of participants who never chose the help/punish option

In our experiments, some participants never chose to intervene either in the punishment scenario, or in the helping scenario, or in both. Please see Table S8 for their proportions in each experiment.

As shown in Table S9, participants who neither punished nor helped were clustered into rational moralists. Those who never punished but sometimes helped were clustered into either rational moralists or pragmatic helpers, depending on their behavioral patterns in the helping scenarios.

### The statistical results of Experiment 2

Both the main and interaction effects of the independent variables on intervention decisions of Experiment 1 (as in Fig. 1e–l) were replicated in Experiment 2 (Fig. S15a–m and Table

S5). In particular, participants preferred helping over punishment (scenario $b = -1.19$, 95% CI [–1.32, –1.06], $p < 0.001$; Fig. S15a), and they intervened more often under higher inequality ($b = 0.59$, 95% CI [0.53, 0.64], $p < 0.001$; Fig. S15b), higher impact ratio ($b = 0.71$, 95% CI [0.65, 0.76], $p < 0.001$, Fig. S15c), and lower cost ($b = -0.78$, 95% CI [–0.84, –0.72], $p < 0.001$, Fig. S15d). We also found similar three-way interactions of cost × inequality × ratio ($b = -0.02$, 95% CI [–0.02 –0.01], $p = 0.021$; Fig. S15e–f), and two-way interactions of scenario × ratio ($b = -0.31$, 95% CI [-0.32, -0.29], $p < 0.001$; Fig. S15g) and cost × ratio ($b = 0.04$, 95% CI [0.03, 0.05], $p < 0.001$; Fig. S15i). The inequality × ratio interaction was marginally significant ($b = -0.01$, 95% CI [–0.03, 0.00], $p = 0.064$; Fig. S15h).

#### Exploratory analyses on cross-cultural differences

This section presents exploratory analyses of potential cross-cultural differences between Eastern and Western participants in our study. It is considered “exploratory” because of the following limitations. First, because Experiment 2's primary objective was to replicate Experiment 1's main findings, exploring cultural differences was beyond our pre-registered scope. Second and relatedly, because we did not initially plan to study specific cultural backgrounds, we did not control for variables like sample size from different cultures, overseas experience, or immigration status. As a result, the sample sizes of different cultural groups were imbalanced; for different groups, the proportions of participants from the on-site Experiment 1 and the online Experiment 2 were also imbalanced. These factors should be kept in mind when interpreting the results of these exploratory analyses.

Despite these limitations, we recognized the value in exploring the cultural aspects of our data. We performed exploratory analyses by categorizing participants from both Experiments 1 and 2 into Eastern and Western cultural backgrounds. To ensure comparable decision-making processes across groups, we first excluded participants whose choice behaviors were best described by the simple-response model (Model 9 in the main text) that linearly combines different independent variables (see Methods and Fig. S9b). This step was necessary because the proportions of simple-response participants differed substantially between the on-site Experiment 1 (0%) and online Experiment 2 (39.11%).

We then categorized participants into Eastern and Western groups based on their countries of origin (Markus & Kitayama, 1991). To minimize the confounding effects of individuals living in different cultural areas, we excluded participants whose records spanned both Eastern and Western regions in terms of nationality, country of birth, or country of residence. For example, a participant born in China but currently holding Western nationality or living in a Western country would be excluded from this analysis. After these exclusions, our final sample consisted of 158 participants in the East group (all from Experiment 1, except for one participant) and 355 participants in the West group (all from Experiment 2). This imbalance in group sizes and experiment representation should be considered when interpreting the results. See Fig. S20 for the comparison of behavioral patterns between East and West groups, and Table S10 for detailed country distributions.

**West group exhibited higher reversal preference and self-centered disadvantageous inequality aversion than East group**

To examine what motive parameter may differ between the East and West groups, we performed Mann-Whitney *U* tests on each motive parameter estimated from Model 8, with Bonferroni correction for the seven comparisons. We found that compared to participants from the East group, participants in the West group had a higher reversal preference (κ: Z = 6.02, $p < 0.001$) and a higher self-centered disadvantageous inequality aversion (α: $Z$ = 2.88, $p$ = 0.028). Other motive parameters showed no significant difference between groups (β: $Z$ = 2.32, $p$ = 0.142; γ: Z = –0.50, $p > 0.999$; ω: $Z$ = 2.59, $p$ = 0.068; $\eta_{no}$: $Z$ = 1.24, $p$ = $p > 0.999$; $\eta_{yes}$: $Z$ = -2.06, $p$ = 0.277; see Fig. S21).

**Justice warriors, pragmatic helpers, and rational moralists in the East and West groups**

We also compared whether the frequencies of justice warriors, pragmatic helpers, and rational moralists differed between the East and West groups. According to chi-square test of independence, the relative frequencies of the three clusters were significantly different between the two groups ($\chi^2(2) = 7.92, p = 0.019$). Following proportion difference test with Bonferroni correction, the proportion of pragmatic helpers was higher in the West group ($Z$ = 2.77, $p$ = 0.017), while the proportions of justice warriors ($Z$ = –1.94, $p$ = 0.053) and rational moralists ($Z$ = –0.44, $p$ = 0.657) did not show significant differences between the two groups (Table S11). Recall that pragmatic helpers had the highest parameter of reverse preference, $\kappa$ (Figs. 4 & 5). The higher proportion of pragmatic helpers in the West group thus echoes the higher $\kappa$ in the West group.

#### Supplementary Tables

**Table S1.** Statistical results of GLMM1

| Fixed effects | Estimated beta value | SE | Z value | P value |
|---|---|---|---|---|
| (Intercept) | -3.61 | 0.28 | -12.81 | $p$ < 0.001 |
| Trial number | -0.20 | 0.05 | -3.79 | $p$ < 0.001 |
| Scenario | -1.22 | 0.21 | -5.71 | $p$ < 0.001 |
| Inequality | 1.61 | 0.11 | 15.08 | $p$ < 0.001 |
| Cost | -2.12 | 0.13 | -16.43 | $p$ < 0.001 |
| Ratio | 0.82 | 0.10 | 8.21 | $p$ < 0.001 |
| Trial number×Scenario | 0.07 | 0.05 | 1.36 | $p$ = 0.174 |
| Trial number×inequality | 0.02 | 0.03 | 0.60 | $p$ = 0.548 |
| Scenario×inequality | -0.03 | 0.05 | -0.67 | $p$ = 0.499 |
| Trial number×Cost | -0.04 | 0.03 | -1.26 | $p$ = 0.209 |
| Scenario×Cost | 0.00 | 0.05 | -1.33 | $p$ = 0.894 |
| Inequality×Cost | -0.09 | 0.03 | -2.92 | $p$ = 0.003 |
| Trial number×Ratio | -0.07 | 0.03 | -2.24 | $p$ = 0.003 |
| Scenario×Ratio | -0.39 | 0.05 | -8.65 | $p$ < 0.001 |
| Inequality×Ratio | -0.08 | 0.03 | -2.50 | $p$ = 0.012 |
| Cost×Ratio | -0.08 | 0.03 | -2.44 | $p$ = 0.015 |

| | | | | |
|---|---|---|---|---|
| Trial number×Scenario×Inequality | -0.08 | -0.05 | -1.17 | $p = 0.087$ |
| Trail number×Scenario×Cost | -0.01 | 0.05 | -0.21 | $p = 0.831$ |
| Trial number×Inequality×Cost | -0.01 | 0.05 | -0.23 | $p = 0.814$ |
| Scenario×Inequality×Cost | -0.01 | 0.03 | 0.18 | $p = 0.861$ |
| Trial number×Scenario×Ratio | -0.23 | 0.05 | 5.19 | $p < 0.001$ |
| Trial number×Inequality×Ratio | 0.01 | 0.03 | 0.22 | $p = 0.823$ |
| Scenario×Inequality×Ratio | 0.04 | 0.04 | 0.99 | $p = 0.322$ |
| Trial number×Cost×Ratio | -0.05 | 0.03 | -1.62 | $p = 0.106$ |
| Scenario×Cost×Ratio | -0.03 | 0.04 | -0.77 | $p = 0.441$ |
| Inequality×Cost×Ratio | -0.21 | 0.03 | -6.97 | $p < 0.001$ |
| Trial number×Scenario×Inequality×Cost | -0.04 | 0.04 | -0.88 | $p = 0.380$ |
| Trial number×Scenario×Inequality×Ratio | -0.05 | 0.04 | -1.25 | $p = 0.210$ |
| Trial number×Scenario×Cost×Ratio | 0.02 | 0.04 | 0.42 | $p = 0.673$ |
| Trial number×Inequality×Cost×Ratio | 0.06 | 0.03 | -1.90 | $p = 0.057$ |
| Scenario×Inequality×Cost×Ratio | 0.05 | 0.04 | 1.21 | $p = 0.226$ |
| Trial number×Scenario×Inequality×Cost×Ratio | 0.04 | 0.04 | 0.93 | $p = 0.352$ |

**Table S2.** Fictitious examples to illustrate the motives in the motive cocktail model.

| Motive | Fictitious example |
|---|---|
| **Self-interest** | Alice allocates resources, keeping a larger share for herself and giving a smaller portion to Bob. Charlie, a third party, observes this unequal distribution between Alice and Bob but chooses not to intervene due to the personal cost involved in taking action. |
| **Self-centered inequality aversion** | **Disadvantageous inequality aversion**: Alice allocates resources, keeping more for herself than she gives to Bob. Charlie, observing this, punishes Alice to ensure that Alice does not end up with more than Charlie himself. In this case, Charlie acts to minimize his own disadvantageous inequality relative to Alice.<br><br>**Advantageous inequality aversion**: Alice allocates resources, keeping more for herself than she gives to Bob. Charlie, observing this, helps Bob to ensure that Bob does not end up with less than Charlie himself. In this case, Charlie acts to minimize his own advantageous inequality relative to Bob. |
| **Victim-centered inequality aversion** | Alice allocates resources, keeping more for herself than she gives to Bob. Charlie, observing this unequal distribution, intervenes by either punishing Alice or helping Bob, with the goal of equalizing their final outcomes. In this case, Charlie acts to minimize the disadvantageous inequality experienced by the victim, Bob. |
| **Efficiency concern** | Alice allocates resources, keeping more for herself than she gives to Bob. Charlie, observing this unequal distribution, chooses to help Bob rather than punish Alice. This action ensures that the total sum of resources for Alice and Bob increases. In this case, Charlie acts to maximize the overall payoff for others. |
| **Reversal preference** | **Reversal aversion**: Alice allocates resources, keeping more for herself than she gives to Bob. Charlie has an opportunity to intervene by either punishing Alice or helping Bob. However, Charlie realizes that such intervention would result in Bob having more than Alice. Charlie decides not to intervene, demonstrating reversal aversion—a preference |

| | |
|---|---|
| | to avoid reversing the original inequality.<br><br>**Reversal preference**: In the same scenario, where Alice keeps more for herself than she gives to Bob, Charlie has the opportunity to intervene. Despite recognizing that intervention would result in Bob having more than Alice, Charlie chooses to intervene anyway. This demonstrates reversal preference - a willingness to create reverse inequality in the process of addressing the original imbalance. |
| **Inaction inequality discounting** | Alice allocates resources, keeping more for herself than she gives to Bob. Charlie observes this unequal distribution and has an opportunity to intervene by either punishing Alice or helping Bob. However, recognizing that such intervention would be costly for himself, Charlie chooses to disregard the disadvantageous inequality Bob is experiencing. Charlie acts as if he cannot see the inequality and decides not to intervene, effectively discounting the observed inequality to justify his inaction. |
| **Action inequality discounting** | Alice allocates resources, keeping more for herself than she gives to Bob. Charlie observes this unequal distribution and has an opportunity to intervene by either punishing Alice or helping Bob. Although Charlie recognizes that his intervention would only slightly reduce the disadvantageous inequality Bob is experiencing, and that Bob would still end up with less than Alice, he decides to take action anyway. Charlie justifies his intervention by discounting the remaining inequality, believing that his effort balances out the persisting disparity. |

**Table S3.** Real-life examples to illustrate the motives in the motive cocktail model.

| Motive | Real-life example |
|---|---|
| **Self-interest** | **Scenario:** In a community garden, volunteers are needed to help with a variety of tasks such as weeding, planting, and watering.<br>**Example:** A community member notices that the garden needs attention and that there's a sign-up sheet for volunteers. Despite having free time, they decide not to sign up or participate, preferring to use their leisure time for personal activities rather than contributing to the community project. |
| **Self-centered inequality aversion** | **Scenario:** A company is distributing annual bonuses to its employees based on performance.<br>**Example 1 (disadvantageous inequality aversion):** An employee learns that their colleague in the same role received a larger bonus. The employee appeals to management for a bonus increase to ensure they don't earn less than their peer.<br><br>**Example 2 (advantageous inequality aversion):** A team leader discovers they received a significantly larger bonus than their team members, despite similar contributions. The leader advocates for their team members to receive larger bonuses to reduce their own discomfort with having a much higher bonus than their peers. |
| **Victim-centered inequality aversion** | **Scenario:** In a small office, the manager consistently assigns the most desirable projects and clients to one team member, Mark, while giving less appealing tasks to another team member, Sarah.<br><br>**Example:** A third team member, Lisa, observes this pattern of unequal distribution of work. Despite not being directly affected, Lisa decides to intervene. She speaks to the manager, ad<br>vocating for a more balanced distribution of projects. Lisa suggests either reassigning some of Mark's high-profile projects to Sarah or providing Sarah with additional resources and support to enhance her current projects. Lisa's primary motivation is to reduce the disadvantage experienced by Sarah, aiming to equalize opportunities and recognition between Mark and Sarah. |
| **Efficiency concern** | **Scenario:** During a community park cleanup event, volunteer |

| | |
|---|---|
| | Alex is actively picking up litter, while volunteer Sam is merely observing and giving occasional directions.<br><br>**Example:** A third volunteer, Jordan, notices this situation. Instead of confronting Sam about their lack of hands-on participation, Jordan chooses to assist Alex in collecting trash. Jordan's decision is motivated by the desire to maximize the overall amount of litter removed from the park. By focusing on increasing the total output of the cleanup effort rather than ensuring equal participation, Jordan prioritizes the efficiency and overall impact of the group's work. |
| **Reversal preference** | **Scenario:** In a small tech startup, the CEO has allocated a limited budget for employee bonuses. The senior developer, Tom, receives a significantly larger bonus than the junior developer, Emily, despite Emily having contributed crucial work to a recent successful project.<br><br>**Example 1 (reversal aversion):** The HR manager, Sarah, notices this disparity and has the authority to adjust the bonuses. She considers redistributing some of Tom's bonus to Emily or advocating for an increase in Emily's bonus. However, Sarah calculates that any meaningful adjustment would result in Emily's total compensation (base salary plus bonus) exceeding Tom's. Concerned about creating a reversed inequality where the junior developer earns more than the senior developer, Sarah decides not to intervene, demonstrating reversal aversion.<br><br>**Example 2 (reversal preference):** In the same situation, the HR manager, Mike, also notices the bonus disparity between Tom and Emily. Mike recognizes that adjusting the bonuses would likely result in Emily's total compensation surpassing Tom's. Despite this, Mike decides to intervene by recommending a significant increase to Emily's bonus, acknowledging her crucial contributions to the recent project. This action demonstrates Mike's reversal preference, as he is willing to create a reversed inequality to address the original imbalance and recognize Emily's performance. |
| **Inaction inequality discounting** | **Scenario:** In a sports team, the coach favors certain players over others, giving them more playtime.<br><br>**Example:** A teammate notices the favoritism but chooses not |

| | |
|---|---|
| | to speak up because challenging the coach could cost them their own playtime or position, thus ignoring the inequality. |
| **Action inequality discounting** | **Scenario:** In a volunteer group, one volunteer does most of the work but gets the same recognition as others.<br><br>**Example:** Another volunteer decides to speak up and advocate for more recognition for the hard-working volunteer, even though the overall recognition still remains somewhat unequal, believing their effort will partially balance the inequality. |

**Table S4.** The nationality of participants in Experiment 2.

| Nationality | N | Percentage | Nationality | N | Percentage |
|---|---|---|---|---|---|
| South Africa | 193 | 15.34% | Australia | 3 | 0.24% |
| Italy | 164 | 13.04% | Pakistan | 2 | 0.16% |
| Mexico | 141 | 11.21% | Moldova | 2 | 0.16% |
| Poland | 132 | 10.49% | Egypt | 2 | 0.16% |
| Portugal | 118 | 9.38% | Norway | 2 | 0.16% |
| Greece | 72 | 5.72% | Vietnam | 2 | 0.16% |
| Spain | 69 | 5.48% | Venezuela, Bolivarian Republic of | 2 | 0.16% |
| Chile | 40 | 3.18% | New Zealand | 2 | 0.16% |
| Hungary | 30 | 2.38% | Indonesia | 2 | 0.16% |
| Germany | 28 | 2.23% | Uganda | 2 | 0.16% |
| United Kingdom | 26 | 2.07% | Sri Lanka | 1 | 0.08% |
| Canada | 26 | 2.07% | Namibia | 1 | 0.08% |
| France | 17 | 1.35% | Lebanon | 1 | 0.08% |
| Netherlands | 16 | 1.27% | Cameroon | 1 | 0.08% |
| Czech Republic | 15 | 1.19% | Nepal | 1 | 0.08% |

| | | | | | |
|---|---|---|---|---|---|
| United States | 14 | 1.11% | India | 1 | 0.08% |
| Slovenia | 13 | 1.03% | Bosnia and Herzegovina | 1 | 0.08% |
| Latvia | 10 | 0.79% | Croatia | 1 | 0.08% |
| Estonia | 10 | 0.79% | Switzerland | 1 | 0.08% |
| Belgium | 9 | 0.72% | Peru | 1 | 0.08% |
| Ireland | 8 | 0.64% | Bulgaria | 1 | 0.08% |
| Austria | 7 | 0.56% | Singapore | 1 | 0.08% |
| Brazil | 6 | 0.48% | Ghana | 1 | 0.08% |
| Zimbabwe | 6 | 0.48% | Argentina | 1 | 0.08% |
| Turkey | 6 | 0.48% | Algeria | 1 | 0.08% |
| Finland | 6 | 0.48% | Lesotho | 1 | 0.08% |
| Sweden | 5 | 0.40% | Bangladesh | 1 | 0.08% |
| Iran | 5 | 0.40% | Morocco | 1 | 0.08% |
| Philippines | 5 | 0.40% | Suriname | 1 | 0.08% |
| Nigeria | 4 | 0.32% | Zambia | 1 | 0.08% |
| Israel | 4 | 0.32% | Colombia | 1 | 0.08% |
| China | 4 | 0.32% | Saudi Arabia | 1 | 0.08% |
| Romania | 3 | 0.24% | | | |
| Ukraine | 3 | 0.24% | | | |

**Table S5.** Statistical results of GLMM2.

| Fixed effects | Estimated beta value | SE | Z value | P value |
|---|---|---|---|---|
| (Intercept) | -1.04 | 0.07 | -15.13 | $p$ < 0.001 |
| Trial number | -0.13 | 0.01 | -8.93 | $p$ < 0.001 |
| Scenario | -1.19 | 0.06 | -18.37 | $p$ < 0.001 |
| Inequality | 0.59 | 0.03 | 20.16 | $p$ < 0.001 |
| Cost | -0.78 | 0.03 | -25.47 | $p$ < 0.001 |
| Ratio | 0.71 | 0.03 | 25.19 | $p$ < 0.001 |
| Trial number×Scenario | -0.03 | 0.01 | -2.43 | $p$ = 0.015 |
| Trial number×inequality | 0.03 | 0.00 | 4.70 | $p$ < 0.001 |
| Scenario×inequality | 0.16 | 0.01 | 15.50 | $p$ < 0.001 |
| Trial number×Cost | -0.02 | 0.00 | -2.89 | $p$ < 0.004 |
| Scenario×Cost | -0.09 | 0.01 | -8.64 | $p$ < 0.001 |
| Inequality×Cost | -0.00 | 0.01 | -0.36 | $p$ = 0.716 |
| Trial number×Ratio | -0.04 | 0.01 | -5.63 | $p$ < 0.001 |
| Scenario×Ratio | -0.31 | 0.01 | -30.67 | $p$ < 0.001 |
| Inequality×Ratio | -0.01 | 0.01 | -1.85 | $p$ = 0.064 |
| Cost×Ratio | 0.04 | 0.01 | 5.46 | $p$ < 0.001 |
| Trial number×Scenario×Inequality | -0.03 | 0.01 | -2.87 | $p$ = 0.004 |

| | | | | |
|---|---|---|---|---|
| Trail number×Scenario×Cost | 0.03 | 0.01 | 2.63 | $p = 0.008$ |
| Trial number×Inequality×Cost | -0.02 | 0.01 | -2.71 | $p = 0.007$ |
| Scenario×Inequality×Cost | 0.02 | 0.01 | 2.12 | $p = 0.034$ |
| Trial number×Scenario×Ratio | 0.05 | 0.01 | 4.87 | $p < 0.001$ |
| Trial number×Inequality×Ratio | 0.01 | 0.01 | 1.55 | $p = 0.119$ |
| Scenario×Inequality×Ratio | 0.03 | 0.01 | 2.66 | $p = 0.008$ |
| Trial number×Cost×Ratio | -0.03 | 0.01 | -4.58 | $p < 0.001$ |
| Scenario×Cost×Ratio | -0.04 | 0.01 | -4.33 | $p < 0.001$ |
| Inequality×Cost×Ratio | -0.02 | 0.00 | -2.30 | $p = 0.021$ |
| Trial number×Scenario×Inequality×Cost | 0.02 | 0.01 | 2.07 | $p = 0.038$ |
| Trial number×Scenario×Inequality×Ratio | -0.02 | 0.01 | -2.09 | $p = 0.036$ |
| Trial number×Scenario×Cost×Ratio | 0.02 | 0.01 | 2.24 | $p = 0.025$ |
| Trial number×Inequality×Cost×Ratio | 0.02 | 0.01 | 1.80 | $p = 0.070$ |
| Scenario×Inequality×Cost×Ratio | -0.01 | 0.01 | -3.14 | $p = 0.002$ |
| Trial number×Scenario×Inequality×Cost×Ratio | 0.00 | 0.01 | -0.70 | $p = 0.485$ |

**Table S6.** Statistical results of LMM1.

| Fixed effects | Estimate | SE | df | t value | P value |
|---|---|---|---|---|---|
| (Intercept) | 896.94 | 45.75 | 163.00 | 19.61 | $p < 0.001$ |
| Trial number | -74.12 | 43.10 | 197.25 | -1.72 | $p = 0.087$ |
| Choice | 467.24 | 83.68 | 126.07 | 5.58 | $p < 0.001$ |
| Scenario | 4.76 | 35.25 | 166.78 | 0.14 | $p = 0.893$ |
| Inequality | 112.89 | 21.65 | 363.48 | 5.21 | $p < 0.001$ |
| Cost | -124.28 | 23.17 | 322.57 | -5.37 | $p < 0.001$ |
| Ratio | 35.26 | 16.49 | 2222.65 | 2.14 | $p = 0.032$ |
| Trial number×Choice | -214.71 | 45.39 | 36910.76 | -4.73 | $p < 0.001$ |
| Trial number×Scenario | -31.55 | 26.16 | 36066.02 | -1.21 | $p = 0.227$ |
| Choice×Scenario | 75.77 | 68.07 | 11285.37 | 1.11 | $p = 0.266$ |
| Trial number×Inequality | -3.76 | 16.72 | 35212.74 | -0.23 | $p = 0.822$ |
| Choice×Inequality | -223.81 | 41.93 | 18985.66 | -5.34 | $p < 0.001$ |
| Scenario×Inequality | 6.71 | 22.09 | 44119.63 | 0.30 | $p = 0.761$ |
| Trial number×Cost | -24.60 | 17.13 | 32627.23 | -1.44 | $p = 0.151$ |
| Choice×Cost | 382.64 | 41.56 | 16281.44 | 9.21 | $p < 0.001$ |
| Scenario×Cost | 17.15 | 22.40 | 44094.58 | 0.77 | $p = 0.444$ |
| Inequality×Cost | -39.82 | 16.43 | 41892.31 | -2.42 | $p = 0.015$ |
| Trial number×Ratio | -5.07 | 16.44 | 35165.99 | -0.31 | $p = 0.758$ |

| | | | | | |
|---|---|---|---|---|---|
| Choice×Ratio | -71.55 | 41.49 | 20079.08 | -1.72 | $p$ = 0.085 |
| Scenario×Ratio | -19.96 | 21.84 | 46036.13 | -0.91 | $p$ = 0.361 |
| Inequality×Ratio | 16.24 | 16.03 | 46063.51 | 1.01 | $p$ = 0.311 |
| Cost×Ratio | -33.50 | 16.28 | 46051.56 | -2.06 | $p$ = 0.039 |
| Trial number×Choice×Scenario | 129.11 | 68.37 | 32043.70 | 1.89 | $p$ = 0.058 |
| Trial number×Choice×Inequality | 29.44 | 41.25 | 40221.26 | 0.71 | $p$ = 0.475 |
| Trial number×Scenario×Inequality | -52.37 | 22.64 | 21527.93 | -2.31 | $p$ = 0.021 |
| Choice×Scenario×Inequality | -7.91 | 62.35 | 33869.84 | -0.13 | $p$ = 0.899 |
| Trial number×Choice×Cost | -18.81 | 39.38 | 40746.60 | -0.48 | $p$ = 0.632 |
| Trial number×Scenario×Cost | 58.78 | 23.38 | 18127.98 | 2.51 | $p$ = 0.012 |
| Choice×Scenario×Cost | -81.51 | 60.42 | 29263.29 | -1.35 | $p$ = 0.187 |
| Trial number×Inequality×Cost | -28.20 | 16.74 | 46405.25 | -1.69 | $p$ = 0.092 |
| Choice×Inequality×Cost | 106.58 | 38.56 | 34442.40 | 2.76 | $p$ = 0.005 |
| Scenario×Inequality×Cost | -1.21 | 22.34 | 46218.66 | -0.05 | $p$ = 0.957 |
| Trial number×Choice×Ratio | -62.39 | 41.01 | 41046.91 | -1.52 | $p$ = 0.128 |
| Trial number×Scenario×Ratio | 3.81 | 22.14 | 20674.98 | 0.17 | $p$ = 0.863 |
| Choice×Scenario×Ratio | -20.11 | 64.08 | 33313.35 | -0.31 | $p$ = 0.754 |
| Trial number×Inequality×Ratio | 12.00 | 16.41 | 46357.25 | 0.73 | $p$ = 0.464 |
| Choice×Inequality×Ratio | 3.03 | 39.81 | 41309.84 | 0.08 | $p$ = 0.939 |
| Scenario×Inequality×Ratio | 5.82 | 21.94 | 46387.69 | 0.27 | $p$ = 0.791 |

| | | | | | |
|---|---|---|---|---|---|
| Trial number×Cost×Ratio | -8.31 | 16.69 | 46380.40 | -0.50 | $p$ = 0.618 |
| Choice×Cost×Ratio | -33.88 | 38.22 | 40072.88 | -0.89 | $p$ = 0.375 |
| Scenario×Cost×Ratio | 15.63 | 22.23 | 46404.12 | 0.70 | $p$ = 0.482 |
| Inequality×Cost×Ratio | -28.87 | 16.27 | 46329.75 | -1.78 | $p$ = 0.076 |
| Trial number×Choice×Scenario×Inequality | 26.38 | 61.42 | 41519.65 | 0.43 | $p$ = 0.668 |
| Trial number×Choice×Scenario×Cost | 43.79 | 58.89 | 41220.79 | 0.74 | $p$ = 0.457 |
| Trial number×Choice×Inequality×Cost | 19.28 | 38.16 | 44991.68 | 0.51 | $p$ = 0.613 |
| Trial number×Scenario×Inequality×Cost | 39.06 | 22.54 | 46346.32 | 1.73 | $p$ =0.083 |
| Choice×Scenario×Inequality×Cost | 5.44 | 57.14 | 44983.34 | 0.10 | $p$ = 0.924 |
| Trial number×Choice×Scenario×Ratio | 66.46 | 62.57 | 43477.95 | 1.06 | $p$ = 0.288 |
| Trial number×Choice×Inequality×Ratio | 9.96 | 40.32 | 45794.41 | 0.25 | $p$ = 0.805 |
| Trial number×Scenario×Inequality×Ratio | -8.41 | 22.08 | 46340.55 | -0.38 | $p$ = 0.703 |
| Choice×Scenario×Inequality×Ratio | 40.75 | 61.14 | 45385.80 | 0.67 | $p$ = 0.505 |
| Trial number×Choice×Cost×Ratio | -59.07 | 38.08 | 45977.19 | -1.55 | $p$ = 0.121 |
| Trial number×Scenario×Cost×Ratio | 2.83 | 22.43 | 46364.57 | 0.13 | $p$ = 0.899 |
| Choice×Scenario×Cost×Ratio | 27.30 | 58.65 | 44765.80 | 0.47 | $p$ = 0.642 |
| Trial number×Inequality×Cost×Ratio | -23.03 | 16.67 | 46330.29 | -1.38 | $p$ = 0.167 |
| Choice×Inequality×Cost×Ratio | 65.75 | 37.47 | 45747.95 | 1.76 | $p$ = 0.079 |
| Scenario×Inequality×Cost×Ratio | 34.56 | 22.27 | 46351.37 | 1.55 | $p$ = 0.121 |
| Trial | -82.37 | 56.43 | 44287.02 | -1.46 | $p$ = 0.144 |

| | | | | | |
|---|---|---|---|---|---|
| number×Choice×Scenario×Inequality×Cost | | | | | |
| Trial number×Choice×Scenario×Inequality×Ratio | 27.35 | 60.35 | 45726.27 | 0.45 | *p* = 0.650 |
| Trial number×Choice×Scenario×Cost×Ratio | -28.45 | 57.30 | 45945.02 | -0.50 | *p* = 0.619 |
| Trial number×Choice×Inequality×Cost×Ratio | 70.88 | 37.86 | 46371.83 | 1.87 | *p* = 0.061 |
| Trial number×Scenario×Inequality×Cost×Ratio | 8.77 | 22.44 | 46326.13 | 0.39 | *p* = 0.696 |
| Choice×Scenario×Inequality×Cost×Ratio | -45.65 | 56.79 | 46362.43 | -0.80 | *p* = 0.421 |
| Trial number×Choice×Scenario×Inequality×Cost×Ratio | 2.87 | 55.99 | 46361.06 | 0.05 | *p* = 0.959 |

**Table S7.** Statistical results of LMM2.

| Fixed effects | Estimate | SE | df | t value | P value |
|---|---|---|---|---|---|
| (Intercept) | 1.61 | 0.09 | 1511 | 18.03 | $p < 0.001$ |
| Trial number | 0.15 | 0.06 | 13730 | 2.43 | $p = 0.015$ |
| Choice | -0.15 | 0.11 | 4271 | -1.32 | $p = 0.188$ |
| Scenario | -0.13 | 0.09 | 3146 | -1.44 | $p = 0.150$ |
| Inequality | -0.08 | 0.07 | 6831 | -1.14 | $p = 0.253$ |
| Cost | -0.14 | 0.06 | 21800 | -2.24 | $p = 0.025$ |
| Ratio | -0.03 | 0.06 | 13370 | -0.54 | $p = 0.593$ |
| Trial number×Choice | -0.06 | 0.09 | 212100 | -0.65 | $p = 0.519$ |
| Trial number×Scenario | -0.06 | 0.08 | 299900 | -0.71 | $p = 0.475$ |
| Choice×Scenario | 0.43 | 0.14 | 76670 | 3.03 | $p = 0.002$ |
| Trial number×Inequality | -0.11 | 0.06 | 373600 | -1.84 | $p = 0.066$ |
| Choice×Inequality | 0.12 | 0.09 | 232100 | 1.31 | $p = 0.191$ |
| Scenario×Inequality | 0.10 | 0.08 | 366100 | 1.24 | $p = 0.216$ |
| Trial number×Cost | -0.05 | 0.06 | 371400 | -0.93 | $p = 0.352$ |
| Choice×Cost | 0.20 | 0.09 | 203200 | 2.14 | $p = 0.032$ |
| Scenario×Cost | 0.08 | 0.08 | 357100 | 0.98 | $p = 0.325$ |
| Inequality×Cost | 0.03 | 0.06 | 366200 | 0.47 | $p = 0.636$ |
| Trial number×Ratio | -0.01 | 0.06 | 369100 | -0.24 | $p = 0.809$ |

| | | | | | |
|---|---|---|---|---|---|
| Choice×Ratio | 0.01 | 0.09 | 207500 | 0.11 | *p* = 0.916 |
| Scenario×Ratio | 0.03 | 0.08 | 353300 | 0.38 | *p* = 0.704 |
| Inequality×Ratio | 0.17 | 0.06 | 369700 | 2.97 | *p* = 0.003 |
| Cost×Ratio | 0.02 | 0.06 | 368700 | 0.35 | *p* = 0.729 |
| Trial number×Choice×Scenario | 0.17 | 0.14 | 313600 | 1.27 | *p* = 0.205 |
| Trial number×Choice×Inequality | 0.08 | 0.09 | 368900 | 0.87 | *p* = 0.382 |
| Trial number×Scenario×Inequality | 0.03 | 0.08 | 372200 | 0.42 | *p* = 0.673 |
| Choice×Scenario×Inequality | -0.29 | 0.14 | 296700 | -2.08 | *p* = 0.037 |
| Trial number×Choice×Cost | 0.01 | 0.09 | 361400 | 0.16 | *p* = 0.877 |
| Trial number×Scenario×Cost | 0.05 | 0.08 | 372500 | 0.66 | *p* = 0.511 |
| Choice×Scenario×Cost | 0.09 | 0.14 | 276200 | 0.68 | *p* = 0.499 |
| Trial number×Inequality×Cost | 0.02 | 0.06 | 375300 | 0.31 | *p* = 0.753 |
| Choice×Inequality×Cost | 0.08 | 0.09 | 358700 | 0.83 | *p* = 0.405 |
| Scenario×Inequality×Cost | -0.07 | 0.08 | 374900 | -0.91 | *p* = 0.366 |
| Trial number×Choice×Ratio | 0.01 | 0.09 | 353400 | 0.16 | *p* = 0.874 |
| Trial number×Scenario×Ratio | 0.01 | 0.08 | 372700 | 0.07 | *p* = 0.942 |
| Choice×Scenario×Ratio | -0.15 | 0.14 | 318000 | -1.09 | *p* = 0.275 |
| Trial number×Inequality×Ratio | 0.06 | 0.06 | 375300 | 1.04 | *p* = 0.300 |
| Choice×Inequality×Ratio | -0.17 | 0.09 | 355600 | -1.82 | *p* = 0.068 |
| Scenario×Inequality×Ratio | -0.15 | 0.08 | 374900 | -1.87 | *p* = 0.062 |

| | | | | | |
|---|---|---|---|---|---|
| Trial number×Cost×Ratio | 0.04 | 0.06 | 375300 | 0.73 | *p* = 0.467 |
| Choice×Cost×Ratio | -0.03 | 0.09 | 353300 | -0.30 | *p* = 0.762 |
| Scenario×Cost×Ratio | -0.03 | 0.08 | 374900 | -0.39 | *p* = 0.694 |
| Inequality×Cost×Ratio | -0.11 | 0.06 | 374900 | -1.87 | *p* = 0.061 |
| Trial number×Choice×Scenario×Inequality | -0.15 | 0.14 | 375100 | -1.12 | *p* = 0.263 |
| Trial number×Choice×Scenario×Cost | 0.08 | 0.13 | 371800 | 0.61 | *p* = 0.539 |
| Trial number×Choice×Inequality×Cost | 0.05 | 0.09 | 375400 | 0.57 | *p* = 0.571 |
| Trial number×Scenario×Inequality×Cost | -0.02 | 0.08 | 375400 | -0.20 | *p* = 0.844 |
| Choice×Scenario×Inequality×Cost | -0.05 | 0.13 | 372800 | -0.38 | *p* = 0.704 |
| Trial number×Choice×Scenario×Ratio | -0.10 | 0.14 | 372300 | -0.74 | *p* = 0.460 |
| Trial number×Choice×Inequality×Ratio | 0.01 | 0.09 | 375400 | 0.12 | *p* = 0.908 |
| Trial number×Scenario×Inequality×Ratio | -0.03 | 0.08 | 375300 | -0.33 | *p* = 0.744 |
| Choice×Scenario×Inequality×Ratio | 0.33 | 0.14 | 374300 | 2.45 | *p* = 0.014 |
| Trial number×Choice×Cost×Ratio | -0.02 | 0.09 | 375200 | -0.19 | *p* = 0.847 |
| Trial number×Scenario×Cost×Ratio | -0.09 | 0.08 | 375400 | -1.17 | *p* = 0.243 |
| Choice×Scenario×Cost×Ratio | -0.03 | 0.13 | 372900 | -0.24 | *p* = 0.810 |
| Trial number×Inequality×Cost×Ratio | -0.05 | 0.06 | 375300 | -0.88 | *p* = 0.377 |
| Choice×Inequality×Cost×Ratio | 0.07 | 0.09 | 375200 | 0.74 | *p* = 0.458 |
| Scenario×Inequality×Cost×Ratio | 0.10 | 0.08 | 375300 | 1.30 | *p* = 0.193 |
| Trial number×Choice×Scenario×Inequality×Cost | -0.11 | 0.13 | 375700 | -0.80 | *p* = 0.426 |

| | | | | | |
|---|---|---|---|---|---|
| Trial number×Choice×Scenario×Inequality×Ratio | 0.25 | 0.13 | 375900 | 1.82 | $p$ = 0.068 |
| Trial number×Choice×Scenario×Cost×Ratio | 0.06 | 0.13 | 375900 | 0.45 | $p$ = 0.652 |
| Trial number×Choice×Inequality×Cost×Ratio | 0.02 | 0.09 | 375400 | 0.26 | $p$ = 0.797 |
| Trial number×Scenario×Inequality×Cost×Ratio | -0.01 | 0.08 | 375300 | -0.08 | $p$ = 0.935 |
| Choice×Scenario×Inequality×Cost×Ratio | -0.01 | 0.13 | 375900 | -0.08 | $p$ = 0.937 |
| Trial number×Choice×Scenario×Inequality×Cost×Ratio | 0.10 | 0.13 | 375900 | 0.73 | $p$ = 0.466 |

**Table S8.** The proportions of participants who never chose to punish, who never chose to help, and who never chose to punish and help.

| | **% Never punish** | **% Never help** | **% Never punish and help** |
|---|---|---|---|
| Experiment 1 (n = 157) | 17.83% | 8.28% | 7.64% |
| Experiment 2 (n = 1258) | 9.14% | 3.66% | 2.78% |
| Experiment 2 (simple-response participants excluded, n = 766) | 12.53% | 6.01% | 4.57% |

**Table S9.** The proportions of justice warriors, pragmatic helpers, and rational moralists who never chose to punish, who never chose to help, and who never chose to punish and help.

| | | % Never punish | % Never help | % Never punish and help |
|---|---|---|---|---|
| | Justice warriors | 0% | 0% | 0% |
| **Experiment 1** | Pragmatic helpers | 17.86% | 0% | 0% |
| | Rational moralists | 31.08% | 17.57% | 16.22% |
| | Justice warriors | 0% | 0% | 0% |
| **Experiment 2** | Pragmatic helpers | 6.88% | 0% | 0% |
| | Rational moralists | 23.82% | 13.53% | 10.29% |

**Table S10.** Participants' nationality distributions in the East and West groups.

| Culture | Nationality | Number of participants | Percentage |
|---|---|---|---|
| East | China | 157 | 30.60% |
| | Indonesia | 1 | 0.19% |
| West | Italy | 102 | 19.88% |
| | Portugal | 73 | 14.23% |
| | Greece | 54 | 10.53% |
| | Spain | 38 | 7.41% |
| | Germany | 19 | 3.70% |
| | France | 11 | 2.14% |
| | United Kingdom | 11 | 2.14% |
| | Netherlands | 10 | 1.95% |
| | Belgium | 7 | 1.36% |
| | United States | 6 | 1.17% |
| | Canada | 5 | 0.97% |
| | Austria | 5 | 0.97% |
| | Ireland | 4 | 0.78% |
| | Sweden | 4 | 0.78% |
| | Australia | 3 | 0.58% |
| | Finland | 2 | 0.39% |
| | Turkey | 1 | 0.19% |

**Table S11.** Participants were clustered as justice warriors, pragmatic helpers, and rational moralists in both Eastern and Western cultures.

| Group | Cluster | N | Percentage |
|---|---|---|---|
| East | Justice warriors | 55 | 34.81% |
| | Pragmatic helpers | 28 | 17.72% |
| | Rational moralists | 75 | 47.47% |
| West | Justice warriors | 93 | 26.20% |
| | Pragmatic helpers | 101 | 28.45% |
| | Rational moralists | 161 | 45.35% |

#### Supplementary Figures

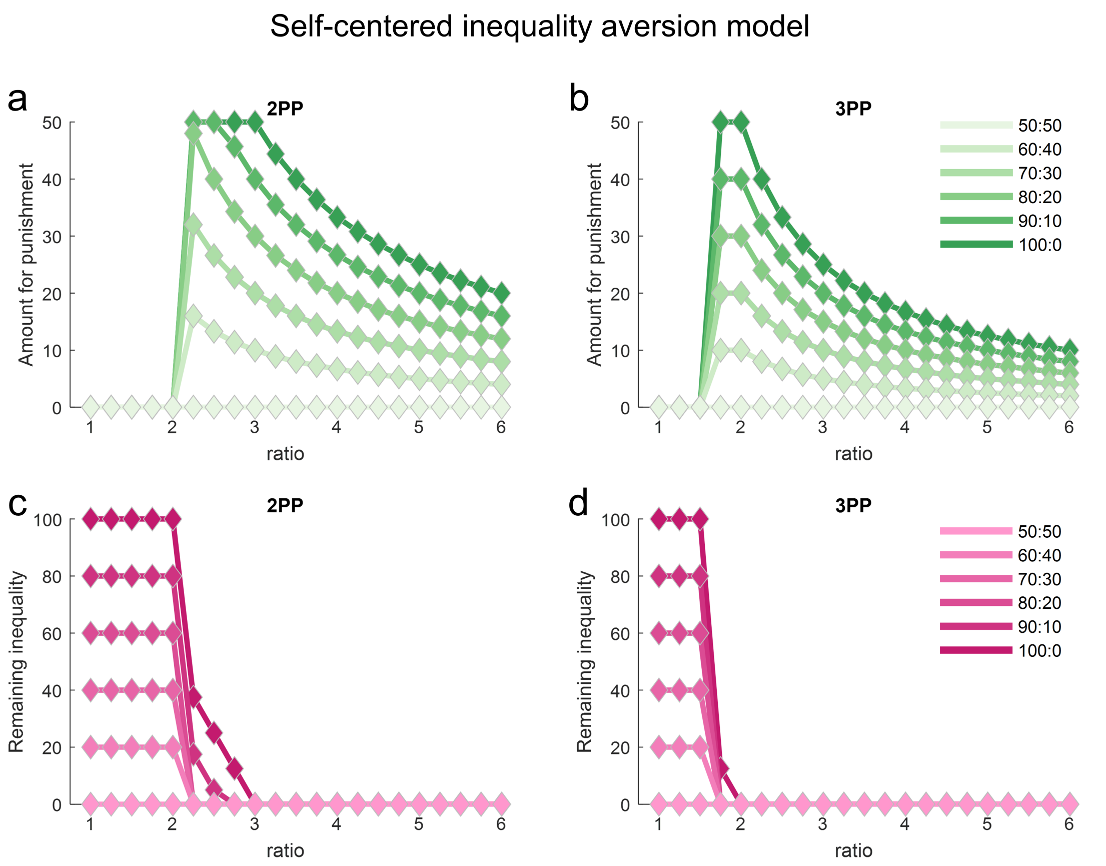


**Fig. S1 | Simulation of the behavior of an agent following the self-centered inequality aversion model. a–b**, The punishment amount as a function of the impact ratio and the inequality level between the transgressor and the victim in (**a**) second-party punishment ("2PP") and (**b**) third-party punishment ("3PP"). **c–d,** The remaining inequality after punishment, calculated by max ($x_1'$ - $x_2'$, 0), as a function of impact ratio and inequality level in (**c**) 2PP and (**d**) 3PP. The color of the lines represents inequality levels, with darker colors indicating higher inequality and lighter colors indicating lower inequality between the transgressor and the victim. The x-axis represents the impact ratio (e.g., ratio = 2 indicates that the amount of punishment reduces the transgressor's resources by twice that amount). Note that the model predicts either no punishment or full punishment to restore equality, depending on whether the impact ratio of the punishment is below or above a certain threshold.

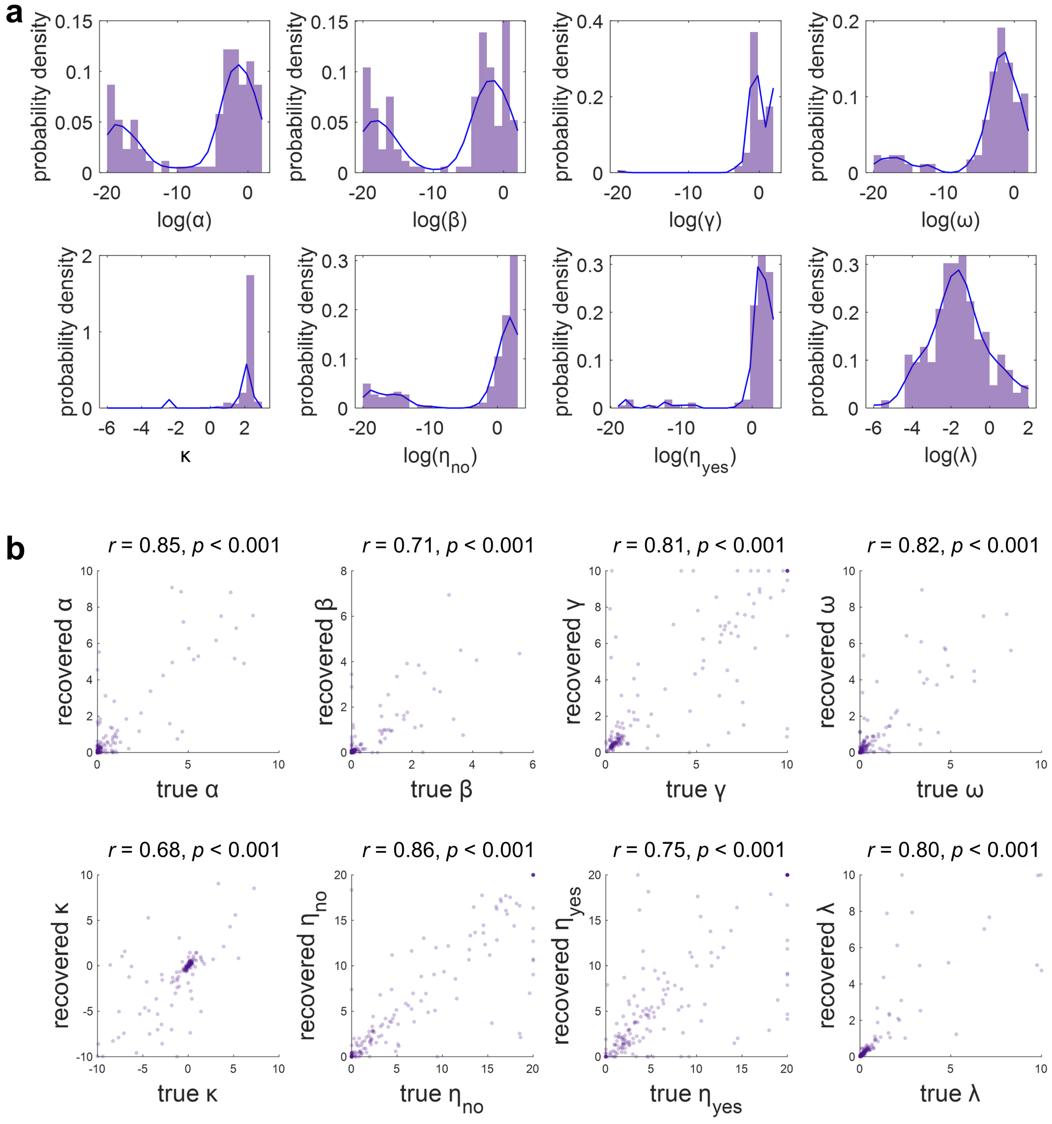


**Fig. S2 | The estimated parameters and parameter identifiability analysis for Experiment 1. a**, The distributions of the estimated parameters of the 157 participants for the motive cocktail model (model 7: SI+SCI+VCI+EC+RP+II). Each panel is for one parameter. The purple bars and blue curve respectively denote the histogram and its kernel fit. Except for $\kappa$, all parameters were transformed into log scale for better visualization. **b**, Results of parameter recovery for the motive cocktail model. The recovered parameters from 157 synthetic datasets are plotted against the generative parameters. Each panel is for one parameter. Each dot is for one virtual participant. The value of *r* indicates Pearson's correlation coefficient between the estimated and recovered parameters.

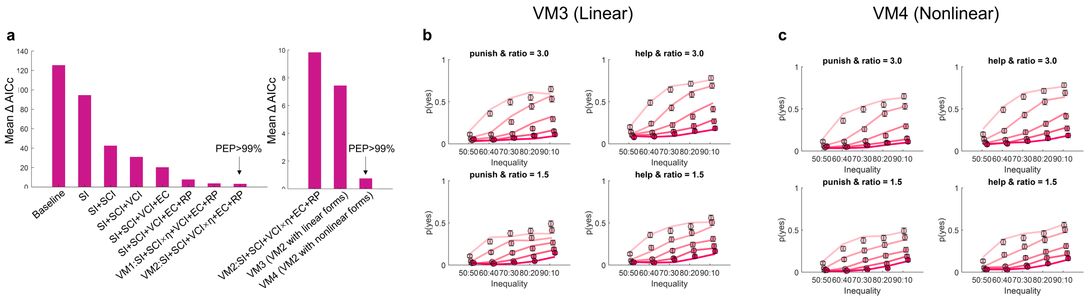


**Fig. S3 | Comparisons between models without and with inequality discounting assumption for Experiment 1.** (a) Model comparisons of models 1 to 6, along with VM1 and VM2. (a right panel) Model comparisons of VM2, VM3 and VM4. Model predictions of VM3 (b) and VM4 (c). For figures b&c, the probability of intervention, *p*(yes), is plotted against the inequality (from 50:50 to 90:10). Different colors code different levels of intervention cost (from 10 to 50, darker color for higher cost). Each sub-panel corresponds to one scenario and impact ratio condition. The dots and error bars respectively denote the mean and SEM across participants. The solid lines denote the predictions of the models.

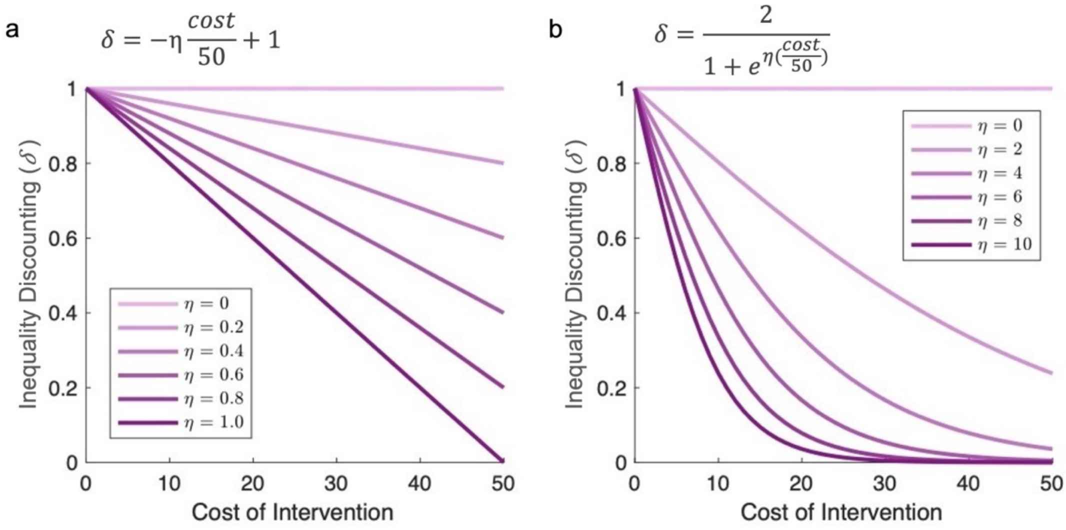


**Fig. S4 | Illustration of linear and non-linear inequality discounting functions. a,** Linear inequality discounting function (from the v3 model specified in the Supplementary Methods and Results), where inequality discounting is a linear function of the cost of intervention, $\delta = -\eta(cost/50) + 1$. **b,** Non-linear inequality discounting function (v4 model, same as Model 7 in the main text) $\delta = \frac{2}{1+e^{\eta(cost/50)}}$. The x-axis represents the cost of intervention. The y-axis represents the degree of inequality discounting, where smaller values indicate stronger discounting for the victim-centered disadvantageous inequality. The parameter $\eta$ controls the rate of discounting, with higher $\eta$ resulting in a faster discounting of inequality with the increases in intervention cost.

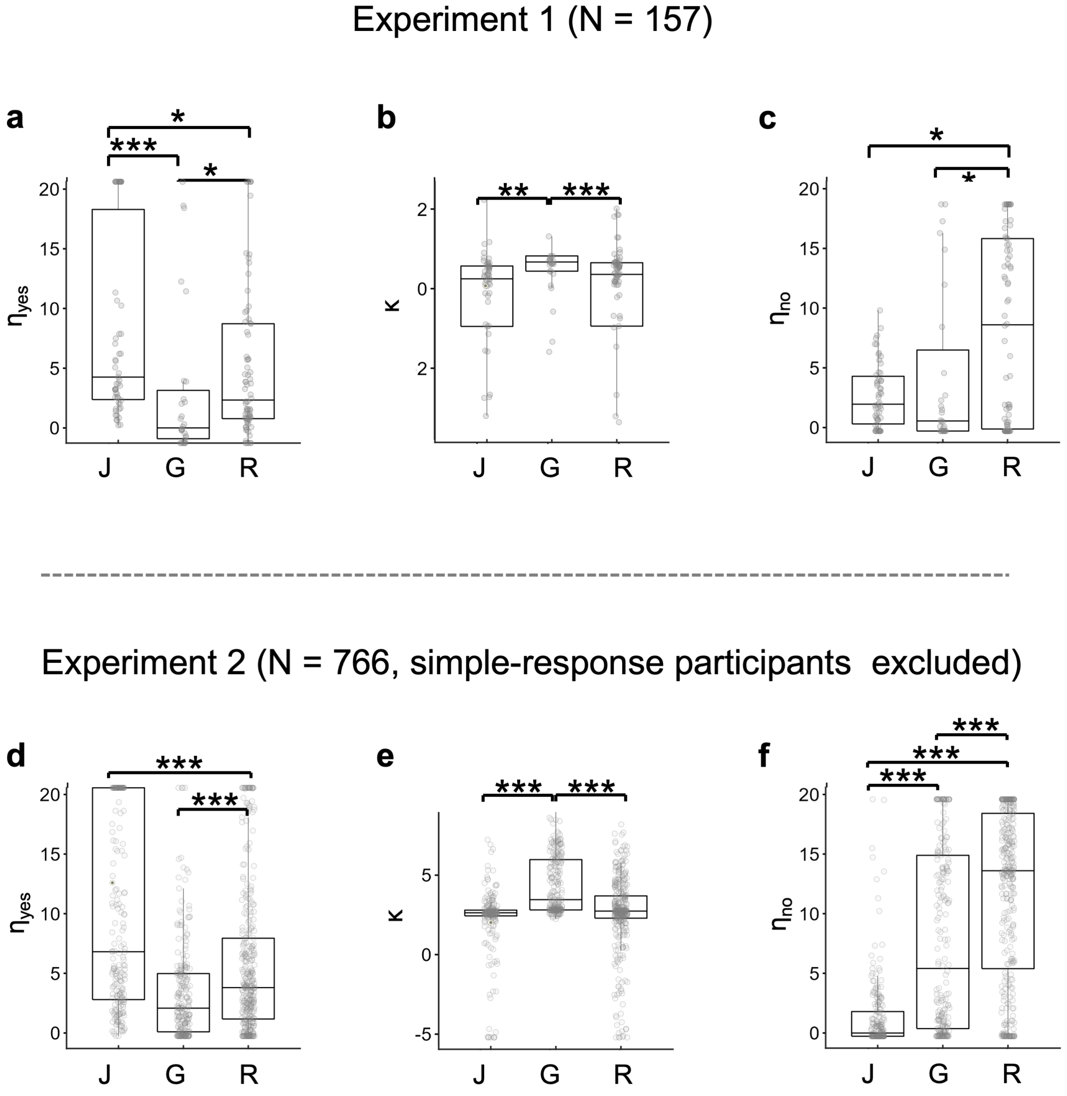


**Fig. S5 | Kruskal-Wallis test on parameter $\eta_{yes}$, $\kappa$ and $\eta_{no}$ across three clusters of participants.** For Experiment 2, participants who used simple-response model (i.e., best fit by Model 9) were excluded from analysis. The bottom/top and middle lines of the box plot represent the first/third quartiles and the median of the data. The lines extending beyond the box refer to 1.5 times the interquartile range (IQR), which is the distance between the third quartile (Q3) and the first quartile (Q1). ***, and *: $p < 0.001$ and $p < 0.05$ after multi-comparison corrections.

Experiment 1 (N = 157)

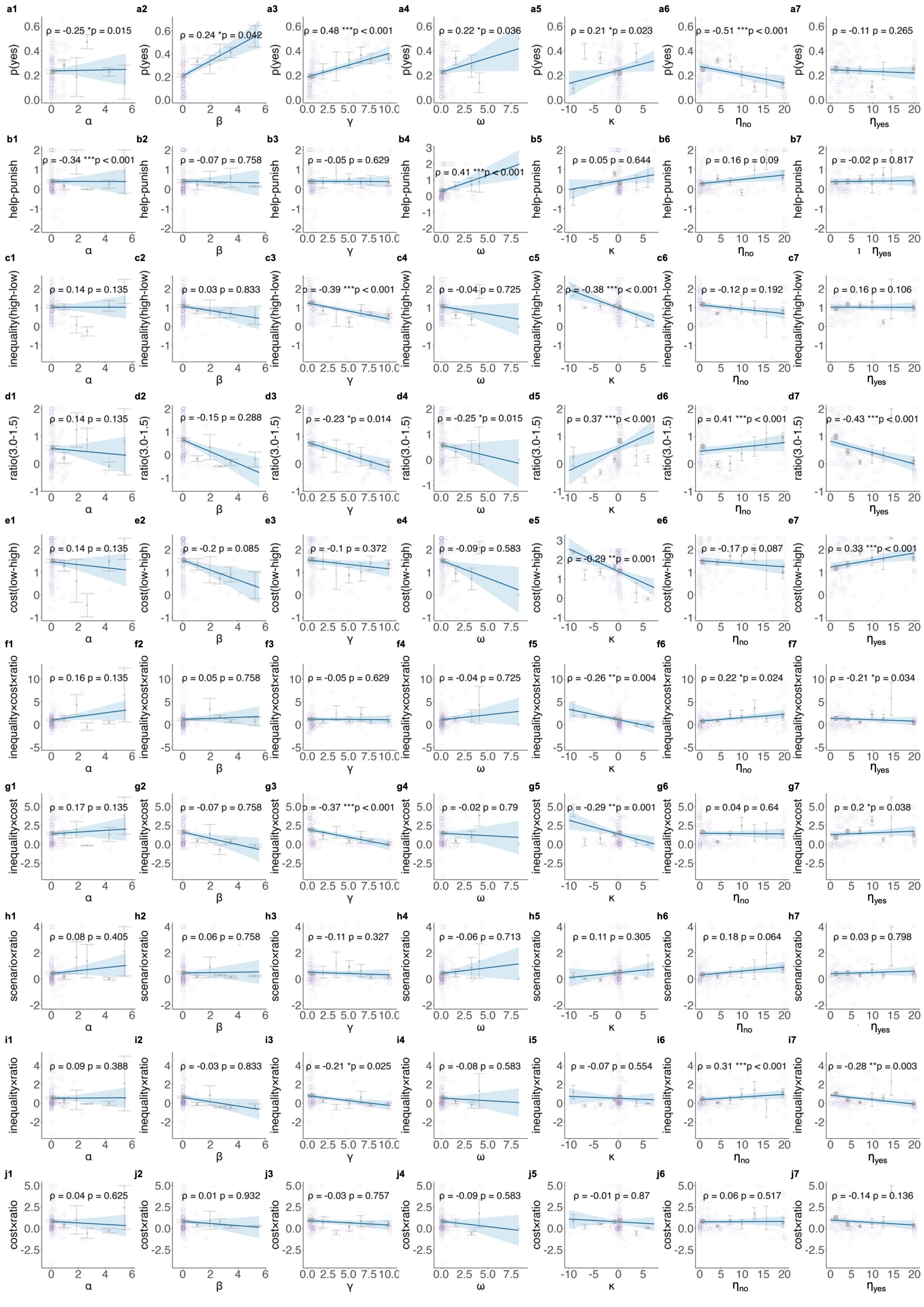


**Fig. S6 | The correlations between parameters estimated from the motive cocktail model and the model-free measurements for Experiment 1**. **a1 – a7,** The y-axis *p*(yes) represents the probability of intervention across all conditions for each participant. **b1 - b7**, The y-axis represents relative preference to help over punish, calculated as the probability of intervention in the helping scenario relative to that in the punishment scenario, normalized by the overall *p*(yes). **c1 - c7**, The y-axis represents the sensitivity to inequality, calculated as the probability of intervention in the high inequality trials (i.e., 70:30, 80:20,

90:10) relative to that in the low inequality trials (i.e., 50:50, 60:40), normalized by the overall *p*(yes). **d1 - d7**, The y-axis represents the sensitivity to ratio, calculated as the intervention probability difference between high impact ratio trials (ratio = 3.0) and low impact ratio trials (ratio = 1.5), normalized by the overall *p*(yes). **e1 - e7**, The y-axis represents the sensitivity to cost, calculated as the probability of intervention in low intervention cost trials (i.e., cost = 10, 20) minus that in high intervention cost trials (i.e., cost = 30, 40 and 50), normalized by the overall *p*(yes). **f1 - f7**, The y-axis represents the sensitivity to inequality under different levels of cost and ratio conditions, calculated as the normalized intervention probability difference in trials with different combinations of inequality, cost, and ratio: [(high ratio & high inequality & low cost - high ratio & high inequality & high cost)- (high ratio & low inequality & low cost - high ratio & low inequality & high cost)] minus [(low ratio & high inequality & low cost - low ratio & high inequality & high cost) - (low ratio & low inequality & low cost - low ratio & low inequality & high cost)]. **g1 - g7**, The y-axis represents the sensitivity to inequality under high-cost versus low-cost condition, calculated as the normalized intervention probability difference in trials with different combinations of inequality and cost: (high inequality & low cost – high inequality & high cost) - (low inequality & low cost – low inequality & high cost). **h1 - h7**, The y-axis represents the relative preference to help over punish under high versus low ratio conditions, calculated as the normalized intervention probability difference in trials with different combinations of scenario and ratio: (help & high ratio – help & low ratio) minus (punish & high ratio – punish & low ratio). **i1 - i7**, The y-axis represents the sensitivity to inequality under high versus low ratio conditions, calculated as the normalized intervention probability difference in trials with different combinations of inequality and ratio: (high inequality & high ratio – high inequality & low ratio) minus (low inequality & high ratio – low inequality & low ratio). **j1 - j1**, The y-axis represent the sensitivity to cost in high versus low ratio conditions, calculated as the normalized intervention probability difference in trials with different combination of cost and ratio: (low cost & high ratio – low cost & low ratio) minus (high cost & high ratio – high cost & low ratio). The x-axis for each column corresponds to one motive parameter of the motive cocktail model. Each panel illustrates the relationship between a motive parameter and a behavioral measure, with the x-axis divided into 8 bins across participants, and the y-axis displaying the mean (points) and standard deviation (error bars) within the corresponding bin. Each light-colored circle represents data from an individual participant. The blue line in each plot represents a linear regression between the original x and y coordinates (i.e., no bins), while the shaded area indicates the 95% confidence interval. The $\rho$ denotes partial correlation coefficient after controlling all other parameters. The *p* value was corrected for multiple comparisons using FDR.

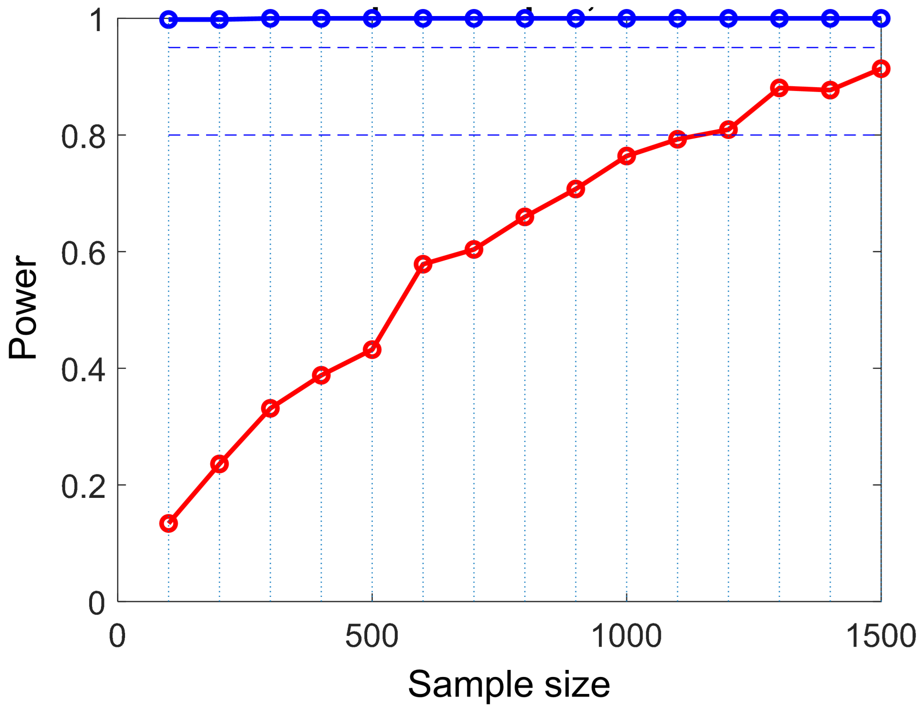


**Fig. S7 | Power analysis to determine the sample size in Experiment 2**. We used a parametric simulation method derived from our best fitting model (model 8: motive cocktail model + lapse rate) to pre-determine the sample size in Experiment 2. The x-axis is the sample size. The y-axis denotes the power, which is defined as the percentage of significance for an effect across all synthetic datasets within a specific sample size. The red line denotes the power curve obtained from the three-way interaction of inequality × cost × ratio. The blue line is obtained from the two-way interaction of inequality × cost and acted as a sanity check to the red line. The simulation result indicates the 80% power criterion can be achieved with at least 1200 participants.

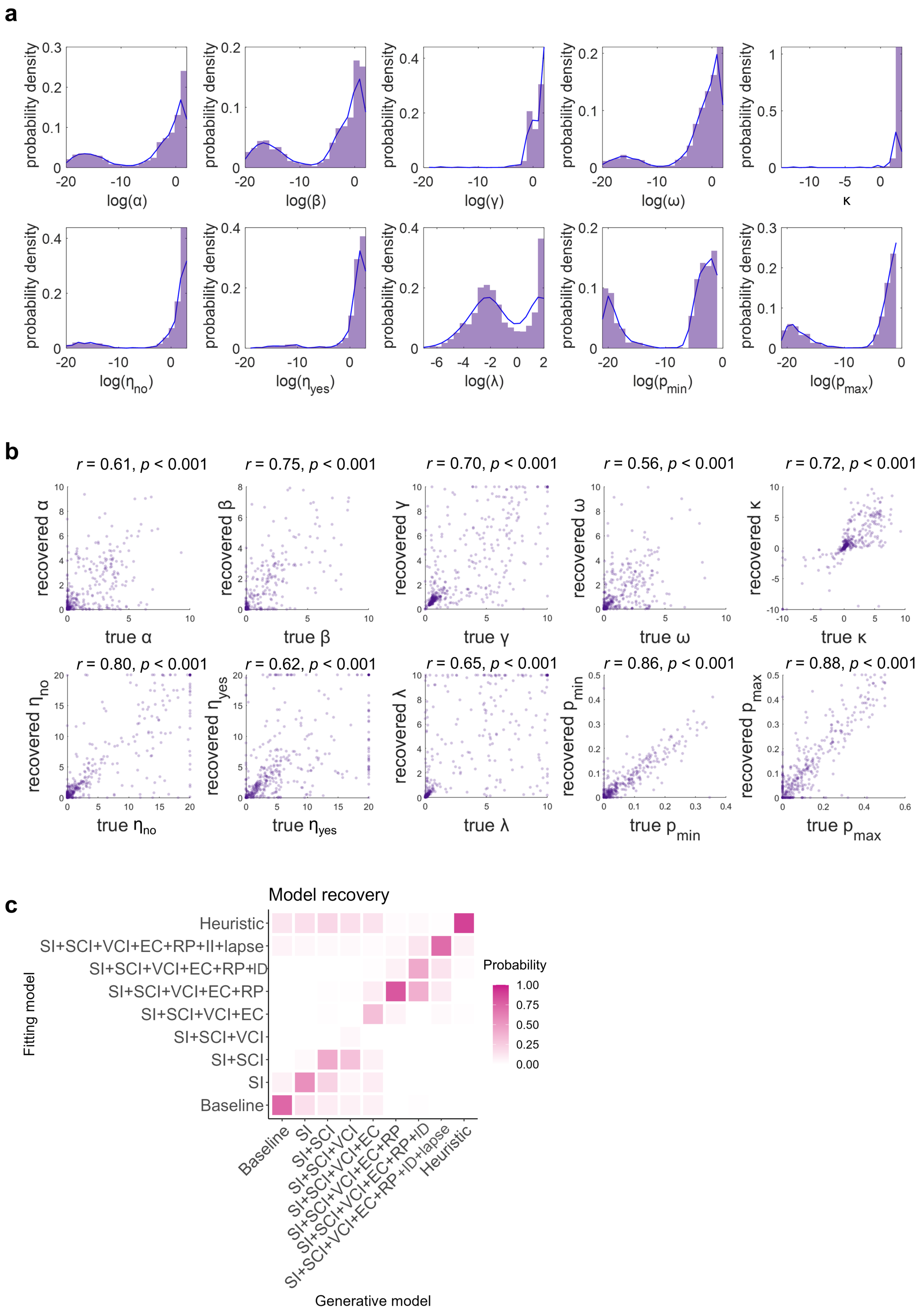


**Fig. S8 | Parameter identifiability and model recovery analyses for Experiment 2. a**, The true parameter distributions for the winning model (model 8: motive cocktail model + lapse). The histogram on each panel is plotted based on a specific parameter of 1258 participants. The blue line is an approximation for the parameter distribution across all participants fitted by a kernel-smoothing function. **b**, Parameter recovery for the full model (model 8). The recovered parameters from the synthetic datasets are plotted against the estimated parameters from the real data. Each panel is for one parameter. Each dot is for one virtual subject. The value of *r* indicates Pearson's correlation coefficient between the estimated and recovered parameters. **c**, Model recovery analysis. Each column and row are for one specific model that was used to generate synthetic datasets and fitted to the synthetic datasets respectively. The darker color in each cell represents a higher probability that the generative model can be best explained by a specific model.

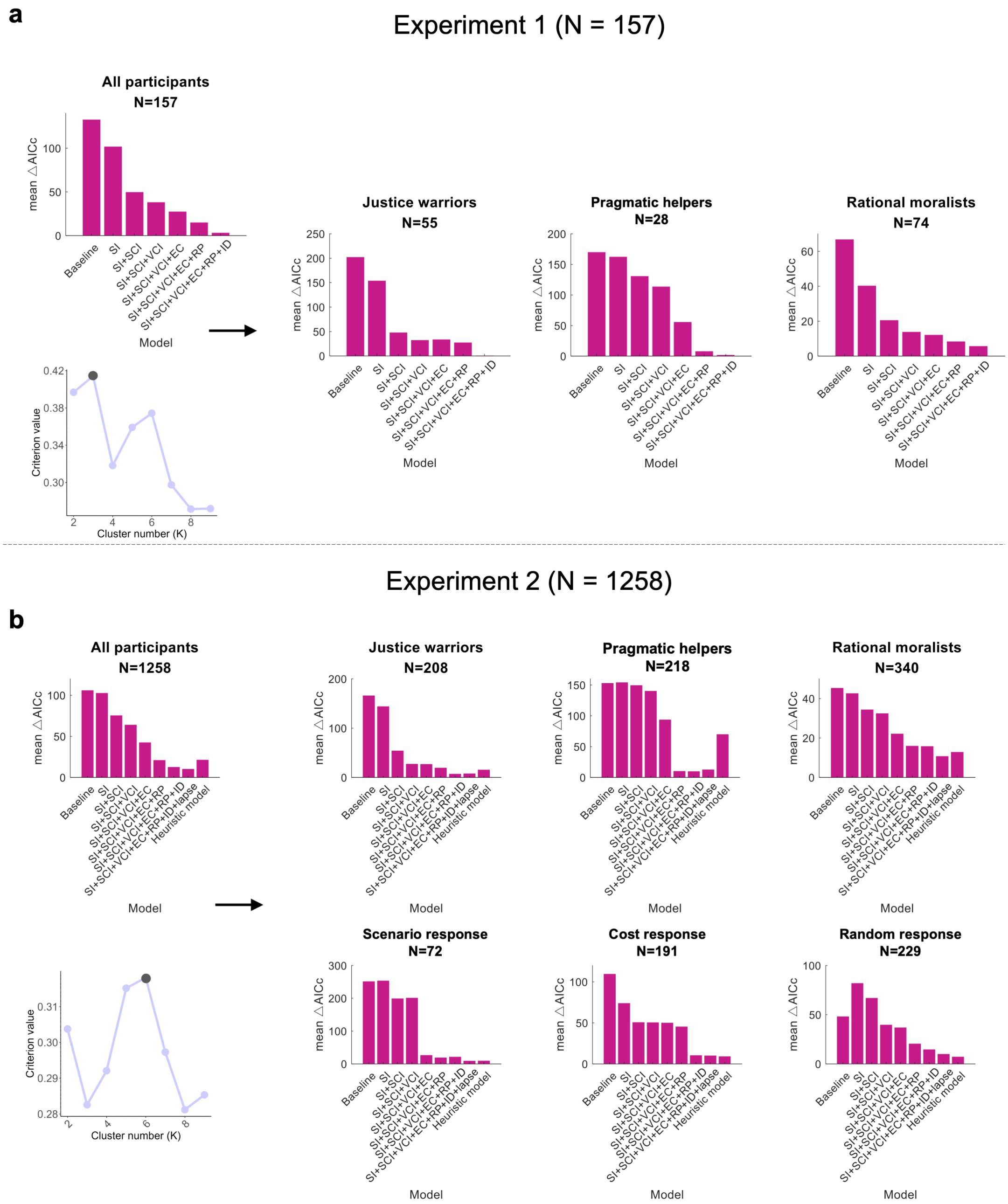


**Fig. S9 | Model comparisons for participants in each cluster**. **a**, Participants in Experiment 1 are best classified into 3 clusters, with all of them best accounted by the motive cocktail model. **b**, Participants in

Experiment 2 are best classified into 6 clusters, with the first 3 clusters best accounted for by the motive cocktail model (model 7 or its derivative model 8: motive cocktail model + lapse rates) and the remaining 3 clusters best by model 9 (simple-response model).

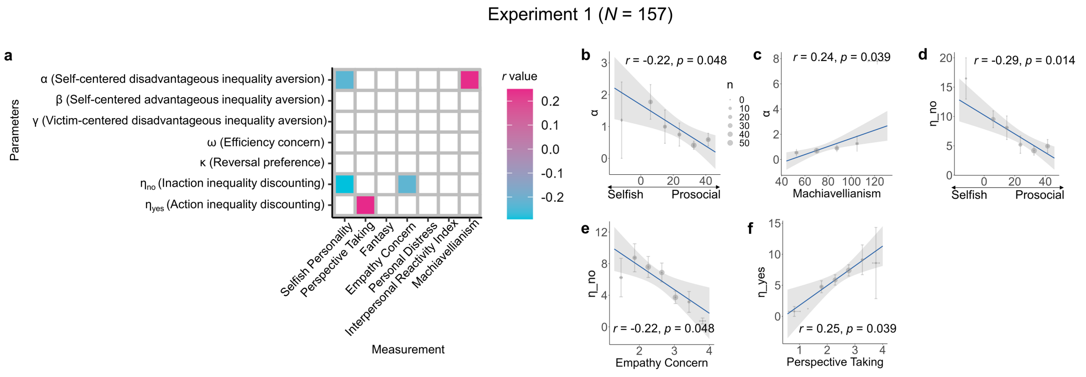


**Fig. S10 | The correlation between free parameters in the motive cocktail model and personality measurements in Experiment 1. a**, The correlation confusion matrix. For the personality measurement, the Selfish Personality was assessed by SVO (Murphy et al., 2011); Perspective Taking, Fantasy, Empathy Concern, Personal Distress and Interpersonal Reactivity Index were assessed by IRI (Davis, 1983); and Machiavellianism was assessed by MACH–IV (Rauthmann, 2013). Colored cells denote significant correlations, with warm and cool color representing the positive and negative correlation values. **b - f**, Regression plot for the six significant correlations. The *r* denotes Pearson's correlation coefficient. The *p* value was corrected for multiple comparisons using FDR.

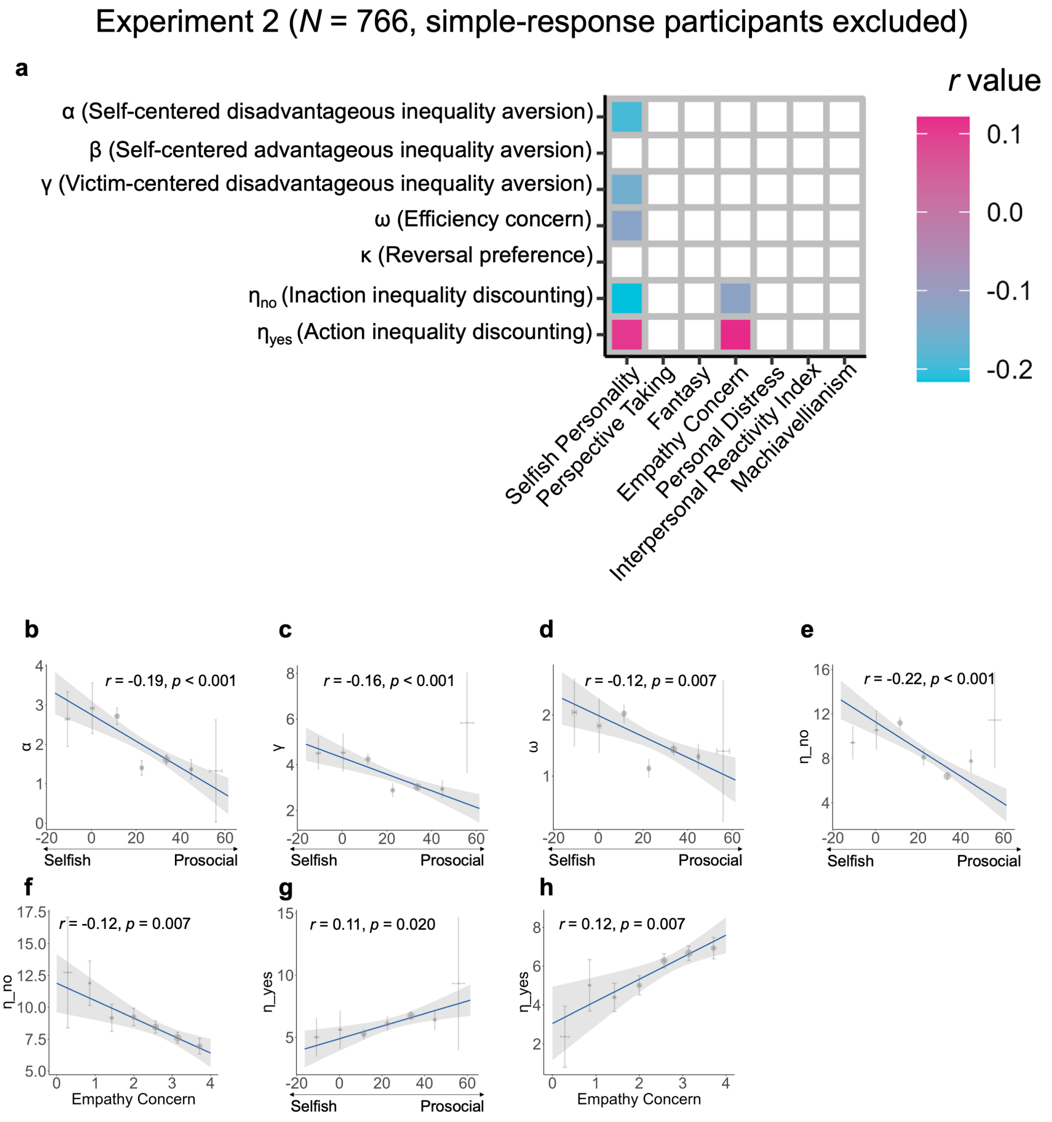


**Fig. S11 | The correlation between the estimated parameters from the motive cocktail model (with lapse rate; model 8) and personality measurements for justice warriors, pragmatic helpers and rational moralists in Experiment 2.** Participants who used simple-response model (i.e., best fit by Model 9) were excluded from analysis. **a**, The correlation confusion matrix. For the measurement, the Selfish Personality was assessed by SVO (Murphy et al., 2011); Perspective Taking, Fantasy, Empathy Concern, Personal Distress and Interpersonal Reactivity Index were assessed by IRI (Davis, 1983); and Machiavellianism was assessed by MACH–IV (Rauthmann, 2013). Colored cells denote significant correlations, with warm and cool color representing the positive and negative correlation values. **b**–**h**, Regression plot for the seven significant correlations. All results reported here are corrected by FDR.

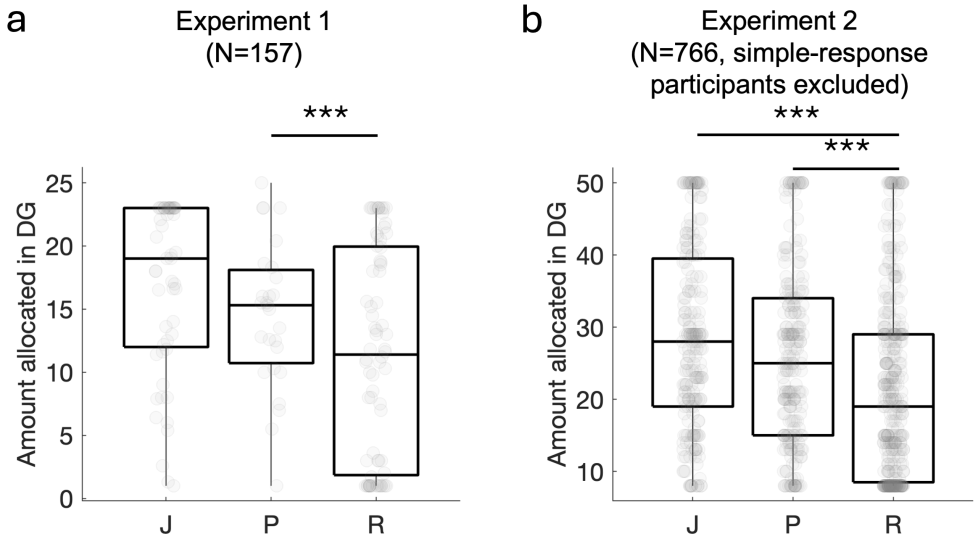


**Fig. S12 | The amount different clusters of participants allocated to the anonymous receiver when acting as the dictator in the dictator game before the main experiment.** **a**, Experiment 1. **b**, Experiment 2. Each data point (gray circle) denotes one participant. The bottom, middle, and top lines of the box plot respectively represent the first quartile, the median, and the third quartile of the data. The lines extending beyond the box refer to 1.5 times the interquartile range (IQR), i.e., the distance between the third quartile (Q3) and the first quartile (Q1). ***: $p < 0.001$ after multi-comparison corrections. DG: dictator game. J: justice warriors. P: pragmatic helpers. R: rational moralists.

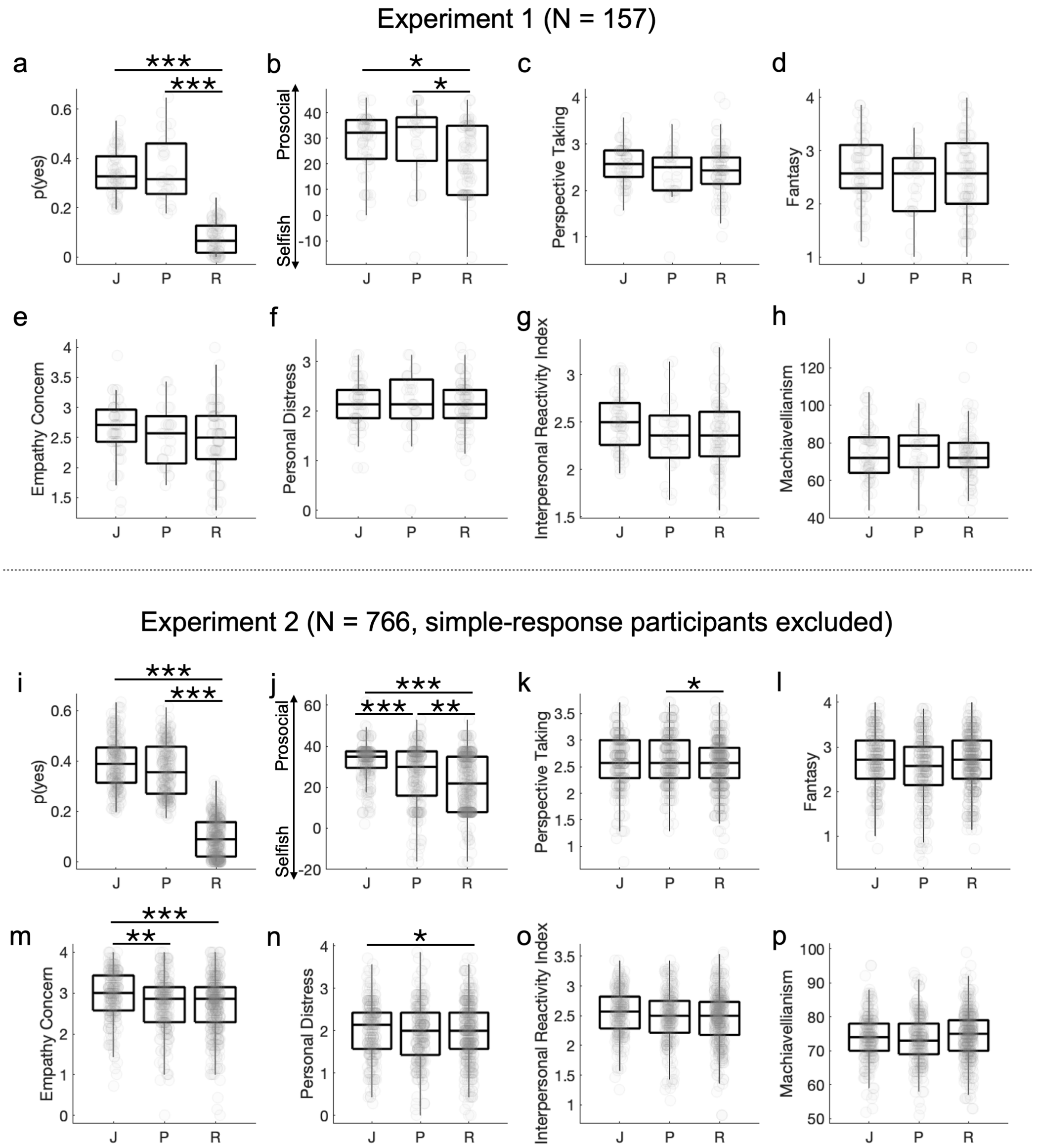


**Fig. S13 | Differences between justice warriors (J), pragmatic helpers (P), and rational moralists (R) in the probability of accepting the intervention offer and in personality measures.** For each panel, the bottom, middle, and top lines of the box plot respectively represent the first quartile, the median, and the third quartile of the data. The lines extending beyond the box refer to 1.5 times the interquartile range (IQR), i.e., the distance between the third quartile (Q3) and the first quartile (Q1). Each gray circle represents one participant. ***, ** and *: $p < 0.001$, $p < 0.01$ and $p < 0.05$ after multi-comparison corrections.

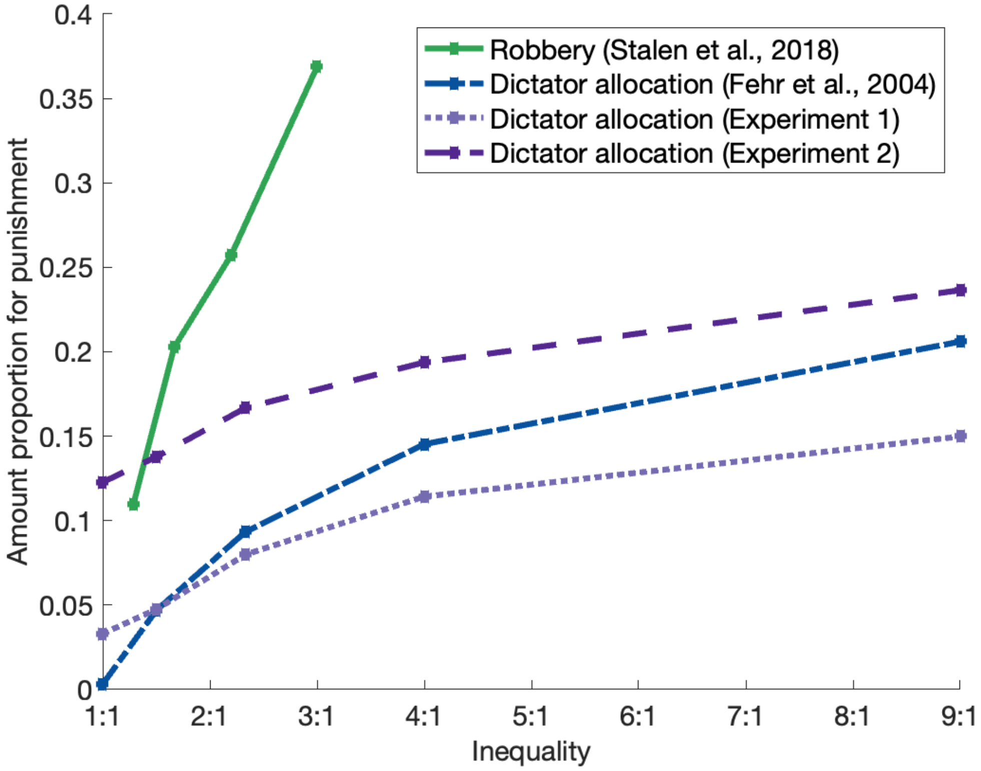


**Fig. S14 | The influence of cause of inequality as well as level of inequality on the proportion of amounts participants (as a third party) used to punish the transgressor.** The x-axis represents the ratio of the original allocation between the transgressor and the victim. A ratio of 1/1 means the amount of money is initially allocated equally between the transgressor and the victim, such as 50:50. A ratio of 9/1 means the transgressor has nine times the amount of money as the victim, such as 90:10. The y-axis represents the proportion of the amount that third parties (the participants) are willing to use for punishment relative to the maximum amount they have. For example, if a third party has up to 50 units and decides to use 10 units to punish the transgressor, this is recorded as 0.2, indicating that the third party is willing to use 20% of their available amount to punish the transgressor. All the plotted experiments are comparable in that the total amount that the transgressors can allocate between themselves and the victims is twice the amount held by the third parties (the participants). To make the experiments further comparable, only data from the punishment scenario and from the conditions with an impact ratio of 3 are plotted, that is, participants' spending of one unit reduces the transgressor's amount by three units. Among the four experiments, the initial inequality between the first and second parties in Fehr and Fischbacher (2004), like in our two experiments, came from a dictator game, where the transgressor (dictator) allocates a fixed amount of money between themselves and the victim (receiver). In contrast, the transgressor in Stallen et al. (2018) was framed as more malicious (or, more severe violation of social norms), who "robbed" a specific amount from the victim. Note that compared with the other three experiments, the punishment in Stallen et al. (2018), increases much faster with the level of inequality.

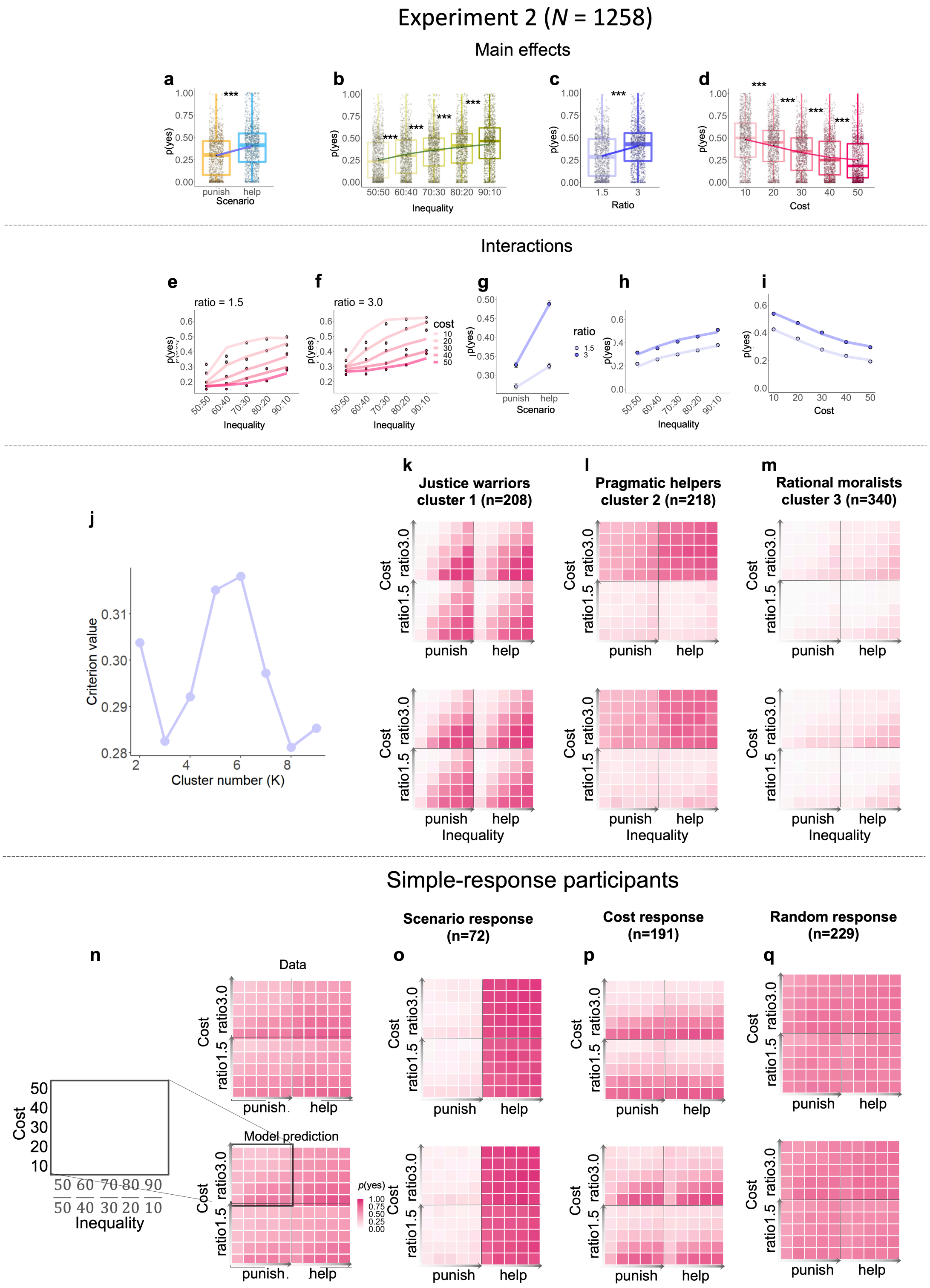


**Fig. S15 | All major findings of Experiment 1 were replicated in Experiment 2**. **a–d**, The main effects of scenario (a), inequality (b), ratio (c) and cost (d) on choice. **e–f**, The interaction of inequality × cost × ratio. **g**, The interaction of scenario × ratio. **h**, The interaction of inequality × ratio. **i**, The interaction of cost × ratio. **j**, Participants in Experiment 2 can be best classified as 6 clusters. **k–m**, The intervention patterns of the first three clusters of participants in Experiment 2: justice warriors, pragmatic helpers and rational moralists. **n–q**, Intervention probability of participants in the remaining 3 clusters. The participants'

intervention patterns of the newly observed three clusters in Experiment 2 were best fit by a simple-responsemodel (model 9). The x-axis of the heatmap is the severity of inequality from near equality (left, 50:50) to extreme inequality (right, 90:10). The y-axis of the heatmap is the cost of intervention from low cost (bottom, 10) to high cost (up, 50). The darker color on the heatmap represents a higher probability of intervention. The sub-maps on the upper left, upper right, bottom left and bottom right corners represent 4 sub-conditions.

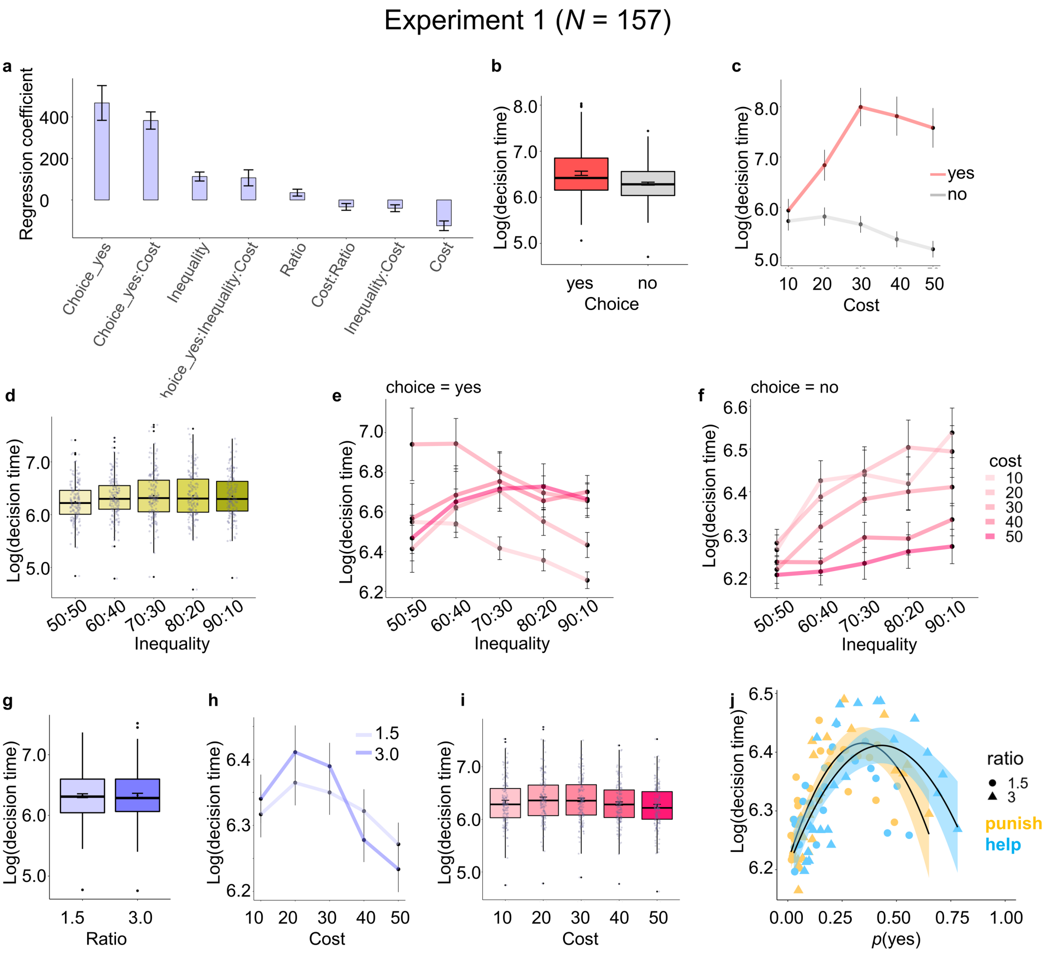


**Fig. S16 | Results of decision times in Experiment 1. a**, The variables that can significantly predict intervention decision time. A linear mixed-effect model was developed to assess the influence of all manipulated variables, participants' choice and their interactions on participants' decision time (see LMM1 and Table S4 for more details). Only the regression coefficients that were statistically significant are plotted. Figures **b - i** are further exhibitions of the effects shown in **a**. Decision time in y-axis (unit: milliseconds) was transformed into log-scale for readability. **b**, The main effect of choice on decision time. The average decision time when participants chose "yes" is longer than they chose "no". **c**, The interaction effect of cost × choice on decision time. In "yes" trials where participants decided to intervene, their decision time increases non-monotonically as intervention cost rises. While in "no" trials, participants' decision time monotonically decreases with rising cost. **d**, The main effect of inequality on decision time. Participants' decision time increases with the increasing inequality severity. **e - f**, The interaction effect of inequality × cost × choice on decision time. The interaction effect of inequality × cost is modulated by participants' choice. In "yes" trials, participants' decision time increases with inequality severity when the intervention cost is high, but decreases with inequality severity when the cost is low and moderately high. In "no" trials, participants' decision time consistently increases with the increasing inequality severity within each intervention cost condition and decreases overall as intervention cost rises. **g**, The main effect of ratio on decision time. Participants' overall decision time is longer in the high-impact ratio condition than in the low-impact ratio condition. **h**, The interaction effect of cost × ratio on decision time. Participants' decision time changes as an inverse-U shape as intervention cost rises. The high-impact ratio triggers a larger extent of amplitude change in the decision time. **i**, The main effect of cost on decision time. Participants' decision time decreases overall as intervention cost increases. **j**, The bell-shape relationship between participants' decision time (y-axis) and their intervention probability (x-axis) either in punishment (orange) or helping (cyan) scenario. Black curves denote the prediction of the multivariable nonlinear function. Shadings denote the SEM. For figures **a**, **c**, **e**, **f**, **h**, the bars or the dots represent mean value of measured variables across all participants, and the error bars denote SEM. For figures **b**, **d**, **g** and **i**, the bottom/top and middle lines of the box respectively indicate the first/third quartiles (Q1/Q3) and the median; the whiskers represent 1.5 times the IQR, which is the distance between the Q3 and the Q1. Data points beyond 1.5 times the IQR from the upper and lower quartiles are considered outliers and are represented by the filled points.

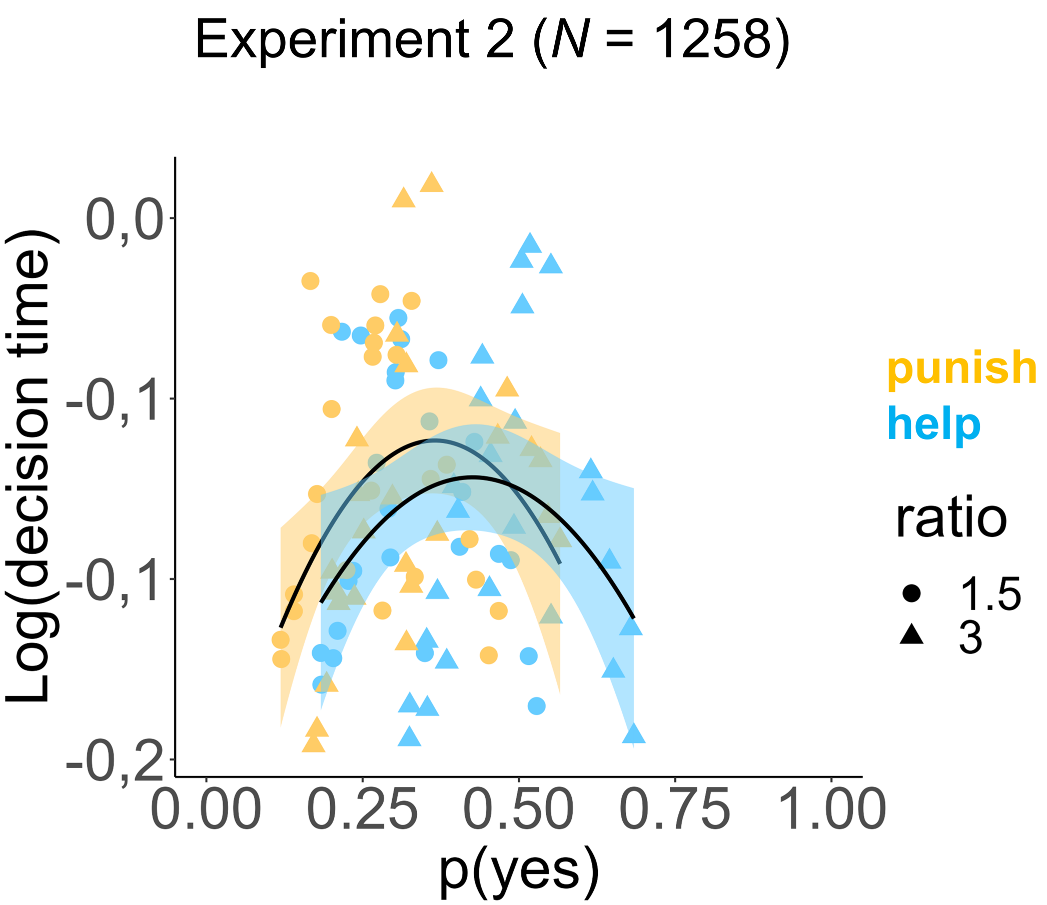


**Fig. S17 | The inverted-U shape between decision times and intervention probability in Experiment 2.** The bell-shape relationship between participants' decision time (y-axis) and their intervention probability (x-axis) either in punishment (orange) or helping (cyan) scenario. Black curves denote the prediction of the multivariable nonlinear function. Shadings denote the SEM.

### Experiment 2 (N = 766, simple-response participants excluded)

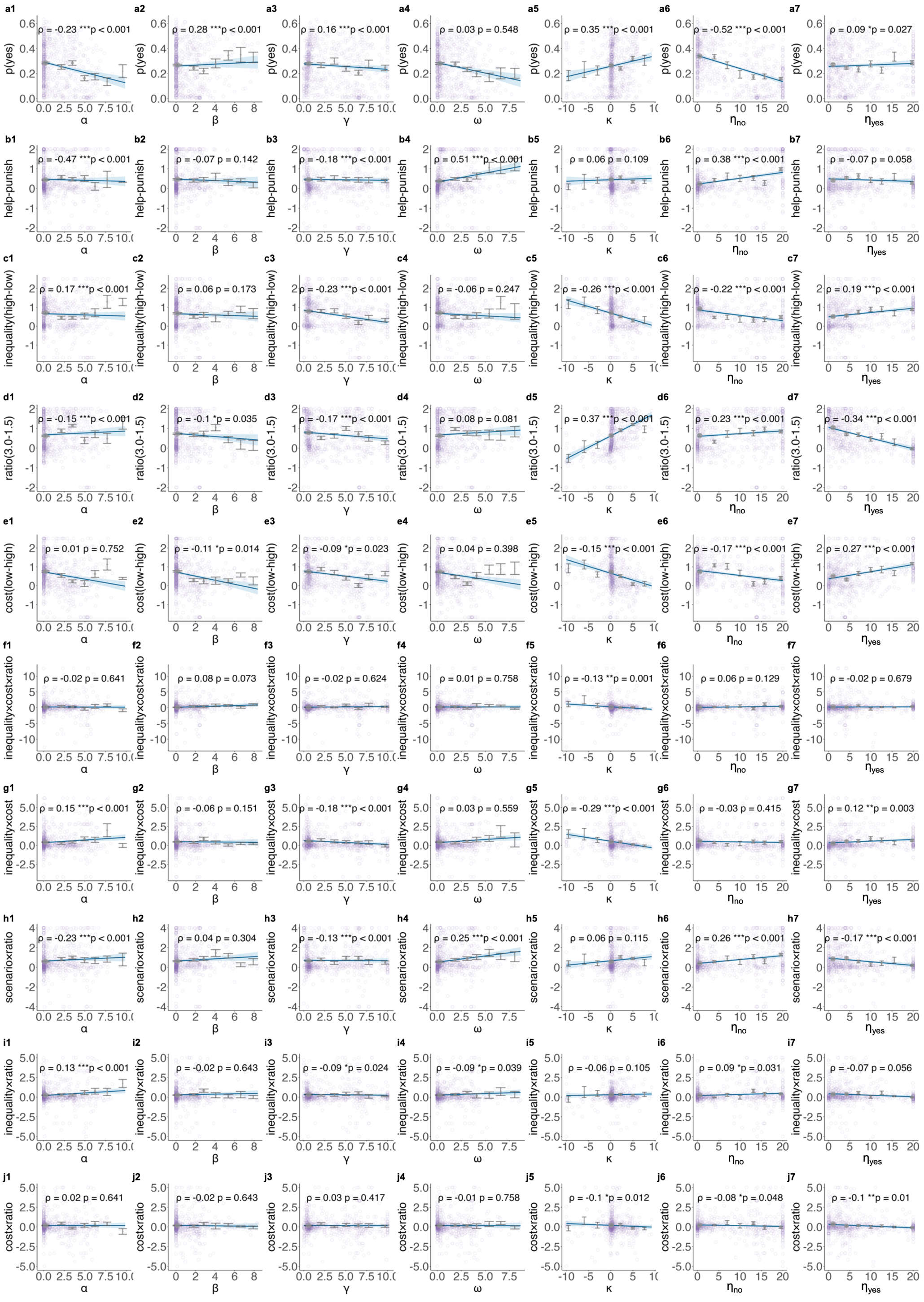


**Fig. S18 | The correlations between parameters estimated from the motive cocktail model and the model-free measurements for Experiment 2 (simple-response participants excluded)**. **a1 - a7**, The y-axis *p*(yes) represents the probability of intervention across all conditions for each participant. **b1 - b7**, The y-axis represents relative preference to help over punish, calculated as the probability of intervention in the helping scenario relative to that in the punishment scenario, normalized by the overall *p*(yes). **c1 - c7**, The y-axis represents the sensitivity to inequality, calculated as the probability of intervention in the high inequality trials (i.e., 70:30, 80:20, 90:10) relative to that in the low inequality trials (i.e., 50:50, 60:40), normalized by the overall *p*(yes). **d1 - d7**, The y-axis represents the sensitivity to ratio, calculated as the intervention probability difference between high impact ratio trials (ratio = 3.0) and low impact ratio trials (ratio = 1.5), normalized by the overall *p*(yes). **e1 - e7**, The y-axis represents the sensitivity to cost, calculated as the probability of intervention in low intervention cost trials (i.e., cost = 10, 20) minus that in high intervention cost trials (i.e., cost = 30, 40 and 50), normalized by the overall *p*(yes). **f1 - f7**, The y-axis represents the sensitivity to inequality under different levels of cost and ratio conditions, calculated as the normalized intervention probability difference in trials with different combinations of inequality, cost, and ratio: [(high ratio & high inequality & low cost - high ratio & high inequality & high cost)- (high ratio & low inequality & low cost - high ratio & low inequality & high cost)] minus [(low ratio & high inequality & low cost - low ratio & high inequality & high cost) - (low ratio & low inequality & low cost - low ratio & low inequality & high cost)]. **g1 - g7**, The y-axis represents the sensitivity to inequality under high-cost versus low-cost condition, calculated as the normalized intervention probability difference in trials with different combinations of inequality and cost: (high inequality & low cost – high inequality & high cost) - (low inequality & low cost – low inequality & high cost). **h1 - h7**, The y-axis represents the relative preference to help over punish under high versus low ratio conditions, calculated as the normalized intervention probability difference in trials with different combinations of scenario and ratio: (help & high ratio – help & low ratio) minus (punish & high ratio – punish & low ratio). **i1 - i7**, The y-axis represents the sensitivity to inequality under high versus low ratio conditions, calculated as the normalized intervention probability difference in trials with different combinations of inequality and ratio: (high inequality & high ratio – high inequality & low ratio) minus (low inequality & high ratio – low inequality & low ratio). **j1 - j7**, The y-axis represent the sensitivity to cost in high versus low ratio conditions, calculated as the normalized intervention probability difference in trials with different combination of cost and ratio: (low cost & high ratio – low cost & low ratio) minus (high cost & high ratio – high cost & low ratio). The x-axis for each column corresponds to one motive parameter of the motive cocktail model. Each panel illustrates the relationship between a motive parameter and a behavioral measure, with the x-axis divided into 8 bins across participants, and the y-axis displaying the mean (points) and standard deviation (error bars) within the corresponding bin. Each light-colored circle represents data from an individual participant. The blue line in each plot represents a linear regression between the original x and y coordinates (i.e., no bins), while the shaded area indicates the 95% confidence interval. The $\rho$ denotes partial correlation coefficient after controlling all other parameters. The *p* value was corrected for multiple comparisons using FDR.

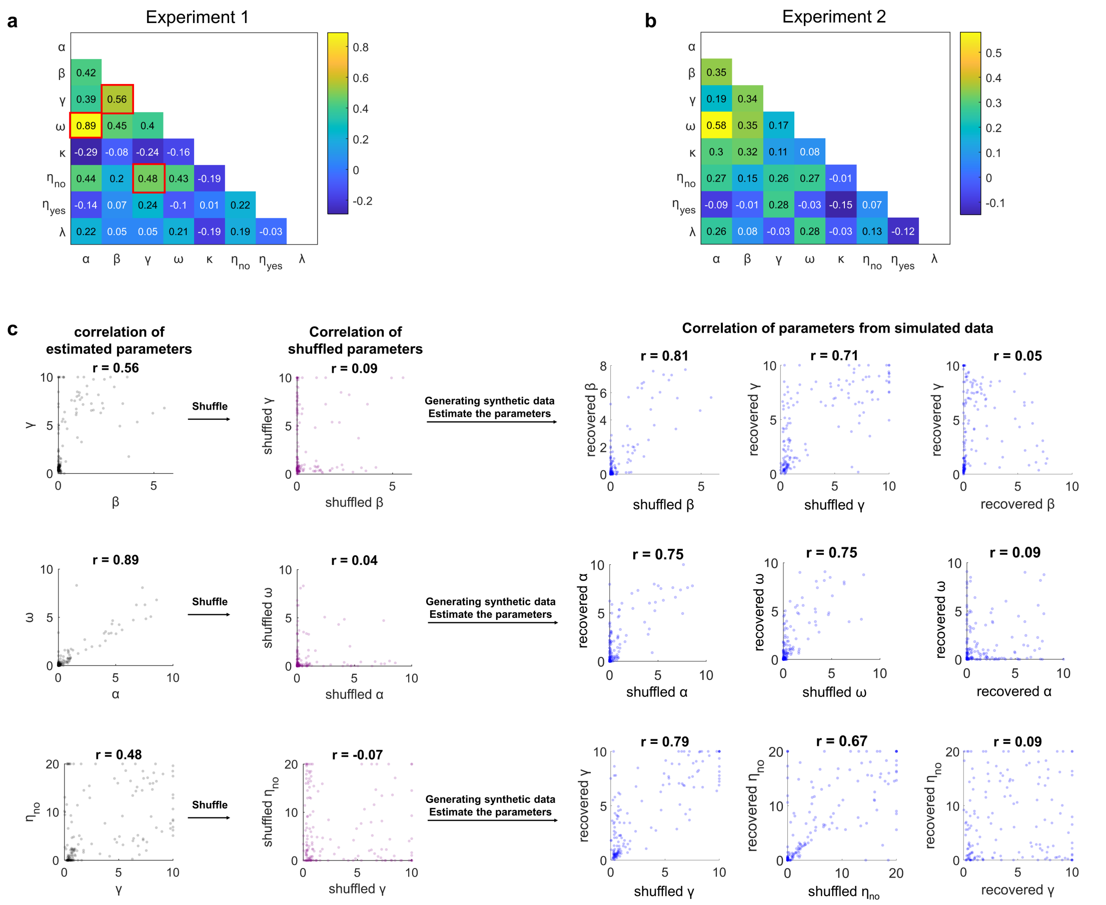


**Fig. S19 | Redundancy checks on the parameter space in the motive cocktail model**. **a–b**, Correlation matrix of parameters for Experiment 1 and Experiment 2. Colors code Pearson's *r*, where more yellow (blue) corresponds to a more positive (negative) correlation. Three high correlations (framed in red) between parameters  and ,  and ,  and  across all participants were examined. **c**, No evidence supports parameter redundancy. We used the following method to reject the possibility of parameter redundancy: we first shuffled the relationship between parameters across participants to decrease their correlation coefficient (Pearson's *r*, see column1 and column 2). The shuffled parameters were assigned to each participant randomly and used to generate synthetic intervention decisions, which were then used to fit the full motive cocktail model. If the high correlation pairs we observed in the real data were due to parameter redundancy, we would expect high correlations between the recovered parameters, although their correlations had been eliminated. In contrast, we observed the shuffled parameters are recoverable (columns 3 and 4) and more importantly, there is no correlation between the recovered parameters, in line with the shuffled pattern. These results suggest that the high correlations observed between  and , between  and , and between  and  reflected human participants' behavioral characteristics instead of redundancy in the model itself.

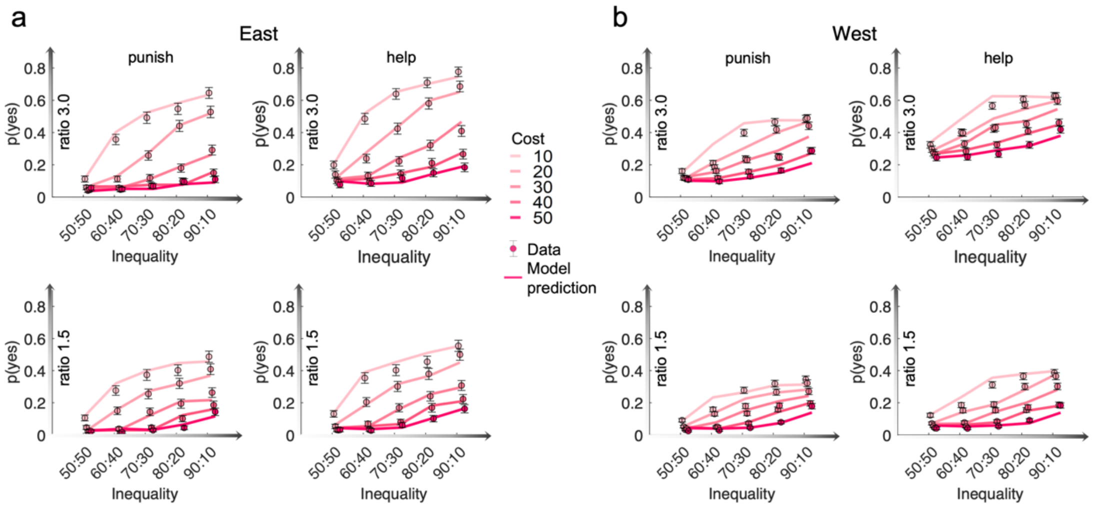


**Fig. S20 | Behavioral patterns comparison between East and West groups.** Figure **a-b** show data versus best-fitting model (model 8) predictions separately for East and West Groups. The probability of intervention, *p*(yes), is plotted against the inequality (from 50:50 to 90:10). Different colors code different levels of intervention cost (from 10 to 50, darker color for higher cost). Each sub-panel corresponds to one scenario and impact ratio condition. The dots and error bars respectively denote the mean and SEM across participants. The solid lines denote the predictions of the models.

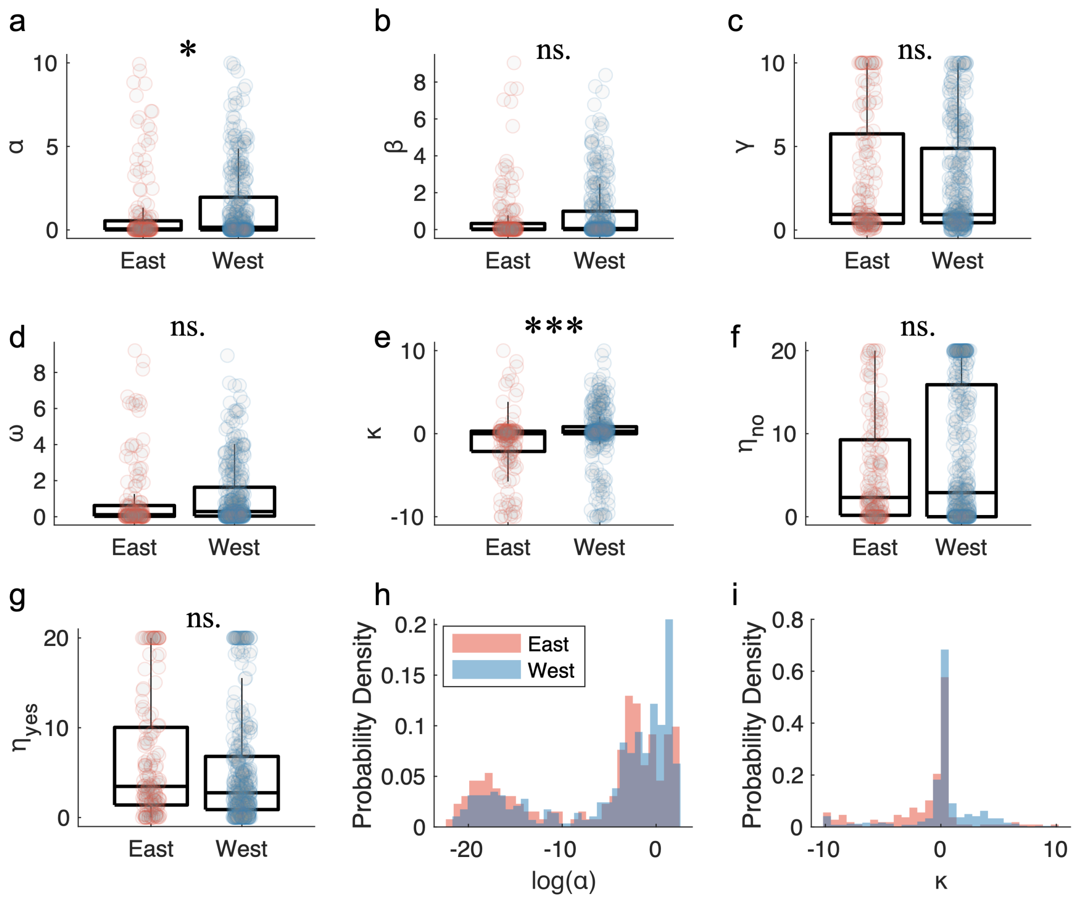


**Fig. S21 | Comparison of motive parameters between the East and West groups, with participants combined from Experiments 1 and 2**. **a–g,** Motives parameter comparison between the East and West groups for the parameters α, β, γ, ω, κ, $\eta_{no}$ and $\eta_{yes}$. The bottom, middle, and top lines of the box plot respectively represent the first quartile, the median, and the third quartile of the data. The lines extending beyond the box refer to 1.5 times the interquartile range (IQR), i.e., the distance between the third quartile (Q3) and the first quartile (Q1). Each light-colored circle represents the parameter value estimated from an individual participant. **h–i,** The distributions of parameter α and κ (the parameters with significant group differences) in each group. Red: East group. Blue: West group. *** and * respectively denote $p < 0.001$ and $p < 0.05$, with Bonferroni corrections for seven comparisons.
